# Localizable fluorescent metal ion indicators with tunable colors

**DOI:** 10.64898/2025.12.05.692539

**Authors:** Ming-Ming Wang, De-en Sun, Kai Johnsson

**Author notes:** M.W. and D.S. contributed equally to this work.

## Abstract

Elucidating the role of metal ion homeostasis in physiological and pathological processes requires detection tools with high sensitivity and selectivity, spectral versatility, and precise subcellular localization. Here, we introduce a modular strategy for generating fluorescent metal ion indicators which are comprised of a sulfonamide-functionalized chelator, a rhodamine derivative and a ligand for bioconjugation to self-labeling proteins. With a concise three-step synthesis, the design allows tuning of both the emission color and ligand specificity of the resulting indicators. Specifically, we developed a family of bright, color-tunable potassium indicators which can be selectively coupled to intra- or extracellular HaloTag and SNAP-tag fusion proteins. The increase in fluorescence upon binding to HaloTag or SNAP-tag fluorogenicity enabled wash-free live-cell imaging. The localized potassium indicators enabled the detection of dynamic potassium efflux in rat hippocampal neurons upon glutamate stimulation. Our work thus establishes a versatile platform for the generation of localizable fluorescent metal ion indicators and opens up new avenues for live-cell potassium sensing.

## Introduction

Metal ions are indispensable for diverse cellular processes, including osmotic regulation, metabolism, and signaling. Perturbations in their homeostasis, which disrupt the spatial distribution and concentration of intracellular ion pools, are tightly associated with aging and a wide spectrum of diseases^1^. Among these ions, potassium (K^+^) is both abundant and physiologically crucial, playing an essential role in maintaining the membrane potential that underlies proper cellular and neuronal signaling. In mammalian cells, this function relies on a pronounced transmembrane gradient, with intracellular concentrations of 140–150 mM compared to 3.5–5 mM in the extracellular milieu^2^. Consequently, the ability to monitor K^+^ dynamics spatiotemporally, both inside and outside cells, provides valuable insights into cellular physiology and disease mechanisms.

Fluorescent indicators provide a robust approach to monitor the spatiotemporal dynamics of metal ions in living systems^3,4^. Conceptually, they consist of two key components: a metal-chelating moiety and one or more fluorophores. Binding of the chelating group to a specific metal ion perturbs the electronic or molecular structure of the fluorophore(s), resulting in changes in fluorescence intensity or emission wavelength, which enabling dynamic metal sensing. For instance, a variety of synthetic K^+^ indicators^5^, including PBFI^6,7^, the TAC series^8–10^, KS2^11^, NK1^12^, RPS-1^13^ and others^14,15^, have been developed by attaching crown ether- or cryptand-based chelators to fluorophores. However, these indicators often lack cell-type or subcellular specificity and exhibit poor membrane permeability, limiting their applicability for precise imaging in complex biological systems. Genetically encoded potassium indicators have been developed by fusing fluorescent proteins (FP) with K^+^-binding proteins, including GEPII^16^, KIRIN1s^17^, GINKOs^17,18^ and KRaIONs^19^. While these FP-based indicators can be genetically targeted to specific cellular populations and subcellular compartments, they generally possess lower brightness and a narrow spectral range, being largely restricted to the green fluorescence channel compared with synthetic indicators. The recently developed RGEPOs^20^, based on red fluorescent protein mApple provide a partial improvement over these constraints.

Chemigenetic fluorescent indicators integrate the brightness and spectral versatility of synthetic fluorophores with the biocompatibility and genetic precision of FP-based indicators, enabled by self-labeling proteins such as SNAP-tag and HaloTag^21–24^. Building on this strategy, several chemigenetic K^+^ indicators have recently been reported, each with distinct advantages and limitations. The FAST-based green-emitting K^+^ indicator, K^+^-FAST-5.1^25^, has an apparent dissociation constant (*K*_*D*_) of 2 mM for K^+^ *in vitro*, which is too high for intracellular imaging and has not yet been validated in living cells. The HaloTag-based green-emitting indicator TLSHalo^26^ can detect extracellular K^+^ changes, but is not suitable for intracellular applications and suffers from photobleaching. More recently, the HaloKbp1^27^ series was developed by inserting a K^+^-binding protein into HaloTag7 and labeling it with environmentally sensitive rhodamine derivatives. These indicators show high brightness in the red-to-far-red region, large responses, and tunable *K*_*D*_ values, but are currently restricted to intracellular K^+^ detection.

Here, we present a series of specific and sensitive fluorescent indicators for various metal ions which can be coupled to HaloTag7 and SNAP-tag2 for subcellular localization. Among them, we highlight the family of indicators for K^+^, which offers distinct color variants extending into the farred spectrum, high brightness, wash-free imaging capability, and enables both dynamic intracellular and extracellular K^+^ sensing in live cells.

## Results and Discussion

### Design and synthesis of localizable fluorescent indicators for various metal ions

We recently introduced the so-called MaP dyes, in which the lactone-forming carboxylic acid of rhodamines is replaced with an amide bearing an electron-withdrawing group on the nitrogen (e.g., sulfonyl or cyano substituents)^28,29^. This modification favors the non-fluorescent spirolactam form when the dye is free in solution, yet promotes conversion to the open, fluorescent state upon binding to biomolecules, including HaloTag, thereby enabling wash-free imaging with low background. Leveraging this scaffold, we subsequently developed MaPCa dyes^30^, a family of cell-permeable calcium indicators designed to couple with HaloTag7 for wash-free and targetable calcium sensing in live cells. By placing the calcium chelator in close proximity to the rhodamine core, the design enables efficient quenching of rhodamine fluorescence through photoinduced electron transfer (PET) from the free chelator^31^.

We envisioned that this design concept could be generalized by attaching suitable metal ion chelators to MaP dyes, creating a versatile platform for the development of fluorogenic metal ion indicators (Figure 1a). A key synthetic challenge, however, was the lack of a mild and efficient method to install sulfonamides onto metal ion chelators. Classical sulfonamide synthesis from aromatic precursors typically involves ClSO_3_H followed by ammonolysis, leading to tedious protection/deprotection steps for calcium chelators and extremely low yields for potassium chelators under such harsh conditions (Supplementary Figure 1).

**Fig. 1.**
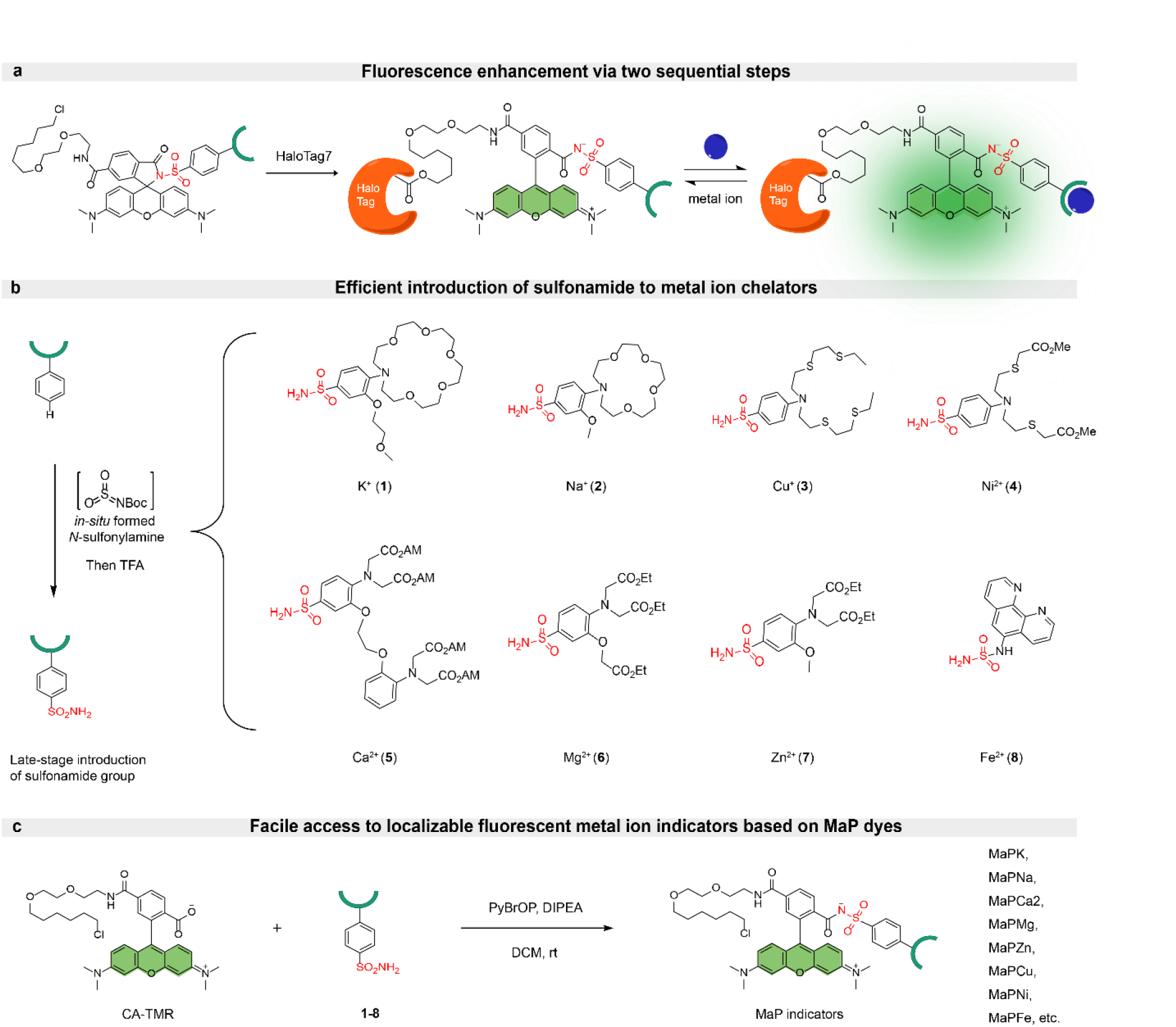
Schematic of the general design strategy for MaP dye-based metal ion indicators. (**a**) Schematic of the double-turn-on mechanism of MaP dye-based indicators. In the unbound state, the indicators exist in a colorless, spirocyclic form. HaloTag7 binding induces ring opening to the zwitterionic state, becoming potentially fluorescent but remaining PET-quenched by the metal ion-binding moiety. Subsequent metal ion binding suppresses PET, resulting in full fluorescence emission. (**b**) Late-stage introduction of sulfonamide to metal ion chelators. (**c**) A general synthetic route to access MaP dye-based indicators for various metal ions.

To overcome this bottleneck, we applied our recently developed synthetic strategy for accessing arylsulfonamides through *in-situ* formed *N*-sulfonylamine^32^, which enabled facile introduction of sulfonamide functional group onto various metal ion chelators (**1**-**7**) for potassium, sodium, calcium, magnesium, zinc, copper(I) and nickel. We also synthesized the iron(II) chelator **8** with a sulfamide functionality under similar conditions, employing 5-amino-1,10-phenanthroline as the precursor (Figure 1b and Supplementary Figure 1). With sulfonamides **1**-**7** and sulfamide **8**, we constructed a MaP558 series of localizable metal ion indicators by incorporating the corresponding sulfonamide into tetramethylrhodamine bearing a chloroalkane ligand (CA-TMR). These indicators were designated with postfixes indicating their absorption maxima in nanometers (i.e., TMR 558). Screening of various coupling reagents identified bromo-tris-pyrrolidino-phosphonium hexafluorophosphate (PyBrOP) as an efficient mediator for the synthesis of MaP dye-based indicators. When combined with HaloTag7, the resulting chemigenetic indicators provide targetable sensing capability. In total, we synthesized eight distinct indicators: MaPK-558, MaPNa-558, MaPCa2-558, MaPMg-558, MaPZn-558, MaPCu-558, MaPFe-558 and MaPNi-558, demonstrating the versatility and general applicability of the MaP dye-based indicator platform (Figure 1c and Supplementary Figure 2).

### *In vitro* characterization of localizable MaP558-based indicators

We then characterized these fluorescent indicators *in vitro* in the presence and absence of HaloTag7 by measuring fluorescence intensities across varying concentrations of the corresponding metal ions (Supplementary Table 1-2). Indicators with ester protecting groups were first saponified with potassium hydroxide (KOH) and purified via high-performance liquid chromatography (HPLC), providing the free acids that mimic their deprotected form in cells. Notably, most synthesized indicators showed a pronounced response to their target ions. MaPNa-558 and MaPK-558 employ aza-crown ethers as chelators (Figure 2a and Figure 3b), with variations in ring size and lariat length that have been reported to maintain strong Na^+^/K^+^ selectivity despite their binding competition^15,33^. MaPNa-558 exhibited 5.1-fold and 5.0-fold fluorescence increases upon HaloTag7 and sodium binding, respectively (*K*_*D*_ = 205 mM; Supplementary Table 1). MaPNa-558 showed excellent Na^+^/K^+^ selectivity, with a ΔF/F_0_ response of 262% over a Na^+^ concentration range of 0–150 mM, and a selectivity ratio of 11 relative to K^+^ (Figure 2a and Supplementary Figure 3a). For MaPK-558, an aza-crown ether was chosen over a cryptand as the chelating motif because its moderate affinity for K^+^ allows sensitive monitoring of potassium levels in both extracellular (3.5–5 mM) and intracellular (140– 150 mM) environments. MaPK-558 showed a 4.2-fold fluorescence turn-on upon binding to HaloTag7 and an additional 3.9-fold increase upon potassium binding (*K*_*D*_ = 33.8 mM). Moreover, MaPK-558 also demonstrated high K^+^/Na^+^ selectivity, yielding a ΔF/F_0_ of 287% across 0–150 mM K^+^, and selectivity ratio of 12.5 over Na^+^ (Figure 3b, g). Compared with the synthesis of the previously published MaPCa-558^30^, the synthesis of MaPCa2-558 was greatly simplified by starting from commercial BAPTA-AM without a methyl group. MaPCa2-558 has a reduced calcium affinity (*K*_*D*_ = 1.8 μM, versus 0.41 μM for MaPCa-558; Figure 2b), thereby offering a complementary measurement range to that of the MaPCa-558 series. MaPMg-558 and MaPZn-558, both employing carboxylic acids as chelators^34–36^, exhibited robust responses to magnesium and zinc (*K*_*D*_ = 3.4 mM and 4.6 μM, respectively; Figure 2c, d), but only weak HaloTag7-induced fluorescence turn-on, as also observed for MaPCa-558. This behavior presumably arises from the negatively charged carboxylic groups in MaPCa-558, MaPZn-558 and MaPMg-558, which may push the spirocyclization equilibrium of TMR to the open form, thereby attenuating its sensitivity to HaloTag7 binding. In particular, binding of MaPMg-558 to HaloTag7, as verified by high-resolution mass spectrometry (HRMS) (Supplementary Figure 4), did not at all effect its spectroscopic properties. MaPCu-558, with a thioether-rich receptor^37^, exhibited strong HaloTag7-induced fluorescence enhancement (8-fold) and high-affinity Cu^+^ binding (*K*_*D*_ = 267 pM; Figure 2e). However, MaPNi-558, bearing thioether and carboxylic acid chelating groups^38^, failed to produce the expected titration response in Ni^2+^ buffer and was therefore not further pursued. Whereas the aforementioned indicators share a common PET-based turn-on mechanism, MaPFe-558 was constructed by incorporating a 1,10-phenanthroline chelator into CA-TMR via the sulfamide bridge and was anticipated to behave as a turn-off probe owing to paramagnetic quenching by ferrous ion^39^. Consistent with this prediction, addition of Fe^2+^ decreased fluorescence intensity, with a measured *K*_*D*_ of 0.46 μM (Figure 2f). However, partial degradation of MaPFe-558 was observed even at −20°C, likely due to the intrinsic instability of sulfamide linkage (Supplementary Figure 5), which limits its practical applicability.

**Fig. 2.**
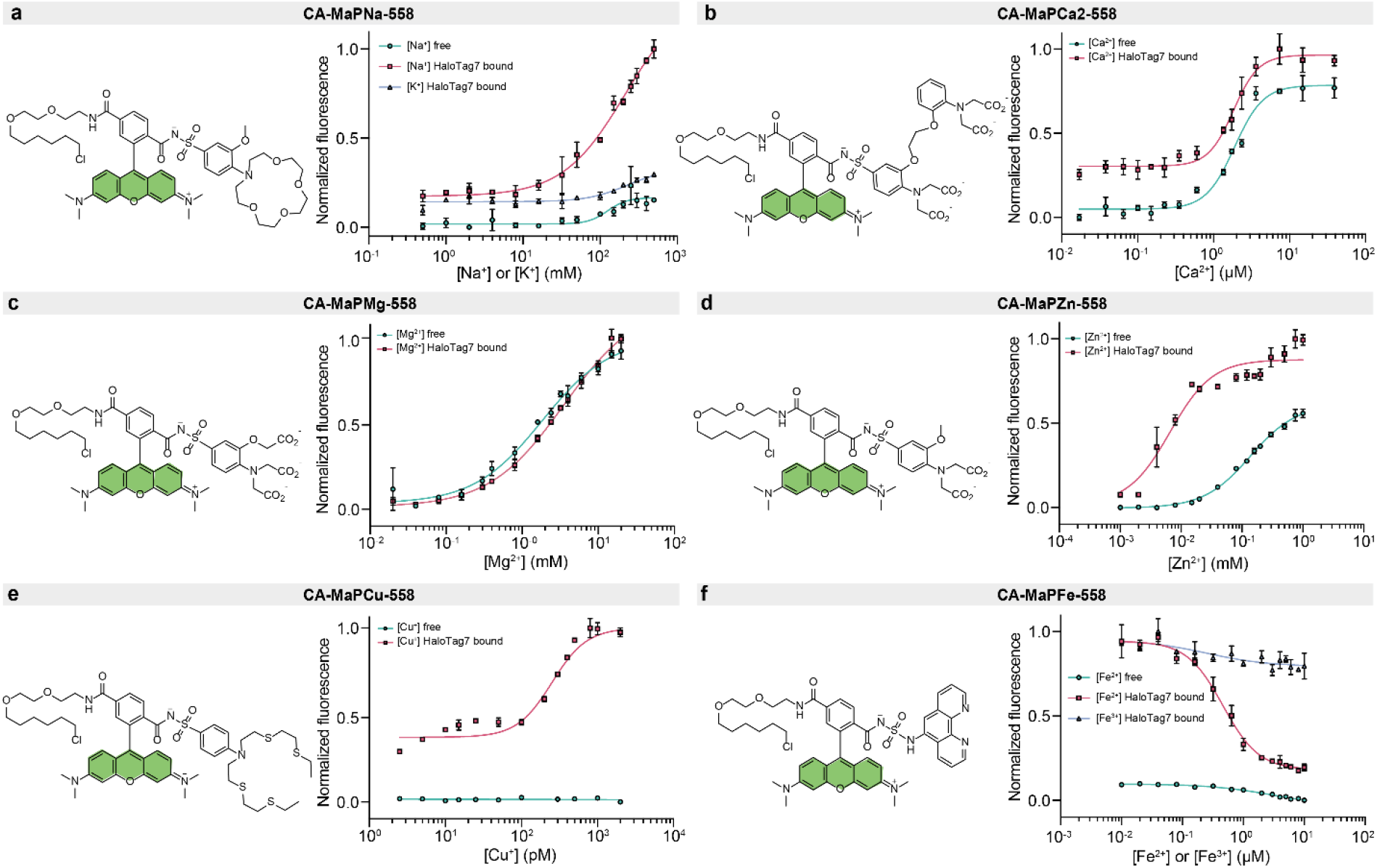
*In-vitro* characterization of MaP dye-based indicators for various metal ions. (**a-f**) metal ion titration curves of MaPNa-558 (**a**), MaPCa2-558 (**b**), MaPMg-558 (**c**), MaPZn-558 (**d**), MaPCu-558 (**e**) and MaPFe-558 (**f**), all derived from CA-TMR. Error bars represent the mean ± standard deviation from three technical replicates. Curves were fitted with sigmoidal function and *K*_*D*_ values are provided in Supplementary Table 1 and 2.

**Fig. 3.**
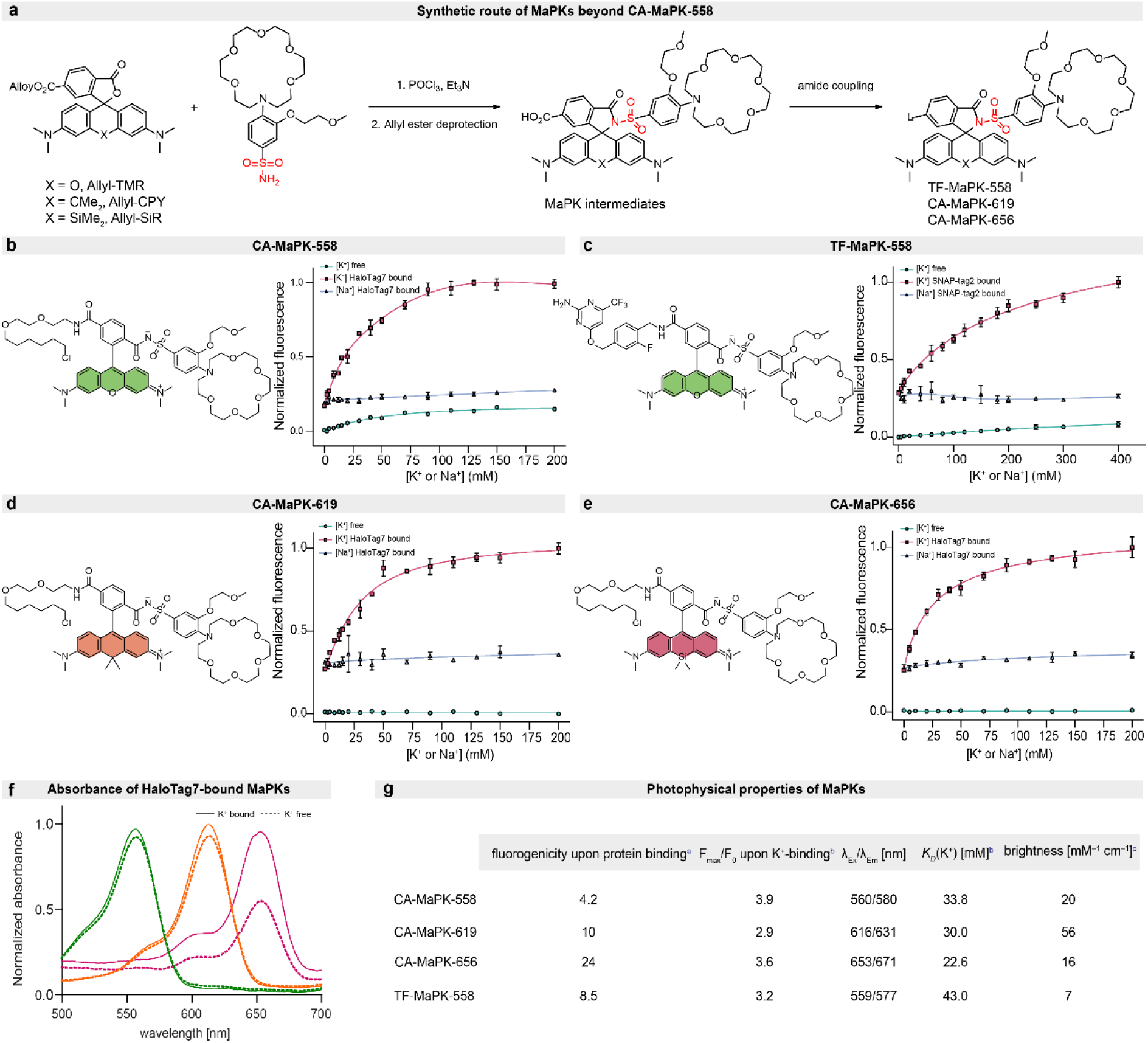
*In-vitro* characterization of MaPK indicators. (**a**) Schematic illustration of the synthetic route for MaPK indicators with different colors and ligands. (**b**-**e**) Potassium and sodium titration curves of MaPK indicators. Error bars represent the mean ± standard deviation from three technical replicates. (**f**) Absorbance spectra of HaloTag7-bound MaPK indicators showing potassium-dependent increase in absorbance for CA-MaPK-656, whereas CA-MaPK-558 and CA-MaPK-619 exhibit minimal changes. Green: CA-MaPK-558, Orange: CA-MaPK-619, Magenta: CA-MaPK-656. (**g**) Photophysical properties of MaPK indicators. ^a^Fluorescence increase at saturating potassium concentration. ^b^In HaloTag7-bound or SNAP-tag2-bound state. ^c^At saturating potassium concentration and HaloTag7-bound or SNAP-tag2-bound.

With the aforementioned particular interest in measuring cellular K^+^ fluctuations in mind, we then expanded our MaPK toolbox by generating MaPK-619 and MaPK-656 through incorporation of sulfonamide **1** into CA-CPY (CA derivative of carbopyronine; absorption maxima 619 nm) and CA-SiR (CA derivative of silicon rhodamine; absorption maxima 656 nm). In addition to HaloTag7 labeling, we further developed MaPK-558 with trifluoromethyl fluorobenzyl pyrimidine (TF) ligand for covalent labeling with SNAP-tag2. These three variants were prepared in a three-step sequence starting from allyl-TMR, allyl-CPY or allyl-SiR. The spirocyclic carboxylic group was first converted to the corresponding acyl chloride and coupled with sulfonamide **1**. Subsequent palladium-catalyzed deprotection and amide coupling afforded the final TF- or CA-conjugated products (Figure 3a). *In-vitro* titration revealed that TF-MaPK-558, CA-MaPK-619 and CA-MaPK-656 all exhibited strong fluorescence enhancement upon conjugation with SNAP-tag2 or HaloTag7 (8.5-24 fold) and significant turn-on upon K^+^ binding (2.9-3.6 fold) (Figure 3c-e, g). The absorbance spectra of CA-MaPK-558 and CA-MaPK-619 showed minimal changes upon K^+^ addition, suggesting that K^+^ binding does not affect the spirocyclization equilibrium and supporting our hypothesis of a PET quenching mechanism. However, for CA-MaPK-656, alterations in the spirocyclization–ring-opening equilibrium may also contribute to the fluorescence enhancement observed under high K^+^ conditions (Figure 3f). All MaPK variants showed high selectivity for K^+^ over Na^+^, with *K*_*D*_ values in the range of 22.6-43.0 mM (Figure 3g). In a similar manner, TF-MaPNa-558, CA-MaPNa-619 and CA-MaPNa-656 were synthesized and demonstrated robust turn-on response and sodium selectivity (Supplementary Figure 3 b-d and Supplementary Table 1). In comparison to the HaloTag7 conjugates (CA-MaPK-558 and CA-MaPNa-558), the SNAP-tag2 variants (TF-MaPK-558 and TF-MaPNa-558) exhibited lower extinction coefficients and quantum yields, resulting in reduced overall brightness (Figure 3g and Supplementary Table 1). This decrease likely reflects differences in the local environment provided by the SNAP-tag2 binding pocket and interface, which may affect the photophysical properties of MaP dyes.

### Extracellular and intracellular applications of localizable MaPKs in live cells

For intracellular K^+^ imaging, MaPK indicators were applied to co-cultures of U2OS cells stably expressing a nuclear-localized HaloTag7-SNAP-tag2 or not. Imaging after 2 h of incubation under no-wash conditions revealed efficient HaloTag7-SNAP-tag2 labeling (Figure 4a), confirming that the indicators are cell permeable. Moreover, comparing the cytosolic background fluorescence in non-expressing U2OS cells with the nuclear signal in expressing cells showed that all MaPKs exhibit excellent signal-to-background ratios. CA-MaPK-558, CA-MaPK-619, CA-MaPK-656 and TF-MaPK-558 displayed F_nuclear_/F_cytoplasm_ values of 2.1, 4.6, 10.5 and 2.9, respectively. This high contrast can be attributed to the strong fluorogenicity of these substrates, as observed *in vitro*. MaPKs possess high selectivity against Na^+^, which is crucial for extracellular K^+^ sensing due to the more than 30-fold excess of extracellular Na^+^. To investigate whether MaPKs can report extracellular K^+^ dynamics in live cells, we used U2OS cells displaying HaloTag7 on the plasma membrane by fusing it to an N-terminal Igκ signal peptide and a C-terminal PDGFR transmembrane domain. Incubation with 1 μM MaPKs for 2 h yielded efficient HaloTag7 labeling with clear membrane localization (Figure 4b). Different KCl concentrations were applied to the culture medium, with a basal KCl concentration of 5.33 mM, and all three indicators detected K^+^ changes in extracellular environments. CA-MaPK-558 showed a *K*_*D*_ value of 34 mM, consistent with measurements using purified HaloTag7, whereas reliable fitting could not be obtained for CA-MaPK-619 and CA-MaPK-656. The maximal K^+^-dependent fluorescence responses were 84%, 31%, and 21% for CA-MaPK-558, CA-MaPK-619, and CA-MaPK-656, respectively (Figure 4d and Supplementary Figure 6). We next assessed intracellular K^+^ responsiveness by applying MaPKs to U2OS cells stably expressing cytoplasmic HaloTag7–SNAP-tag2. Labeling was evenly distributed throughout the cytoplasm (Figure 4c), with different indicators showing distinct labeling kinetics: CA-MaPK-619 and CA-MaPK-656 showed complete labeling within around 1 h, whereas CA-MaPK-558 required over 2 h. These differences might be due to differences in permeability and/or the known differences in labeling kinetics of their dye backbones TMR, CPY, and SiR to HaloTag7^40^. Notably, MaPK-558 with a TF ligand labeled faster than with a CA ligand (Supplementary Figure 7), likely due to improved dye permeability^24^. To evaluate intracellular performance, cells stably expressing cytoplasmic HaloTag7–SNAP-tag2 were permeabilized with digitonin to deplete cytosolic K^+^ and then exposed to defined K^+^ concentrations. Under these conditions, TF-MaPK-558, CA-MaPK-558, CA-MaPK-619, and CA-MaPK-656 exhibited ΔF/F_0_ values of 89%, 72%, 35% and 40%, respectively (Figure 4e and Supplementary Figure 6).

**Fig. 4.**
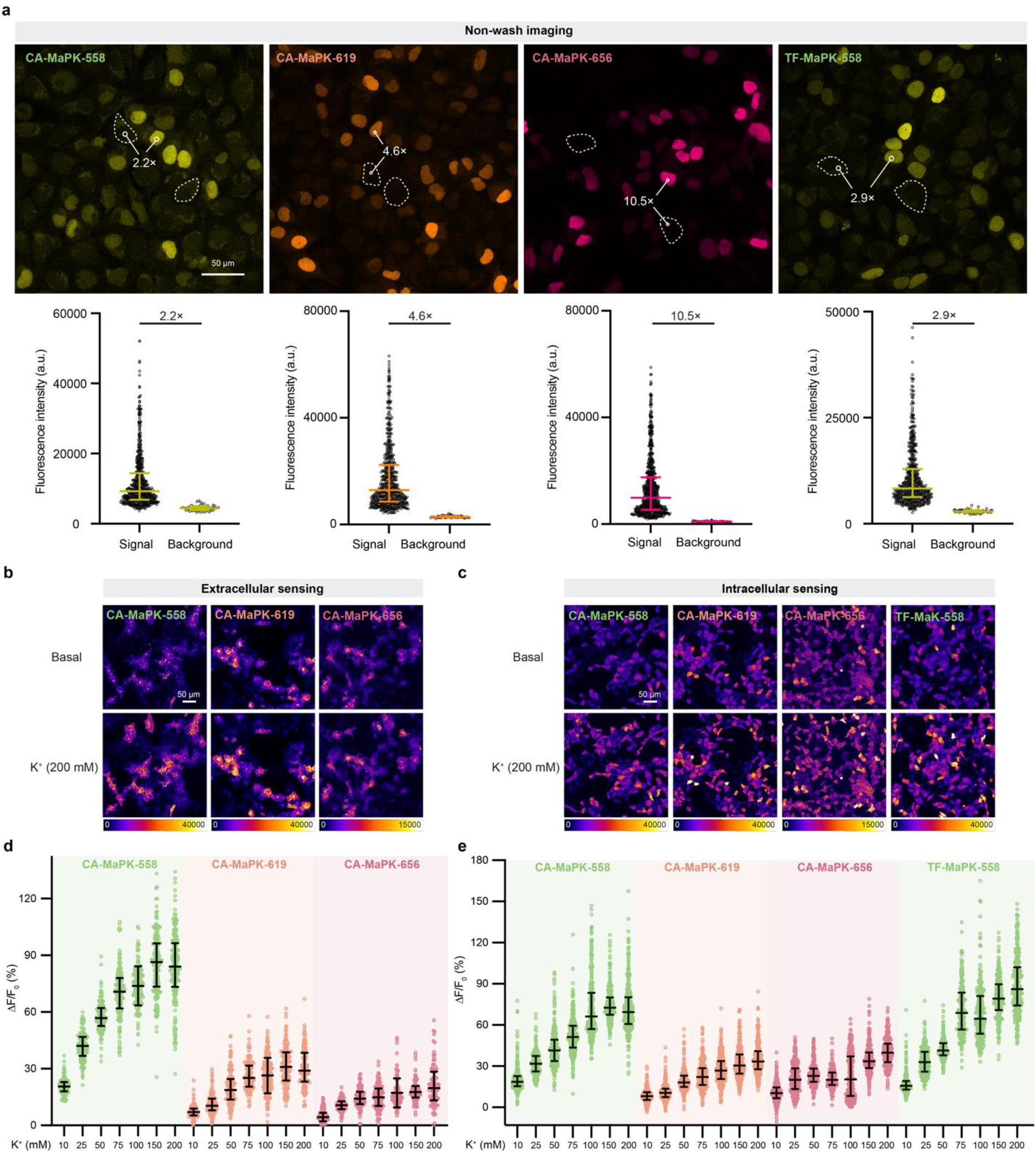
Extracellular and intracellular characterization of MaPKs. (**a**) Fluorescence microscopy images of a co-culture of HaloTag7–P30–SNAP-tag2–NLS–P2A–NLS–mTurquoise2-expressing and non-expressing U2OS cells. After incubation with 1 μM MaPKs for 2 h, cells were imaged under no-wash conditions. Turn-on values, reported as signal-to-background ratios with associated cell numbers were: CA-MaPK-558 (585/54), CA-MaPK-619 (582/54), CA-MaPK-656 (704/51) and TF-MaPK-558 (574/53). (**b**) Fluorescence microscopy images of U2OS cells stably expressing Igκ-HaloTag7-PDGFR, incubated with 1 μM CA-MaPKs and imaged before and after the addition of 200 mM KCl. (**c**) Fluorescence images of U2OS cells stably expressing HaloTag7–P30–SNAP-tag2–P2A–NLS–mTurquoise2, incubated with 1 μM-MaPKs and imaged before and after the addition of 200 mM KCl. Cells were permeabilized with 4 μM digitonin for 10 min prior to imaging. (**d**) Extracellular K^+^ titration dose−response curves of MaPKs as described in (**b**). The *K*_*D*_ value of CA-MaPK-558 was estimated to be ∼34 mM by fitting the titration curve with a sigmoidal function. n = 101, 135, 137, 144, 163, 159 and 187 cells for CA-MaPK-558 group; n = 135, 182, 136, 133, 184, 147 and 142 cells for CA-MaPK-619 group; n = 118, 109, 112, 133, 106, 107 and 99 cells for CA-MaPK-656 group. (**e**) Intracellular K^+^ titration dose−response curve of MaPKs as described in (**c**). n = 317, 284, 340, 389, 274, 384 and 393 cells for CA-MaPK-558 group; n = 274, 321, 292, 298, 354, 337 and 302 cells for CA-MaPK-619 group; n = 401, 339, 271, 263, 352, 295 and 228 cells for CA-MaPK-656 group; n = 263, 242, 263, 212, 229, 272 and 323 cells for TF-MaPK-558 group. Error bars represent the median with interquartile range (**a, d, e**). Data are pooled from one (**a**) two (**d**) or three (**e**) independent experiments. Scale bars: 50 μm.

### Localizable MaPKs report on potassium dynamics in neurons

Potassium is essential for maintaining neuronal physiology. Elevated extracellular K^+^, for instance, drives membrane depolarization and promotes neuronal hyperexcitability, and disruptions in K^+^ homeostasis are strongly linked to neurological disorders such as epilepsy^41,42^. To enable extracellular K^+^ sensing, HaloTag7 was displayed on the plasma membrane in rat primary hippocampal neurons via rAAV-mediated expression, by fusing with an N-terminal Igκ signal peptide and a C-terminal PDGFR transmembrane domain. HaloTag7-expressing neurons were then individually labeled with CA-MaPK-558, CA-MaPK-619, or CA-MaPK-656, consistently achieving efficient labeling and uniform membrane-localized fluorescence (Figure 5a). Upon stimulation with 30 mM or 200 mM K^+^, labeled neurons exhibited robust fluorescence increases of approximately 55% and 107% for CA-MaPK-558, 15% and 46% for CA-MaPK-619, while 21% and 38% for CA-MaPK-656, respectively (Figure 5b), showing a trend consistent with the extracellular titration results in U2OS cells. Previous studies have shown that glutamate-induced overactivation of its receptors triggers a substantial potassium efflux from neurons^43,44^. To assess whether these intracellular K^+^ changes could be monitored using MaPKs, we performed glutamate stimulation experiments in primary neuronal cultures expressing cytosolically localized HaloTag7-EGFP. As wash-free labeling helps to preserve cell viability^45^, neurons were labeled with CA-MaPK-558, and imaged under no-wash conditions. CA-MaPK-558 labeled both soma and dendrites uniformly (Figure 5c). Upon addition of 500 μM glutamate, the fluorescence ratio (MaPK-558/EGFP) dropped sharply by approximately 30% within 90 s (Figure 5d), indicating rapid potassium efflux. These results suggest that MaPKs are well-suited for monitoring transient intracellular and extracellular potassium dynamics in cultured primary hippocampal neurons.

**Fig. 5.**
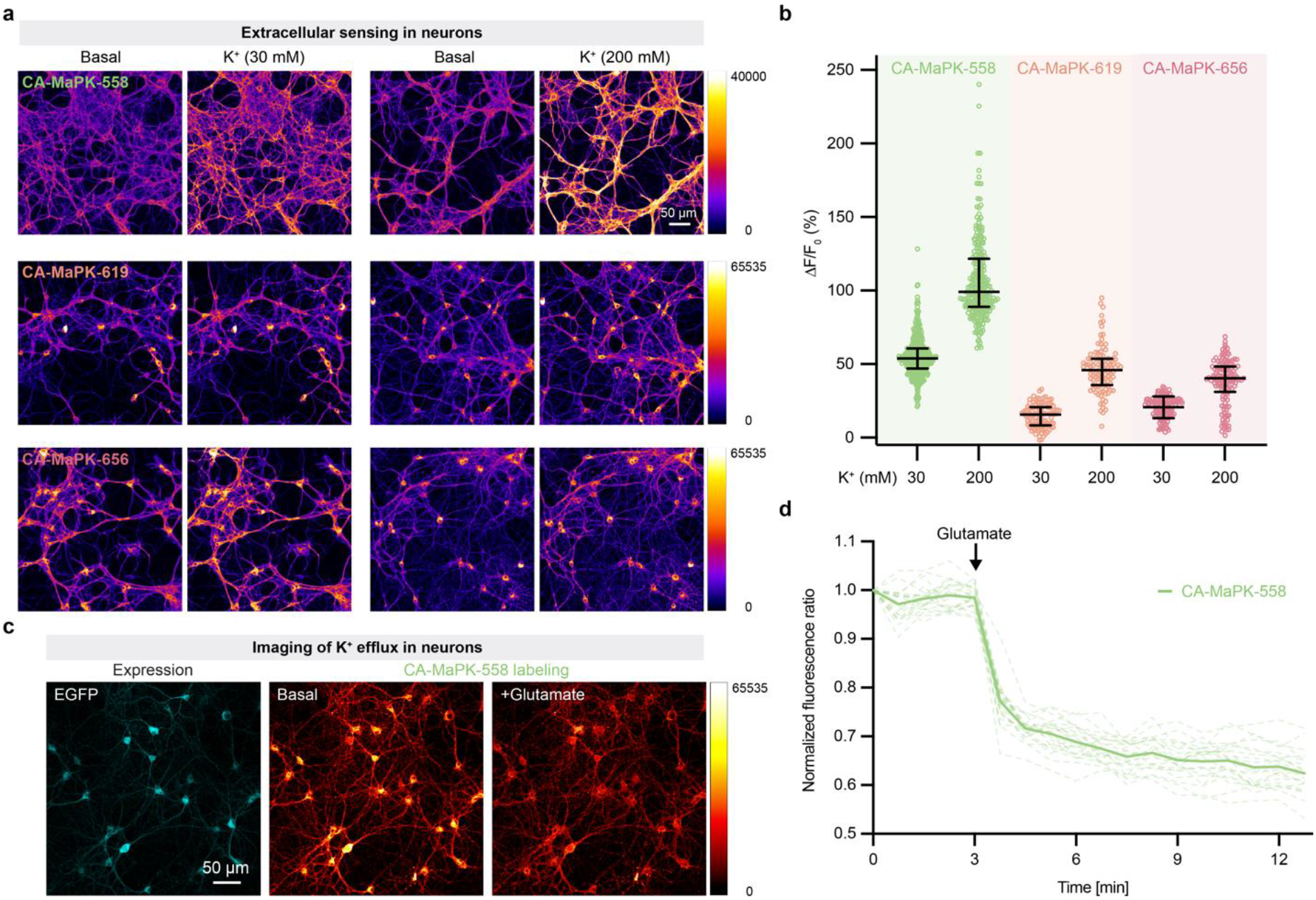
MaPKs can monitor potassium concentration changes in neurons. (**a**) Fluorescence images of primary rat hippocampal neurons expressing Igκ-HaloTag7-PDGFR, incubated with 1 μM CA-MaPKs and imaged before and after adding KCl (30 mM or 200 mM); MaPKs fluorescence (fire). (**b**) Quantification ΔF/F_0_ of MaPKs in response to 30 mM or 200 mM KCl as described in (**a**). Error bars represent the median with interquartile range; n = 406, 251, 100, 105, 126 and 133 neurons, pooled from three independent experiments. (**c**) Fluorescence microscopy images of primary rat hippocampal neurons expressing NES-HaloTag7-EGFP, incubated with 1 μM CA-MaPK-558 and imaged under no-wash conditions. Glutamate (500 μM) was added as indicated. EGFP channel (cyan, left) and MaPK-558 fluorescence (red hot, right). (**d**) Fluorescence time traces of neurons in (**c**). Shown are the mean CA-MaPK-558/EGFP ratio (solid line) and individual single-cell traces (faint lines), normalized to 1 at t = 0 min. The addition of glutamate is indicated by an arrow. n = 22 neurons from (**c**). Representative of three independent experiments. Scale bars: 50 μm.

### Bioluminescence as a readout

The indicators developed in this study have the potential to be integrated into bioluminescent metal ion indicators through combination with H-Luc^46^, a HaloTag–NanoLuc chimera that operates via bioluminescence resonance energy transfer (BRET). Labeling H-Luc with CA-MaPK-558 indeed resulted in efficient BRET from NanoLuc to the dye, producing dual emissions at 450 nm and 593 nm (Figure 6). Increasing K^+^ concentrations from 0-200 mM K^+^ resulted in an increase in light emission at 593 nm, with a ΔF/F_0_ value of 78%. Notably, its turn-on response contrasts with the turn-off of BRIPO^47^, a recently developed bioluminescent K^+^ indicator that integrates an engineered NanoLuc mutant with a K^+^-binding luciferin and which displays a decrease in bioluminescent signal upon K^+^ binding. For cellular applications, further engineering might be required to enhance the BRET response of CA-MaPK-619 and CA-MaPK-656. However, these experiments establish a versatile platform for developing bioluminescent K^+^ indicators with tunable and far-red-shifted emission spectra.

**Fig. 6.**
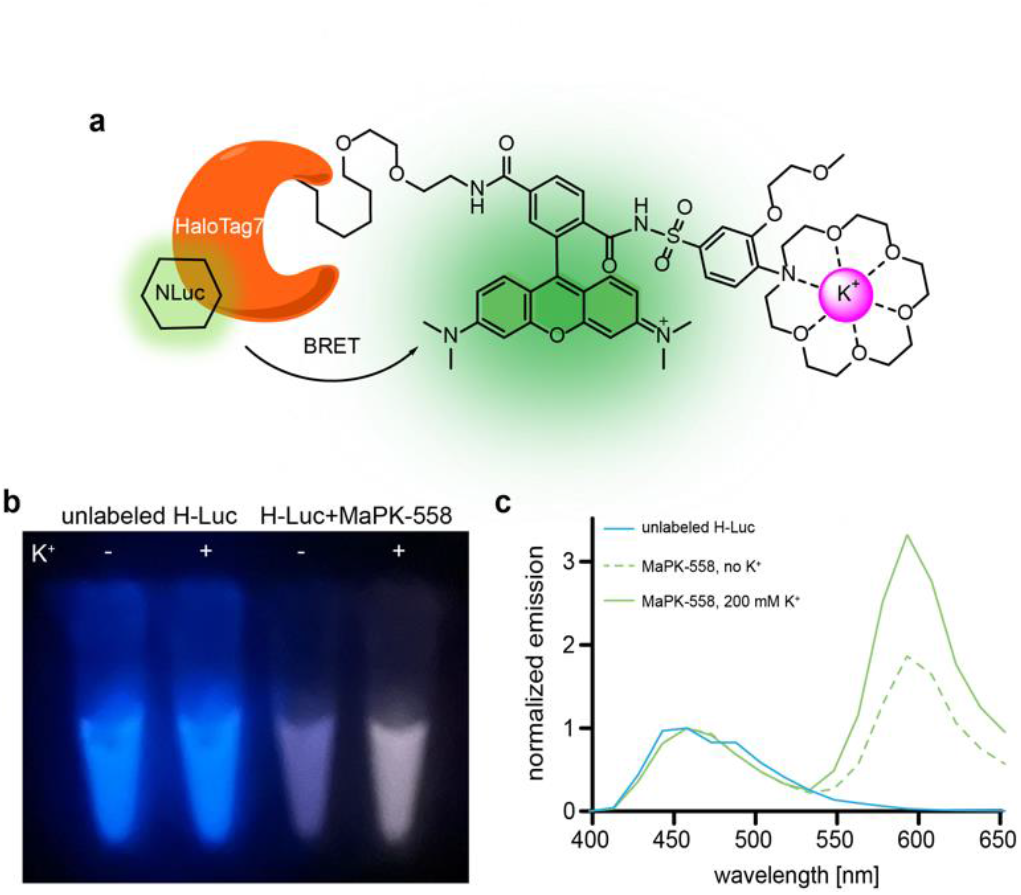
Characterization of bioluminescent K^+^ indicators. (**a**) Bioluminescent H-Luc transfer energy (BRET) to CA-MaPK-558. (**b**) Picture of H-Luc in Eppendorf tubes labeled with or without MaPK-558 in the absence or presence of 200 mM potassium. (**c**) Normalized *in vitro* emission spectra of H-Luc labeled MaPK-558, with and without potassium. Data was averaged from three technical replicates.

## Conclusion

We have reported a general strategy for developing chemigenetic metal ion indicators. From a chemistry perspective, our approach provides a short, modular synthetic route to construct fluorescent indicators built on the MaP dye scaffold and a PET-based turn-on mechanism. Eight indicators for metal ions were rapidly generated, including potassium, sodium, calcium, magnesium, zinc, iron(II), copper(I), and nickel. With our focus on potassium and sodium, we developed further several potassium and sodium indicators with different emission wavelengths, including far-red variants, as well as ligands compatible with self-labeling proteins such as HaloTag7 and SNAP-tag2. By conjugating MaPKs to HaloTag7 expressed in the cytoplasm or fused to PDGFR, we successfully monitored potassium concentration changes in both extracellular and intracellular environments. In comparison with previous reported K^+^ sensors, MaPKs offer a good response magnitude, brightness, spectral diversity and cellular targetability (Supplementary Figure 8 and Supplementary Table 3). A major advantage of MaPKs over previous designs is their good permeability and the potential for use without additional washing steps to remove unbound indicators. We demonstrated the applicability of MaPKs in rat hippocampal neurons, where glutamate stimulation induced potassium efflux, resulting in a measurable decrease in fluorescence intensity of cytosolically localized indicator.

Our strategy provides a versatile platform for developing fluorescent metal ion indicators based on a binding-induced fluorescence change mechanism. It could also be extended in the future to activity-based sensing^48^ for detecting metabolites and metal ions. Leveraging the broad spectrum of fluorogenic, spectrally distinguishable rhodamines, these MaP dye-based indicators could facilitate the simultaneous observation of multiple biological processes in cells and living organisms.

## Supporting information

Wang&Sun et al_MaPK_Supplementary information_final submission

## Acknowledgment

We thank A. Bergner for protein purification, J. Kress for the rAAV production, S. Kühn and J. Kompa for providing stable cell lines, B. Koch, N. Mertes, N. Porzberg, E. D’Este, E. Bonedeau and C. Gondrand for helpful discussion and technical assistance. We are grateful to the mass spectrometry facility (S. Fabritz, T. Rudi and J. Kling) of the MPIMR for its support. This work was funded by the Max Planck Society, Deutsche Forschungsgemeinschaft (DFG, grant no. SFB TRR 186). M.W. was supported by a Postdoc. Mobility fellowship of the Swiss National Science Foundation (P500PN_206863). D.S. was supported by a Humboldt Research Fellowship and an interinstitutional postdoctoral fellowship of The Health and Life Science Alliance Heidelberg Mannheim.

## Conflict of interest

K.J. is an inventor of the patent Cell-permeable fluorogenic fluorophores (EP18210676.5) which was filed by the Max Planck Society.

## Table of Contents (TOC)

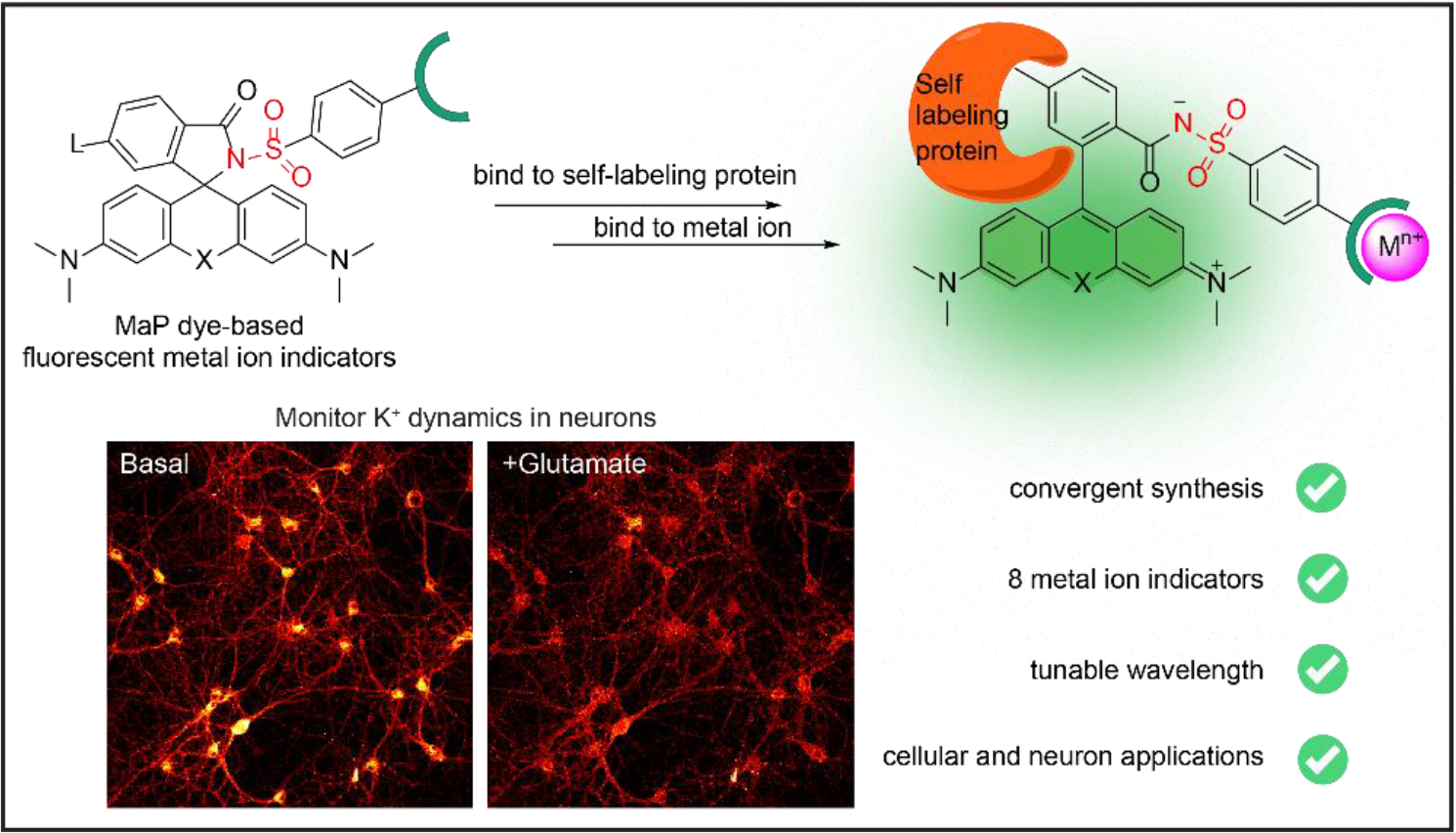

## Reference

(1) Rodriguez, R.; Müller, S.; Colombeau, L.; Solier, S.; Sindikubwabo, F.; Cañeque, T. Metal ion signaling in biomedicine. Chem. Rev. 2025, 125 (2), 660–744.

(2) Palmer, B. F. Regulation of potassium homeostasis. Clin. J. Am. Soc. Nephrol. 2015, 10 (6), 1050–1060.

(3) Domaille, D. W.; Que, E. L.; Chang, C. J. Synthetic fluorescent sensors for studying the cell biology of metals. Nat. Chem. Biol. 2008, 4 (3), 168–175.

(4) Carter, K. P.; Young, A. M.; Palmer, A. E. Fluorescent sensors for measuring metal ions in living systems. Chem. Rev. 2014, 114 (8), 4564–4601.

(5) Sambath, K.; Liu, X.; Wan, Z.; Hutnik, L.; Belfield, K. D.; Zhang, Y. Potassium ion fluorescence probes: Structures, properties and bioimaging. ChemPhotoChem 2021, 5 (4), 317–325.

(6) Minta, A.; Tsien, R. Y. Fluorescent indicators for cytosolic sodium. J. Biol. Chem. 1989, 264 (32), 19449–19457.

(7) Meuwis, K.; Boens, N.; De Schryver, F. C.; Gallay, J.; Vincent, M. Photophysics of the fluorescent K+ indicator PBFI. Biophys. J. 1995, 68 (6), 2469–2473.

(8) Padmawar, P.; Yao, X.; Bloch, O.; Manley, G. T.; Verkman, A. S. K+ waves in brain cortex visualized using a long-wavelength K+-sensing fluorescent indicator. Nat. Methods 2005, 2 (11), 825–827.

(9) Magzoub, M.; Padmawar, P.; Dix, J. A.; Verkman, A. S. Millisecond association kinetics of K+ with triazacryptand-based K+ indicators measured by fluorescence correlation spectroscopy. J. Phys. Chem. B 2006, 110 (42), 21216–21221.

(10) Namkung, W.; Padmawar, P.; Mills, A. D.; Verkman, A. S. Cell-based fluorescence screen for K+ channels and transporters using an extracellular triazacryptand-based K+ sensor. J. Am. Chem. Soc. 2008, 130 (25), 7794–7795.

(11) Zhou, X.; Su, F.; Tian, Y.; Youngbull, C.; Johnson, R. H.; Meldrum, D. R. A new highly selective fluorescent K+ sensor. J. Am. Chem. Soc. 2011, 133 (46), 18530–18533.

(12) Ning, J.; Tian, Y. Development of a new simple mitochondria-targeted fluorescent K+ sensor and the application in high-throughput monitoring K+ fluxes. Sens. Actuators B: Chem. 2020, 307, 127659.

(13) Wang, Z.; Detomasi, T. C.; Chang, C. J. A Dual-fluorophore sensor approach for ratiometric fluorescence imaging of potassium in living cells. Chem. Sci. 2021, 12 (5), 1720–1729.

(14) He, H.; Mortellaro, M. A.; Leiner, M. J. P.; Fraatz, R. J.; Tusa, J. K. A fluorescent sensor with high selectivity and sensitivity for potassium in water. J. Am. Chem. Soc. 2003, 125 (6), 1468–1469.

(15) Ast, S.; Schwarze, T.; Müller, H.; Sukhanov, A.; Michaelis, S.; Wegener, J.; Wolfbeis, O. S.; Körzdörfer, T.; Dürkop, A.; Holdt, H. A highly K+-selective phenylaza-[18]crown-6-lariatether-based fluoroionophore and its application in the sensing of K+ions with an optical sensor film and in cells. Chem. Eur. J. 2013, 19 (44), 14911–14917.

(16) Bischof, H.; Rehberg, M.; Stryeck, S.; Artinger, K.; Eroglu, E.; Waldeck-Weiermair, M.; Gottschalk, B.; Rost, R.; Deak, A. T.; Niedrist, T.; Vujic, N.; Lindermuth, H.; Prassl, R.; Pelzmann, B.; Groschner, K.; Kratky, D.; Eller, K.; Rosenkranz, A. R.; Madl, T.; Plesnila, N.; Graier, W. F.; Malli, R. Novel genetically encoded fluorescent probes enable real-time detection of potassium in vitro and in vivo. Nat Commun 2017, 8 (1), 1422.

(17) Shen, Y.; Wu, S.-Y.; Rancic, V.; Aggarwal, A.; Qian, Y.; Miyashita, S.-I.; Ballanyi, K.; Campbell, R. E.; Dong, M. Genetically encoded fluorescent indicators for imaging intracellular potassium ion concentration. Commun. Biol. 2019, 2 (1), 18.

(18) Wu, S.-Y.; Wen, Y.; Serre, N. B. C.; Laursen, C. C. H.; Dietz, A. G.; Taylor, B. R.; Drobizhev, M.; Molina, R. S.; Aggarwal, A.; Rancic, V.; Becker, M.; Ballanyi, K.; Podgorski, K.; Hirase, H.; Nedergaard, M.; Fendrych, M.; Lemieux, M. J.; Eberl, D. F.; Kay, A. R.; Campbell, R. E.; Shen, Y. A sensitive and specific genetically-encoded potassium ion biosensor for in vivo applications across the tree of Life. PLoS Biol. 2022, 20 (9), e3001772.

(19) Torres Cabán, C. C.; Yang, M.; Lai, C.; Yang, L.; Subach, F. V.; Smith, B. O.; Piatkevich, K. D.; Boyden, E. S. Tuning the sensitivity of genetically encoded fluorescent potassium indicators through structure-guided and genome mining strategies. ACS Sens. 2022, 7 (5), 1336–1346.

(20) Yang, L.; Pathiranage, V.; Zhou, S.; Sun, X.; Zhang, H.; Lai, C.; Gu, C.; Subach, F. V.; Drobizhev, M.; Walker, A. R.; Piatkevich, K. D. Sensitive red fluorescent indicators for real-time visualization of potassium ion dynamics in vivo. PLoS Biol. 2025, 23 (9), e3002993.

(21) Keppler, A.; Gendreizig, S.; Gronemeyer, T.; Pick, H.; Vogel, H.; Johnsson, K. A general method for the covalent labeling of fusion proteins with small molecules in vivo. Nat. Biotechnol. 2003, 21 (1), 86–89.

(22) Los, G. V.; Encell, L. P.; McDougall, M. G.; Hartzell, D. D.; Karassina, N.; Zimprich, C.; Wood, M. G.; Learish, R.; Ohana, R. F.; Urh, M.; Simpson, D.; Mendez, J.; Zimmerman, K.; Otto, P.; Vidugiris, G.; Zhu, J.; Darzins, A.; Klaubert, D. H.; Bulleit, R. F.; Wood, K. V. HaloTag: A novel protein labeling technology for cell imaging and protein analysis. ACS Chem. Biol. 2008, 3 (6), 373–382.

(23) Tsao, K. K.; Imai, S.; Chang, M.; Hario, S.; Terai, T.; Campbell, R. E. The best of both worlds: Chemigenetic fluorescent sensors for biological imaging. Cell Chem. Biol. 2024, 31 (9), 1652–1664.

(24) Kühn, S.; Nasufovic, V.; Wilhelm, J.; Kompa, J.; De Lange, E. M. F.; Lin, Y.-H.; Egoldt, C.; Fischer, J.; Lennoi, A.; Tarnawski, M.; Reinstein, J.; Vlijm, R.; Hiblot, J.; Johnsson, K. SNAP-Tag2 for faster and brighter protein labeling. Nat. Chem. Biol. 2025, 21, 1754–1761.

(25) Broch, F.; El Hajji, L.; Pietrancosta, N.; Gautier, A. Engineering of tunable allosteric-like fluorogenic protein sensors. ACS Sens. 2023, 8 (10), 3933–3942.

(26) Hirata, T.; Terai, T.; Yamamura, H.; Shimonishi, M.; Komatsu, T.; Hanaoka, K.; Ueno, T.; Imaizumi, Y.; Nagano, T.; Urano, Y. Protein-coupled fluorescent probe to visualize potassium ion transition on cellular membranes. Anal. Chem. 2016, 88 (5), 2693–2700.

(27) Cheng, D.; Ouyang, Z.; He, X.; Nasu, Y.; Wen, Y.; Terai, T.; Campbell, R. E. Highperformance chemigenetic potassium ion indicator. J. Am. Chem. Soc. 2024, 146 (51), 35117–35128.

(28) Wang, L.; Tran, M.; D’Este, E.; Roberti, J.; Koch, B.; Xue, L.; Johnsson, K. A general strategy to develop cell permeable and fluorogenic probes for multicolour nanoscopy. Nat. Chem. 2020, 12 (2), 165–172.

(29) Lardon, N.; Wang, L.; Tschanz, A.; Hoess, P.; Tran, M.; D’Este, E.; Ries, J.; Johnsson, K. Systematic tuning of rhodamine spirocyclization for super-resolution microscopy. J. Am. Chem. Soc. 2021, 143 (36), 14592–14600.

(30) Mertes, N.; Busch, M.; Huppertz, M.-C.; Hacker, C. N.; Wilhelm, J.; Gürth, C.-M.; Kühn, S.; Hiblot, J.; Koch, B.; Johnsson, K. Fluorescent and bioluminescent calcium indicators with tuneable colors and affinities. J. Am. Chem. Soc. 2022, 144 (15), 6928–6935.

(31) Deo, C.; Sheu, S.-H.; Seo, J.; Clapham, D. E.; Lavis, L. D. Isomeric tuning yields bright and targetable red Ca2+ indicators. J. Am. Chem. Soc. 2019, 141 (35), 13734–13738.

(32) Wang, M.-M.; Johnsson, K. Metal-free introduction of primary sulfonamide into electron-rich aromatics. Chem. Sci. 2024, 15 (31), 12310–12315.

(33) He, H.; Mortellaro, M. A.; Leiner, M. J. P.; Young, S. T.; Fraatz, R. J.; Tusa, J. K. A Fluorescent chemosensor for sodium based on photoinduced electron transfer. Anal. Chem. 2003, 75 (3), 549–555.

(34) Liu, M.; Yu, X.; Li, M.; Liao, N.; Bi, A.; Jiang, Y.; Liu, S.; Gong, Z.; Zeng, W. Fluorescent probes for the detection of magnesium ions (Mg2+): From design to application. RSC Adv. 2018, 8 (23), 12573–12587.

(35) Gee, K. R.; Zhou, Z.-L.; Ton-That, D.; Sensi, S. L.; Weiss, J. H. Measuring Zinc in Living Cells. Cell Calcium 2002, 31 (5), 245–251.

(36) Zhang, J.; Peng, X.; Wu, Y.; Ren, H.; Sun, J.; Tong, S.; Liu, T.; Zhao, Y.; Wang, S.; Tang, C.; Chen, L.; Chen, Z. Red- and far-red-emitting zinc probes with minimal phototoxicity for multiplexed recording of orchestrated insulin secretion. Angew Chem Int Ed 2021, 60 (49), 25846–25855.

(37) Zeng, L.; Miller, E. W.; Pralle, A.; Isacoff, E. Y.; Chang, C. J. A selective turn-on fluorescent sensor for imaging copper in living cells. J. Am. Chem. Soc. 2006, 128 (1), 10–11.

(38) Dodani, S. C.; He, Q.; Chang, C. J. A turn-on fluorescent sensor for detecting nickel in living cells. J. Am. Chem. Soc. 2009, 131 (50), 18020–18021.

(39) Petrat, F.; De Groot, H.; Rauen, U. Determination of the chelatable iron pool of single Intact cells by laser scanning microscopy. Arch. Biochem. Biophys. 2000, 376 (1), 74–81.

(40) Wilhelm, J.; Kühn, S.; Tarnawski, M.; Gotthard, G.; Tünnermann, J.; Tänzer, T.; Karpenko, J.; Mertes, N.; Xue, L.; Uhrig, U.; Reinstein, J.; Hiblot, J.; Johnsson, K. Kinetic and Structural Characterization of the self-labeling protein tags HaloTag7, SNAP-Tag, and CLIP-Tag. Biochem. 2021, 60 (33), 2560–2575.

(41) Yellen, G. The voltage-gated potassium channels and their relatives. Nature 2002, 419 (6902), 35–42.

(42) Köhling, R.; Wolfart, J. Potassium channels in epilepsy. Cold Spring Harb. Perspect. Med. 2016, 6 (5), a022871.

(43) Shen, Y.; Wu, S.-Y.; Rancic, V.; Aggarwal, A.; Qian, Y.; Miyashita, S.-I.; Ballanyi, K.; Campbell, R. E.; Dong, M. Genetically encoded fluorescent indicators for imaging intracellular potassium ion concentration. Commun. Biol. 2019, 2 (1), 18.

(44) Yu, S. P. NMDA Receptor-Mediated K+ Efflux and Neuronal Apoptosis. Science 1999, 284 (5412), 336–339.

(45) Kaech, S.; Banker, G. Culturing hippocampal neurons. Nat. Protoc. 2006, 1 (5), 2406–2415.

(46) Hiblot, J.; Yu, Q.; Sabbadini, M. D. B.; Reymond, L.; Xue, L.; Schena, A.; Sallin, O.; Hill, N.; Griss, R.; Johnsson, K. Luciferases with tunable emission wavelengths. Angew Chem Int Ed 2017, 56 (46), 14556–14560.

(47) Zhao, S.; Xiong, Y.; Sunnapu, R.; Zhang, Y.; Tian, X.; Ai, H. Bioluminescence imaging of potassium ion using a sensory luciferin and an engineered luciferase. J. Am. Chem. Soc. 2024, 146 (19), 13406–13416.

(48) Grover, K.; Koblova, A.; Pezacki, A. T.; Chang, C. J.; New, E. J. Small-molecule fluorescent probes for binding- and activity-based sensing of redox-active biological metals. Chem. Rev. 2024, 124 (9), 5846–5929.

