## Supplementary material for "Localizable fluorescent metal ion indicators with tunable colors": Wang&Sun et al_MaPK_Supplementary information_final submission

#### A general strategy to develop fluorescent metal ion indicators with tunable colors

### 1. Supplementary Figures

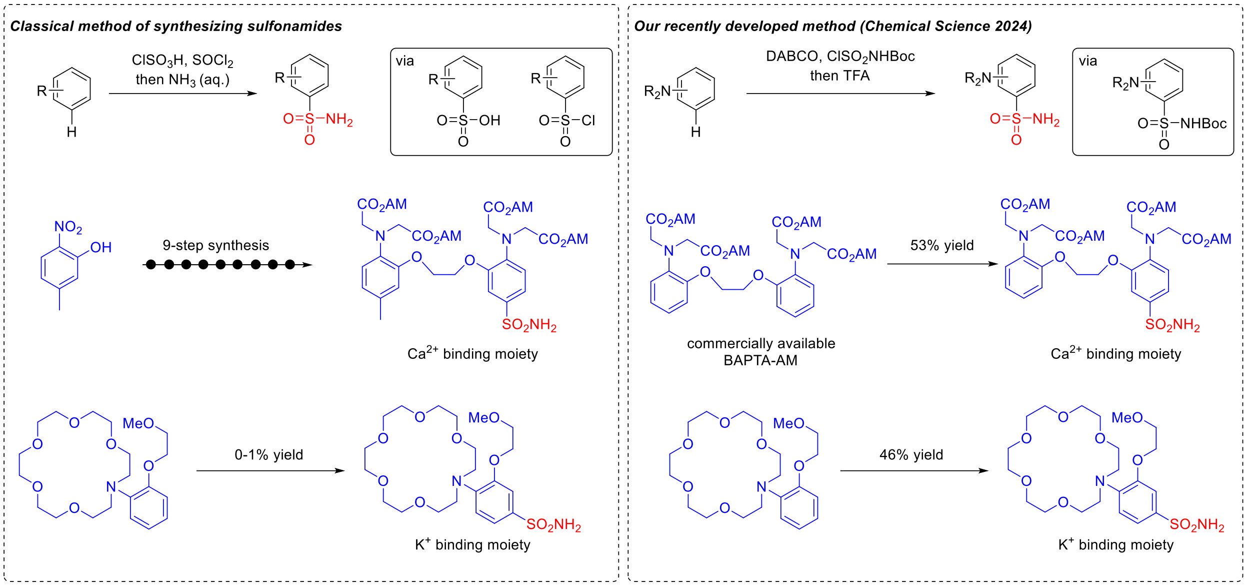

**Supplementary Fig. 1 | Comparison between the classical and our novel method for introducing a sulfonamide group to metal-ion chelators.**

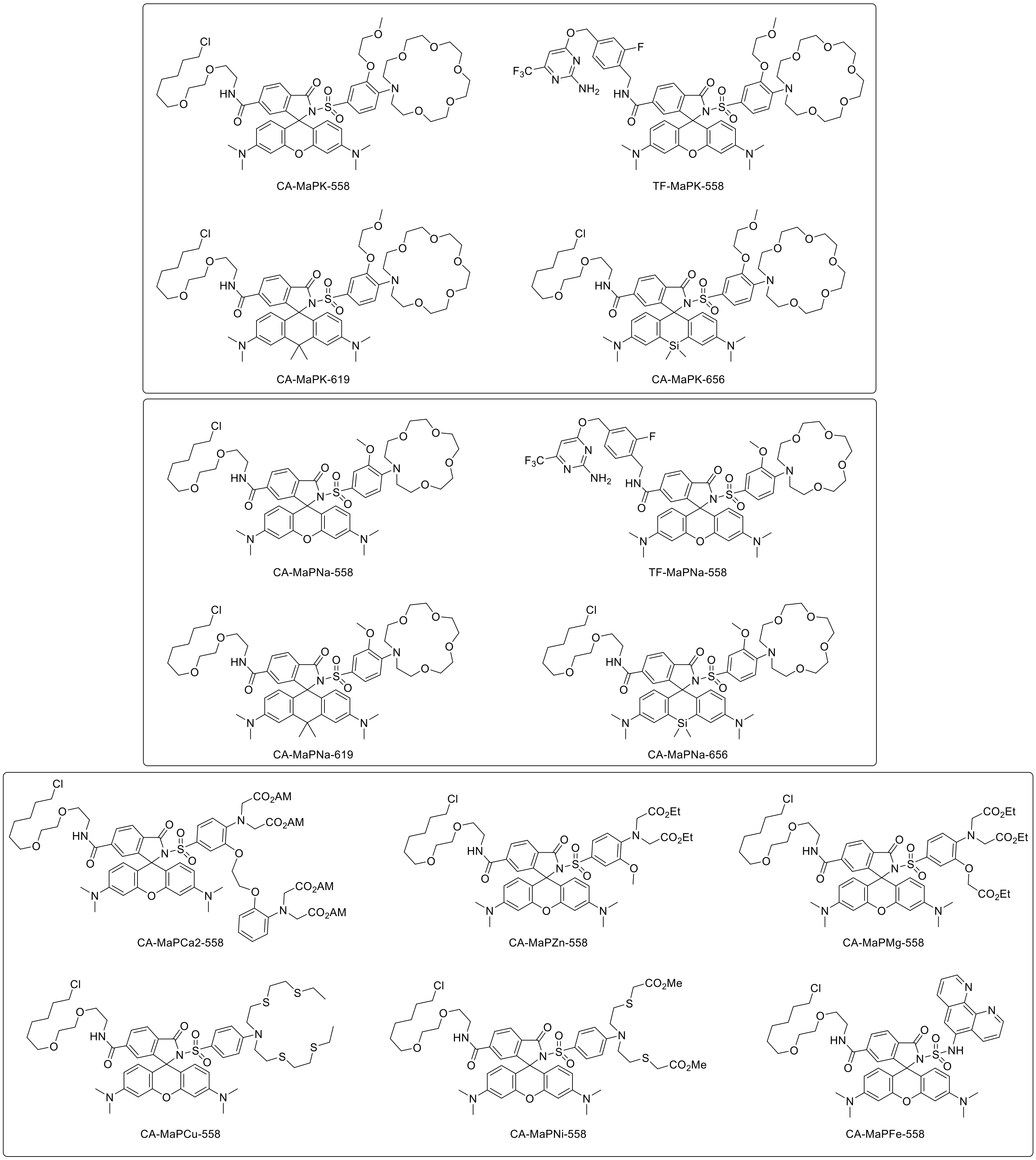

**Supplementary Fig. 2 | Chemical structures of the MaP dye-based metal ion indicators in this work.**

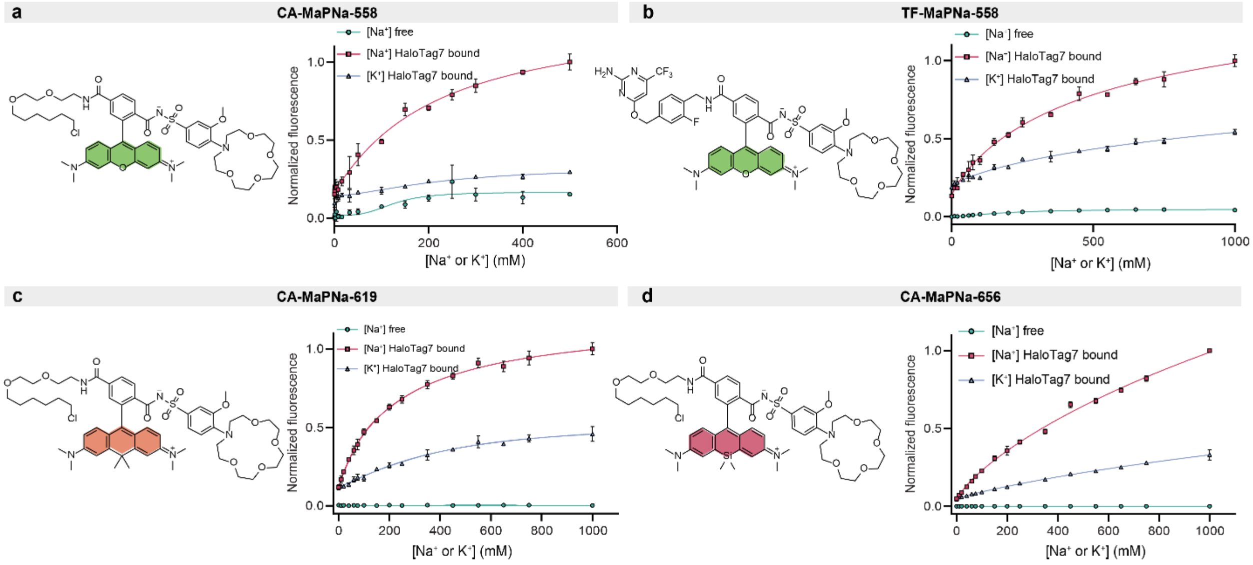

**Supplementary Fig. 3 | *In-vitro* titration curves of MaPNa indicators.** Sodium and potassium titration curves of MaPNa indicators. (a) is also shown in Fig. 2a. Error bars represent the mean ± standard deviation from three technical replicates.

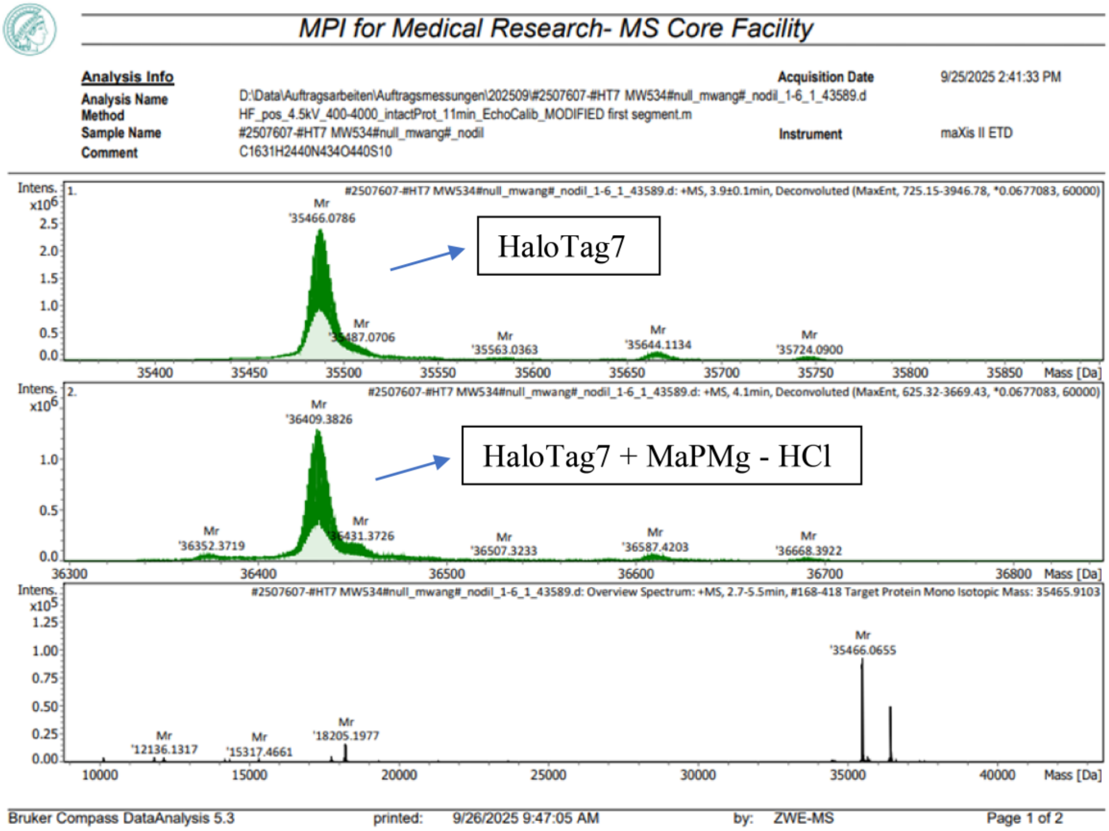

**Supplementary Fig. 4 | HRMS analysis showing efficient labeling of HaloTag7 with MaPMg.** HaloTag7 (400 µM) was labeled with MaPMg free acid (100 µM) for 2 h at r.t. before being diluted for HRMS analysis.

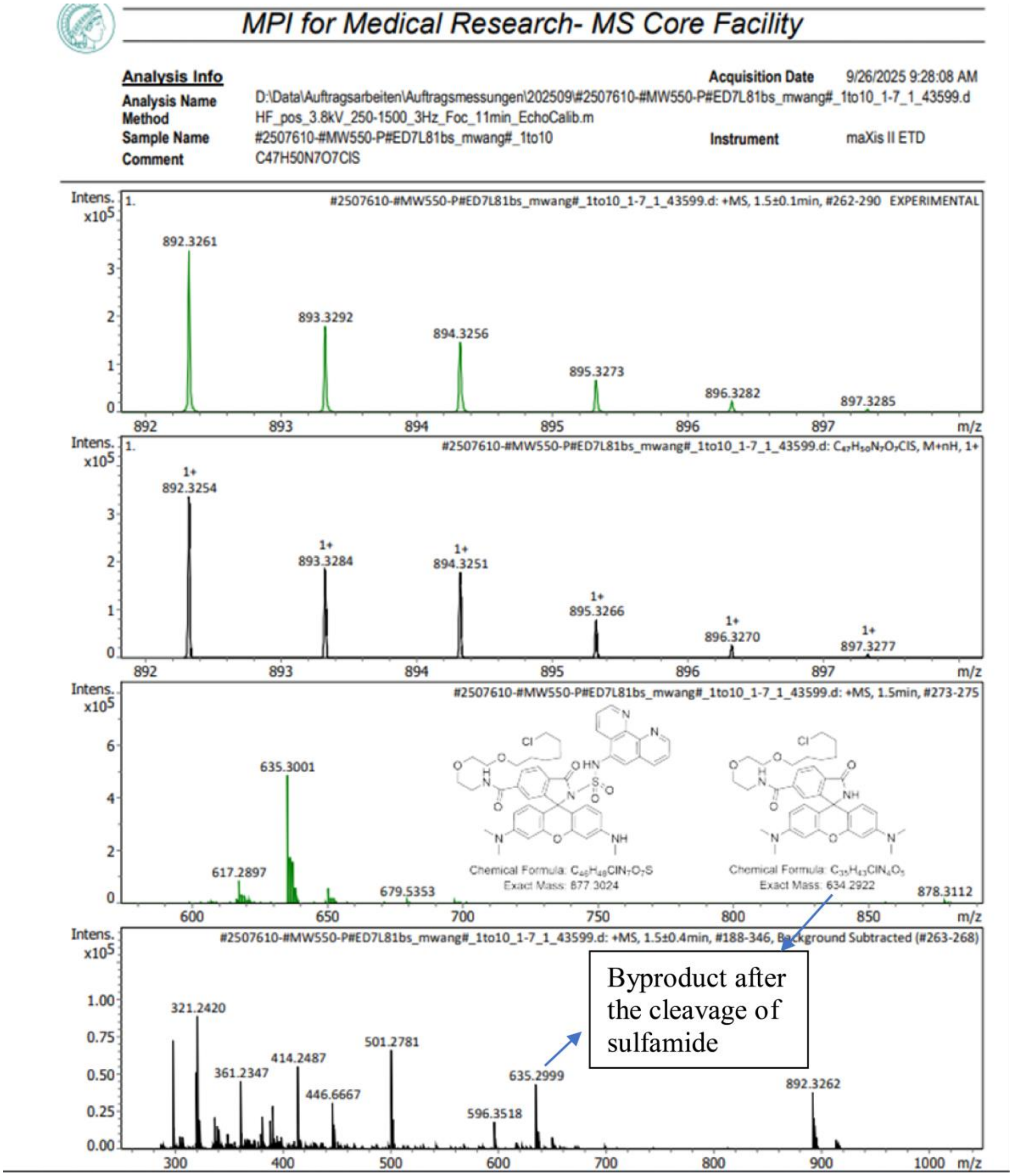

**Supplementary Fig. 5 | HRMS analysis of MaPFe degradation.** The major degradation product corresponds to the loss of 1,10-phenanthroline moiety through the sulfamide cleavage.

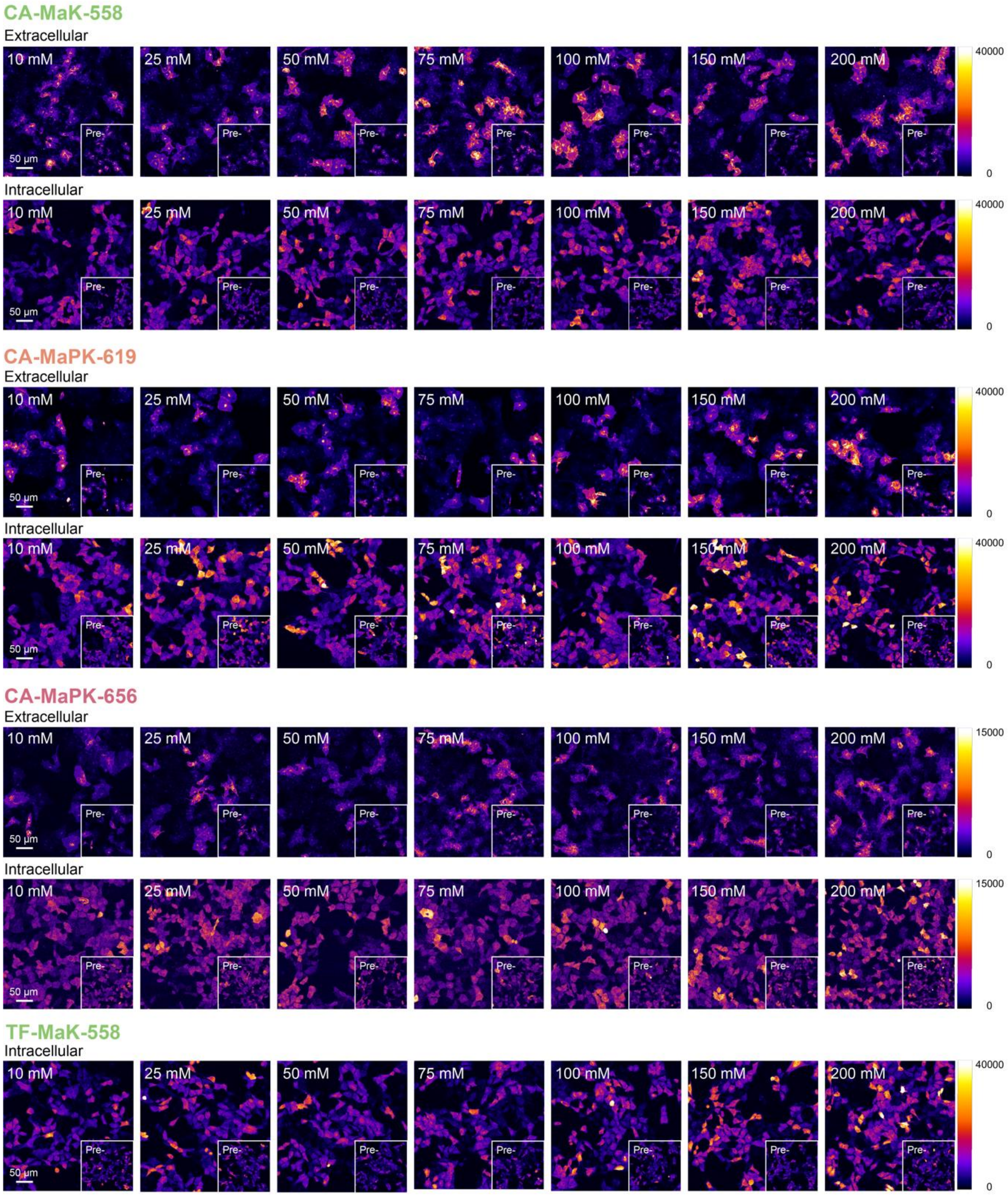

**Supplementary Fig. 6 | Extracellular and intracellular characterization of MaPKs in U2OS cells.** Fluorescence microscopy images of U2OS cells stably expressing Igκ-HaloTag7-PDGFR (extracellular) or HaloTag7–P30–SNAP-tag2–P2A–NLS–mTurquoise2 (intracellular), incubated with 1 μM MaPKs and imaged before and after the addition of different concentrations of KCl. Representative images from two (extracellular) or three (intracellular) independent experiments. Scale bars: 50 μm.

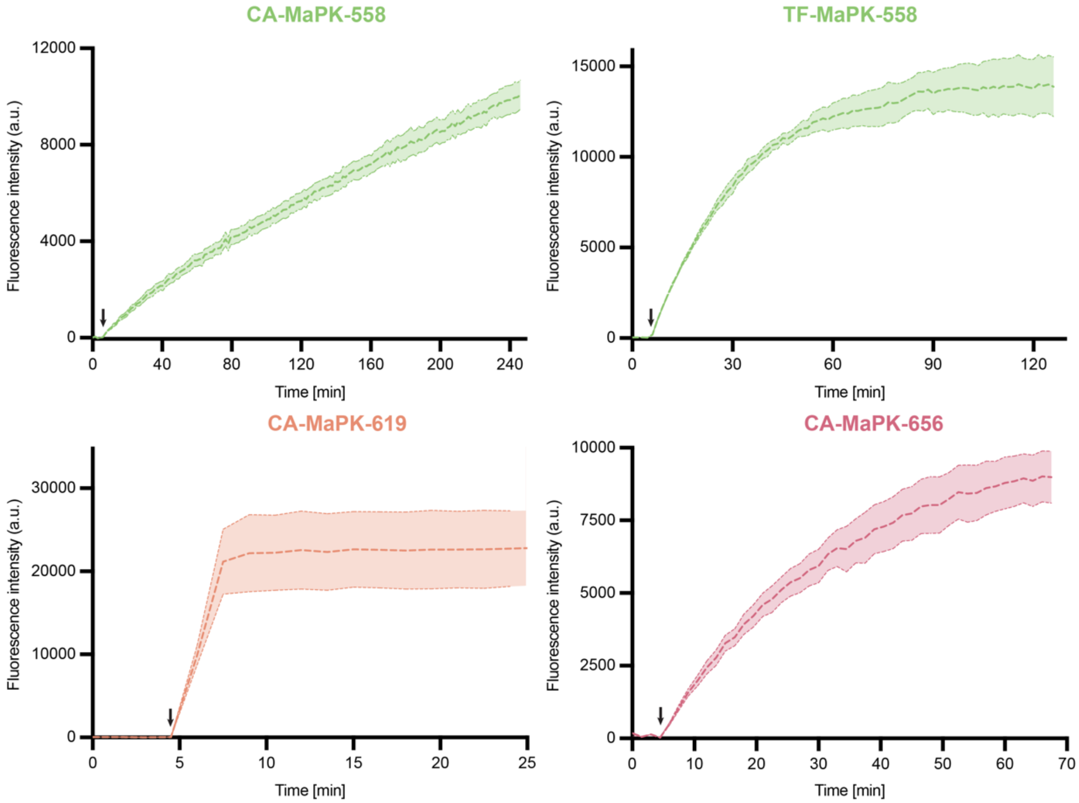

**Supplementary Fig. 7 | Labeling kinetics of MaPK indicators.** Time-resolved labeling of U2OS cells stably expressing HaloTag7–P30–SNAP-tag2–P2A–NLS–mTurquoise2. During time-lapse imaging, four baseline frames were acquired before the addition of the corresponding MaPK indicator (1 µM). Imaging was then continued after indicator addition. Background from unbound MaPKs was subtracted using regions without cells. Representative traces from two independent experiments.

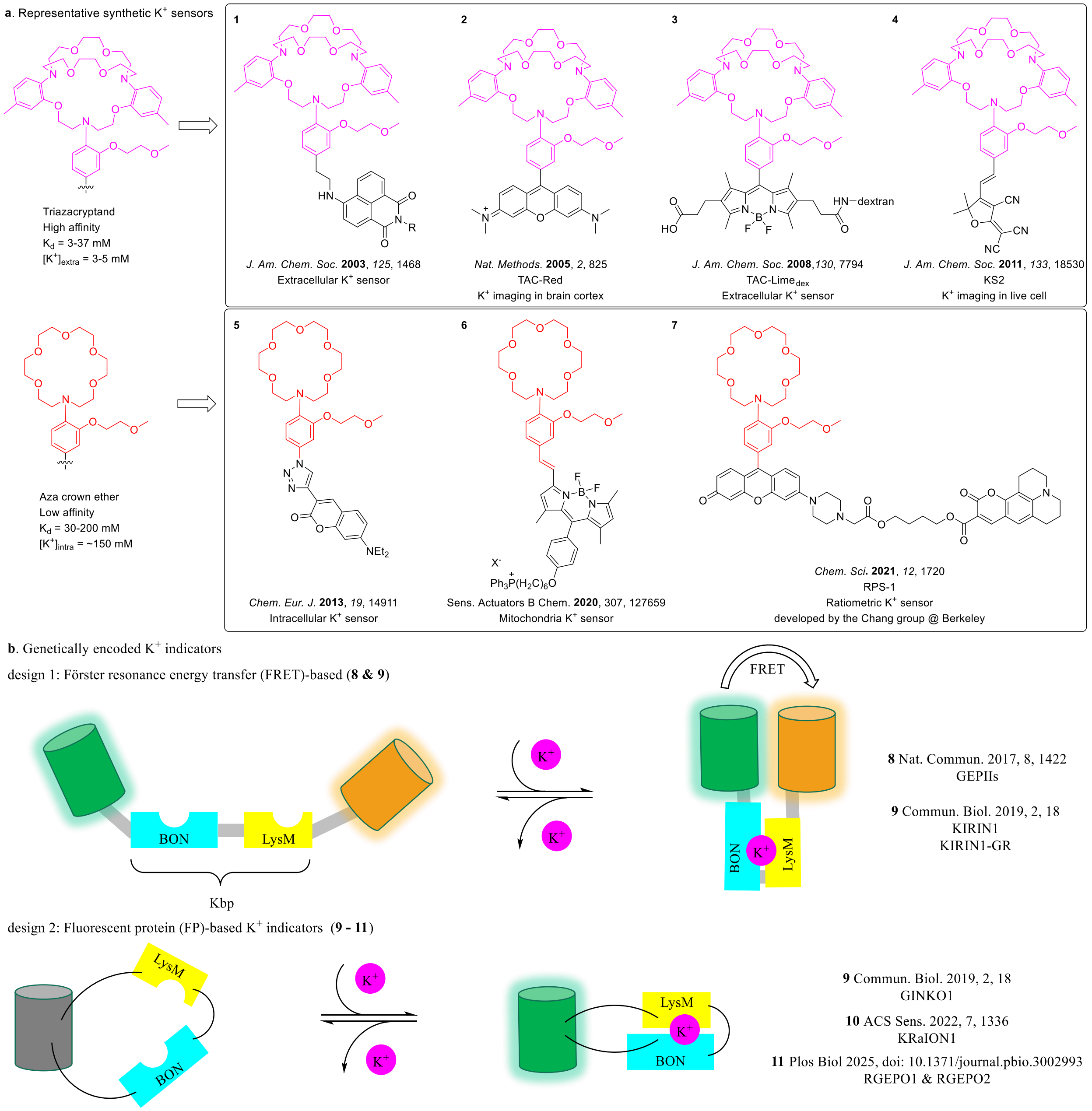
a. Representative synthetic K+ sensors
Triazacryptand
High affinity
Kd = 3-37 mM
[K+]extra = 3-5 mM
Aza crown ether
Low affinity
Kd = 30-200 mM
[K+]intra = ~150 mM
1
J. Am. Chem. Soc. 2003, 125, 1468
Extracellular K+ sensor
2
Nat. Methods. 2005, 2, 825
TAC-Red
K+ imaging in brain cortex
3
J. Am. Chem. Soc. 2008, 130, 7794
TAC-Limedex
Extracellular K+ sensor
4
J. Am. Chem. Soc. 2011, 133, 18530
KS2
K+ imaging in live cell
5
Chem. Eur. J. 2013, 19, 14911
Intracellular K+ sensor
6
Sens. Actuators B Chem. 2020, 307, 127659
Mitochondria K+ sensor
7
Chem. Sci. 2021, 12, 1720
RPS-1
Ratiometric K+ sensor
developed by the Chang group @ Berkeley
b. Genetically encoded K+ indicators
design 1: Förster resonance energy transfer (FRET)-based (8 & 9)
BON
LysM
Kbp
FRET
K+
8 Nat. Commun. 2017, 8, 1422
GEPIIs
9 Commun. Biol. 2019, 2, 18
KIRIN1
KIRIN1-GR
design 2: Fluorescent protein (FP)-based K+ indicators (9 - 11)
9 Commun. Biol. 2019, 2, 18
GINKO1
10 ACS Sens. 2022, 7, 1336
KRaION1
11 Plos Biol 2025, doi: 10.1371/journal.pbio.3002993
RGEPO1 & RGEPO2

**c**. Chemigenetic $K^+$ indicators

**12**

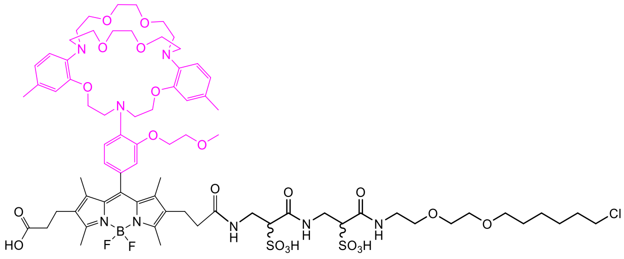

Anal. Chem. 2016, 88, 2693
TLSHalo

**13**

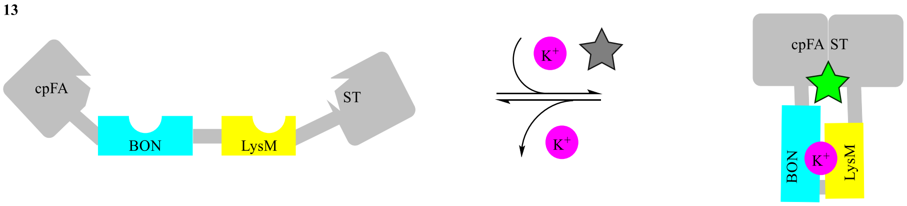

ACS Sens. 2023, 8, 3933
K+-FAST-5.1

**14**

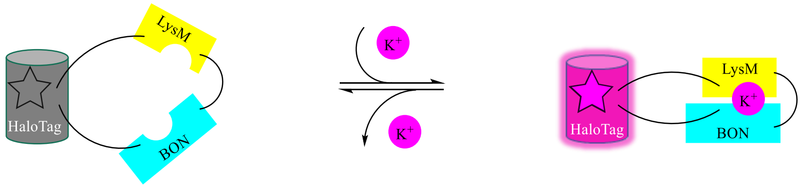

J. Am. Chem. Soc. 2024, 146, 35117
HaloKbp1

**15**

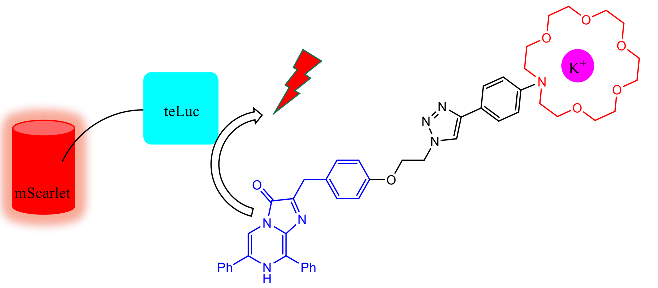

J. Am. Chem. Soc. 2024, 146, 13406
BRIPO

**Supplementary Fig. 8 | Schematic representation of existing $K^+$ indicators classified as synthetic (a), genetically encoded (b), and chemigenetic (c)**. A summary table comparing different properties can be found in Supplementary Table 3.

#### 2. Supplementary Tables

**Supplementary Table 1 | Photophysical properties of MaPNa indicators.**

| | **Fluorogenicity upon protein binding**[a] | **$F_{max}/F_0$ upon $Na^+$-binding**[b] | **$\lambda_{Ex}/\lambda_{Em}$ [nm]** | **$K_D(Na^+)$ [mM]**[b] | **Brightness [$mM^{-1}$ $cm^{-1}$]**[c] |
|---|---|---|---|---|---|
| CA-MaPNa-558 | 5.1 | 5.0 | 560/578 | 205 | 9 |
| CA-MaPNa-619 | 50 | 7.5 | 616/631 | 239 | 29 |
| CA-MaPNa-656 | 390 | 20 | 653/669 | >300 | 12 |
| TF-MaPNa-558 | 15 | 6.5 | 559/575 | >300 | 6 |

[a]Fluorescence increase at saturating sodium concentration.
[b]In protein-bound state (HaloTag7 or SNAP-tag2).
[c]At saturating sodium concentration and in protein-bound state.

**Supplementary Table 2 | Photophysical properties of MaP dye-based indicators for $Ca^{2+}$, $Mg^{2+}$, $Zn^{2+}$, $Cu^{+}$, $Fe^{2+}$ and $Ni^{2+}$.**

| | **Fluorogenicity upon protein binding**[a] | **$F_{max}/F_0$ upon ion-binding**[b] | **$\lambda_{Ex}/\lambda_{Em}$ [nm]** | **$K_D$**[b] | **Brightness [$mM^{-1}$ $cm^{-1}$]**[c] |
|---|---|---|---|---|---|
| MaPCa2 | 1.3 | 3.2 | 559/579 | 1.8 µM | 31 |
| MaPMg | 1.1 | 5.3 | 561/580 | 3.4 mM | 20 |
| MaPZn | 1.6 | 5.8 | 559/579 | 4.6 µM | 15 |
| MaPCu | 8 | 3.0 | 559/576 | 267 pM | 4 |
| MaPFe | 14 | 0.25 | 559/578 | 0.46 µM | 5[d] |
| MaPNi | NA | NA | NA | NA | NA |

[a]Fluorescence increase at saturating concentration of the corresponding metal ion.
[b]In HaloTag7-bound state.
[c]At saturating concentration of the corresponding metal ion and in HaloTag7-bound state.
[d]In buffer with no addition of $Fe^{2+}$ and in HaloTag7-bound state.

#### Supplementary Table 3 | Comparison of existing $K^+$ indicators and this work.

| Sensor types | Representative example* | Sensing domain | Ex/Em [nm] | $K_D$ [mM] | $\Delta F/F_0$ | Brightness [$mM^{-1}$ $cm^{-1}$] | Applications | Cellular localization | Sensing mechanism |
|---|---|---|---|---|---|---|---|---|---|
| **Synthetic** | He *et al.* JACS 2003 (**1**) | cryptand | 470/540 | 5 | | - | buffer | - | turn-on |
| | Verkman *et al.* Nat. Methods 2005 (**2**) | cryptand | 547/574 | - | 23 (200 mM $K^+$) | - | in vivo | extracellular | turn-on |
| | Verkman *et al.* JACS 2008 (**3**) | cryptand | 520/532 | - | - | - | cells | extracellular | turn-on |
| | Tian *et al.* JACS 2011 (**4**) | cryptand | 561/650 | 88 | 3 (140 mM $K^+$) | 0.2 | cells | intracellular, organelle non-specific | turn-on |
| | Dürkop, Holdt *et al.* Chem. Eur. J 2013 (**5**) | crown ether | 425/493 | 26 | 1.5 (100 mM $K^+$) | - | cells | intracellular, organelle non-specific | turn-on |
| | Tian *et al.* Sens. Actuators B Chem. 2011 (**6**) | crown ether | -/572 | 200 | 160 (1000 mM $K^+$) | 1.5 | cells | mitochondria | turn-on |
| | Chang *et al.* Chem. Sci 2021 (**7**) | crown ether | 525/552 | 137 | 6 (200 mM $K^+$) | 9.8 | cells | intracellular, organelle non-specific | FRET |
| **Genetically encoded** | Malli *et al.* Nat. Commun. 2017 (**8**) | Kbp | 413/475, 525 | 0.4-27 | 2.2 (1000 mM $K^+$) | - | cells, *in vivo* | intracellular, organelle-specific | FRET |
| | Shen, Campbell, Dong *et al.* Commun. Biol. 2019 (**9**) | Kbp | 502/514 | 0.4-2.56 | 1.5 (150 mM $K^+$) | 7.5 | cells, neurons | intracellular, organelle-specific | FRET/turn on |
| | Piatkevich, Boyden *et al.* ACS Sens. 2022 (**10**) | Kbp | 407, 504/517 | 69-138 | 4.3 (700 mM $K^+$) | 9-16 | cells | intracellular, organelle-specific | FRET |
| | Piatkevich, Walker *et al.* Plos Biol. 2025 (**11**) | Kbp | 575/599 | 2.4, 43.3 | 8.4, 5.4 (150 mM $K^+$) | 10.2, 6.0 | cells, neurons, *in vivo* | extracellular, cytoplasm | turn on |
| **Chemi-genetic** | Terai, Urano *et al.* Anal. Chem. 2016 (**12**) | cryptand | 522/537 | 19 | 10-20 (150 mM $K^+$) | - | cells | extracellular | turn on |
| | Gautier *et al.* ACS Sens. 2023 (**13**) | Kbp | -/~530 | 2.2 | 1.2 (~100 mM $K^+$) | - | buffer | - | turn on |
| | Terai, Campbell *et al.* JACS 2024 (**14**) | Kbp | 585/600 635/650 | 13-70 | 28-35 (1000 mM $K^+$) | 56-65 | cells | intracellular, organelle-specific | turn on |
| | Ai *et al.* JACS 2024 (**15**) | crown ether | -/~600 | 26 | 5 (150 mM $K^+$) | - | cells, neurons, *in vivo* | intracellular | bioluminescence /turn off |
| | **This work** | crown ether | 560/580 616/631 653/671 | 23-43 | 2-3 (150 mM $K^+$) | 16-56 | cells, neurons | extracellular, cytoplasm, organelle-specific | turn on |

*Schematic representation of existing $K^+$ indicators and details of reference **1-15** can be found in Supplementary Fig. 8.

### 3. Supplementary Note

**Supplementary Note 1 | Protein sequences**

>His–TEV–HaloTag7 purified protein

MHHHHHHHHHHENLYFQGIGTGFPFDPHYVEVLGERMHYVDVGPRDGTPVLFLHGNP
TSSYVWRNIIPHVAPTHRCIAPDLIGMGKSDKPDLGYFFDDHVRFMDAFIEALGLEEVVL
VIHDWGSALGFHWAKRNPERVKGIAFMEFIRPIPTWDEWPEFARETFQAFRTTDVGRKL
IIDQNVFIEGTLPMGVVRPLTEVEMDHYREPFLNPVDREPLWRFPNELPIAGEPANIVALV
EEYMDWLHQSPVPKLLFWGTPGVLIPPAEAARLAKSLPNCKAVDIGPGLNLLQEDNPDL
IGSEIARWLSTLEI

>Strep–Enterokinase CDS–SNAP-tag2–His purified protein

MASWSHPQFEKGADDDDKVPHMDKDCEMKRTTYDSPLGKLLLSGCEQGLHRIYFVGN
GQGEQGPPGPEPLMQATAWLNAYFHQPEAIEEFPVPALHHPVFQQESFTRQVLWKLLK
VVKFGEVISYSQLAALAGNPAATAAVKTALRGNPVPILIPCHRVVQGDGDVGGYEGPLY
VKEWLLAHEGHRLGKPGLGAPGFSSISAHHHHHHHHHH

>Igκ-chain leader sequence (IgκchL)–HA–HaloTag7–Myc–PDGFR transmembrane domain (PDGFRtmb) expressed in mammalian cells and cultured neurons

METDTLLLWVLLLWVPGSTGDYPYDVPDYAGAQPARSGSEIGTGFPFDPHYVEVLGER
MHYVDVGPRDGTPVLFLHGNPTSSYVWRNIIPHVAPTHRCIAPDLIGMGKSDKPDLGYF
FDDHVRFMDAFIEALGLEEVVLVIHDWGSALGFHWAKRNPERVKGIAFMEFIRPIPTWD
EWPEFARETFQAFRTTDVGRKLIIDQNVFIEGTLPMGVVRPLTEVEMDHYREPFLNPVDR
EPLWRFPNELPIAGEPANIVALVEEYMDWLHQSPVPKLLFWGTPGVLIPPAEAARLAKSL
PNCKAVDIGPGLNLLQEDNPDLIGSEIARWLSTLEISGGVDEQKLISEEDLNAVGQDTQE
VIVVPHSLPFKVVVISAILALVVLTIISLIILIMLWQKKPR

>HaloTag7–P30–SNAP-tag2–P2A–NLS–mTurquoise2 expressed in mammalian cells

MGSEIGTGFPFDPHYVEVLGERMHYVDVGPRDGTPVLFLHGNPTSSYVWRNIIPHVAPT
HRCIAPDLIGMGKSDKPDLGYFFDDHVRFMDAFIEALGLEEVVLVIHDWGSALGFHWA
KRNPERVKGIAFMEFIRPIPTWDEWPEFARETFQAFRTTDVGRKLIIDQNVFIEGTLPMGV
VRPLTEVEMDHYREPFLNPVDREPLWRFPNELPIAGEPANIVALVEEYMDWLHQSPVPK
LLFWGTPGVLIPPAEAARLAKSLPNCKAVDIGPGLNLLQEDNPDLIGSEIARWLSTLEISG
SGRPPPPPPPPPPPPPPPPPPPPPPPPPPPPPPGGRSRSLEMDKDCEMKRTTYDSPLGKLLLS
GCEQGLHRIYFVGNGQGEQGPPGPEPLMQATAWLNAYFHQPEAIEEFPVPALHHPVFQQ
ESFTRQVLWKLLKVVKFGEVISYSQLAALAGNPAATAAVKTALRGNPVPILIPCHRVVQ
GDGDVGGYEGPLYVKEWLLAHEGHRLGKPGLGDNGSGATNFSLLKQAGDVEENPGPS
RMAPKKKRKVMVSKGEELFTGVVPILVELDGDVNGHKFSVSGEGEGDATYGKLTLKFI
CTTGKLPVPWPTLVTTLSWGVQCFARYPDHMKQHDFFKSAMPEGYVQERTIFFKDDGN
YKTRAEVKFEGDTLVNRIELKGIDFKEDGNILGHKLEYNYFSDNVYITADKQKNGIKAN

FKIRHNIEDGGVQLADHYQQNTPIGDGPVLLPDNHYLSTQSKLSKDPNEKRDHMVLLEF
VTAAGITLGMDELYK

>HaloTag7–P30–SNAP-tag2–NLS3×–P2A–NLS–mTurquoise2 expressed in mammalian cells

MGSEIGTGFPFDPHYVEVLGERMHYVDVGPRDGTPVLFLHGNPTSSYVWRNIIPHVAPT
HRCIAPDLIGMGKSDKPDLGYFFDDHVRFMDAFIEALGLEEVVLVIHDWGSALGFHWA
KRNPERVKGIAFMEFIRPIPTWDEWPEFARETFQAFRTTDVGRKLIIDQNVFIEGTLPMGV
VRPLTEVEMDHYREPFLNPVDREPLWRFPNELPIAGEPANIVALVEEYMDWLHQSPVPK
LLFWGTPGVLIPPAEAARLAKSLPNCKAVDIGPGLNLLQEDNPDLIGSEIARWLSTLEISG
SGRPPPPPPPPPPPPPPPPPPPPPPPPPPPPPPGGRSRSLEMDKDCEMKRTTYDSPLGKLLLS
GCEQGLHRIYFVGNGQGEQGPPGPEPLMQATAWLNAYFHQPEAIEEFPVPALHHPVFQQ
ESFTRQVLWKLLKVVKFGEVISYSQLAALAGNPAATAAVKTALRGNPVPILIPCHRVVQ
GDGDVGGYEGPLYVKEWLLAHEGHRLGKPGLGGAPDPKKKRKVDPKKKRKVDPKKK
RKELRASPQATNFSLLKQAGDVEENPGPSRMAPKKKRKVMVSKGEELFTGVVPILVELD
GDVNGHKFSVSGEGEGDATYGKLTLKFICTTGKLPVPWPTLVTTLSWGVQCFARYPDH
MKQHDFFKSAMPEGYVQERTIFFKDDGNYKTRAEVKFEGDTLVNRIELKGIDFKEDGNI
LGHKLEYNYFSDNVYITADKQKNGIKANFKIRHNIEDGGVQLADHYQQNTPIGDGPVLL
PDNHYLSTQSKLSKDPNEKRDHMVLLEFVTAAGITLGMDELYK

>NES–HaloTag7–mEGFP expressed in cultured neurons

MLQNELALKLAGLDINKTGGSGSEIGTGFPFDPHYVEVLGERMHYVDVGPRDGTPVLFL
HGNPTSSYVWRNIIPHVAPTHRCIAPDLIGMGKSDKPDLGYFFDDHVRFMDAFIEALGLE
EVVLVIHDWGSALGFHWAKRNPERVKGIAFMEFIRPIPTWDEWPEFARETFQAFRTTDV
GRKLIIDQNVFIEGTLPMGVVRPLTEVEMDHYREPFLNPVDREPLWRFPNELPIAGEPANI
VALVEEYMDWLHQSPVPKLLFWGTPGVLIPPAEAARLAKSLPNCKAVDIGPGLNLLQED
NPDLIGSEIARWLSTLEISGMVSKGEELFTGVVPILVELDGDVNGHKFSVSGEGEGDATY
GKLTLKFICTTGKLPVPWPTLVTTLTYGVQCFSRYPDHMKQHDFFKSAMPEGYVQERTI
FFKDDGNYKTRAEVKFEGDTLVNRIELKGIDFKEDGNILGHKLEYNYNSHNVYIMADK
QKNGIKVNFKIRHNIEDGSVQLADHYQQNTPIGDGPVLLPDNHYLSTQSKLSKDPNEKR
DHMVLLEFVTAAGITLGMDELYK

>H-Luc purified protein

Strep–FLAG–H-Luc–(NanoLuc circular permuted at positions 67-68)-His-tag

MASWSHPQFEKGADDDDKVPHGSEIGTGFPFDPHYVEVLGERMHYVDVGPRDGTPVLF
LHGNPTSSYVWRNIIPHVAPTHRCIAPDLIGMGKSDKPDLGYFFDDHVRFMDAFIEALGL
EEVVLVIHDWGSALGFHWAKRNPERVKGIAFMEFIRPIPTWDEWPEFARETFQAFRTSG
DQMGQIEKIFKVVYPVDDHHFKVILHYGTLVIDGVTPNMIDYFGRPYEGIAVFDGKKITV
TGTLWNGNKIIDERLINPDGSLLFRVTINGVTGWRLCERILAGGTGGSGGTGGSMVFTLE
DFVGDWRQTAGYNLDQVLEQGGVSSLFQNLGVSVTPIQRIVLSGENGLKIDIHVIIPYEG
LDVGRKLIIDQNVFIEGTLPMGVVRPLTEVEMDHYREPFLNPVDREPLWRFPNELPIAGE
PANIVALVEEYMDWLHQSPVPKLLFWGTPGVLIPPAEAARLAKSLPNCKAVDIGPGLNL
LQEDNPDLIGSEIARWLSTLEISGAPGFSSISAHHHHHHHHHH

### 4. General Experimental Information

**Chemical synthesis and general information.** For quantitative flash chromatography, distilled technical grade solvents were used. THF, $Et_2O$, toluene, hexane and $CH_2Cl_2$ were dried by passage over activated alumina under nitrogen atmosphere ($H_2O$ content < 7 ppm, Karl-Fischer titration). All chemicals were purchased and used as received unless stated otherwise. Chromatographic purification was performed as flash chromatography using Macherey-Nagel silica 40-63, 60 Å or Alumina neutral 55 Å, using the solvents indicated as eluent with 0.1-0.5 bar pressure. TLC was performed on Merck silica gel 60 F254 TLC plastic or aluminium plates and visualized with UV light, permanganate stain. Preparative RP-HPLC was performed on an UltiMate 3000 system (Thermo Fisher Scientific) using a C18 5 µm, 21.2 × 250 mm column (flow rate 8 mL $min^{-1}$, Supelco). A typical run was over 60 min, solvent ratios are given in the procedures (solvent A: 0.1% TFA in water, solvent B: MeCN). Compounds were dried under high vacuum using a lyophilizer (Christ) equipped with a vacuum pump (Vacuubrand).

$^1$H-NMR spectra were recorded at room temperature on a Brucker DPX-400 400 MHz spectrometer in $CDCl_3$, Acetone-$d_6$, $CD_3CN$, DMSO-$d_6$ or $CD_3OD$, all signals are reported in ppm with the internal chloroform signal at 7.26 ppm, the internal acetone signal at 2.09 ppm, the internal acetonitrile signal at 1.94 ppm, the internal acetonitrile signal at 2.50 ppm and the internal methanol signal at 3.34 ppm as standard. The data is being reported as (s = singlet, d = doublet, t = triplet, q = quadruplet, p = quintet, m = multiplet or unresolved, br = broad signal, integration, coupling constant(s) in Hz, interpretation). $^{13}$C-NMR spectra were recorded with 1H-decoupling on a Brucker DPX-400 101 MHz spectrometer in $CDCl_3$, Acetone-$d_6$, $CD_3CN$, DMSO-$d_6$ or $CD_3OD$, all signals are reported in ppm with the internal chloroform signal at 77.0 ppm, Acetone-$d_6$ signal at 29.8 ppm, $CD_3CN$ signal at 1.3 ppm, $CD_3CN$ signal at 39.5 ppm or $CD_3OD$ signal at 49.0 ppm as standard. High resolution mass spectrometric measurements were performed by the mass spectrometry service of MPIMR on a MICROMASS (ESI) Q-TOF Ultima API.

**Molecular cloning.** Plasmids encoding HaloTag7 and SNAP-tag2 for protein expression in bacterial and mammalian systems, including various subcellular localizations (e.g., nucleus, cytosol, outer membrane leaflet), have been previously described[1–3]. For bacterial protein expression, a pET51b(+) vector (Novagen) was used. For mammalian expression, pcDNA5/FRT or pcDNA5/FRT/TO vectors (Thermo Fisher Scientific) were employed, either for transient transfection or for the generation of stable cell lines. The pAAV-hSyn vector was used for recombinant adeno-associated virus (rAAV) production. All cloning procedures were performed using Gibson Assembly, following the manufacturer's instructions. For cloning of pAAV plasmids, NEB Stable Competent *E. coli* cells were used. All constructs were verified by Sanger sequencing. For virus-related plasmids, the integrity of the inverted terminal repeats (ITRs) and recombination sites was further confirmed by full-plasmid sequencing (Microsynth) following MaxiPrep DNA extraction (Qiagen).

**Protein expression and purification.** HaloTag7, SNAP-tag2 and H-Luc proteins were expressed and purified as previously described[1,2]. Briefly, pET51b(+) plasmids were transformed into E. coli BL21(DE3)-pLysS (Novagen) cells. Cultures were grown in Luria–Bertani (LB) medium supplemented with ampicillin at 37°C until reaching an optical density at 600 nm ($OD_{600}$) of 0.6–0.8. Protein expression was induced with 0.5 mM isopropyl-β-d-1-thiogalactopyranoside (IPTG) and carried out overnight at 16°C. Cells were harvested by centrifugation (4,500 g, 4°C, 10 min), resuspended in His-tag extraction buffer (50 mM $KH_2PO_4$, 150 mM NaCl, 5 mM imidazole, 1 mM PMSF,0.25 mg $mL^{-1}$ lysozyme, pH 8.0), and lysed by sonication on wet ice. Proteins were purified by immobilized metal ion affinity chromatography (IMAC), using either gravity-flow columns or an ÄKTA Pure FPLC system (Cytiva) equipped with HisTrap FF crude columns (Cytiva). Buffer exchange to activity buffer (50 mM HEPES, 50 mM NaCl, pH 7.3) was performed using Zeba spin desalting columns (7-kDa MWCO, 0.5 mL; Thermo Fisher Scientific) or HiPrep 26/10 desalting columns (Cytiva) on the FPLC system. Proteins were concentrated using Amicon Ultra centrifugal filters (10-kDa MWCO; Millipore). Purity and molecular weight were confirmed by SDS–PAGE and high-resolution mass spectrometry (HRMS). Final protein preparations were aliquoted, flash-frozen in liquid nitrogen, and stored at −80°C.

**Optical Spectroscopy.** Fluorescent indicators were prepared as stock solutions in anhydrous DMSO and diluted to ensure a final DMSO content below 1% (v/v) in all assays. Fluorescence, absorbance and extinction coefficient measurements were performed on a Spark® 20M plate reader (Tecan) equipped with filters and a monochromator. Absolute quantum yields were determined on a Quantaurus-QY spectrometer (model C11374, Hamamatsu) using diluted samples ($A \approx 0.1$). All reported values represent averages of independent measurements or wells ($n \geq 3$).

**Fluorescence and absorbance measurements.** Fluorescent indicators (100 µM) were pre-incubated with protein (400 µM; HaloTag7, SNAP-tag2, or BSA) for 2 h at room temperature (r.t.) in 1× PBS. The resulting dye–protein complexes were diluted to a final dye concentration of 1 µM in 100 µL buffer containing the indicated metal ion at varying concentrations in non-binding black flat-bottom 96-well plates (PerkinElmer). Note: HaloTag7 protein with His-tag removed was used for the titration of MaPCu and MaPFe.

Metal ion buffers were prepared as follows:

- $K^+$ (for MaPK): by mixing different proportions of buffers containing 200 mM KCl or 200 mM choline chloride.
- $Na^+$ (for MaPNa): by mixing different proportions of buffers containing 1 M NaCl or 1 M choline chloride.
- $Ca^{2+}$ (for MaPCa): by mixing buffers from a commercial kit (Invitrogen, Cat. no. C3008MP) following the vendor's protocol: EGTA buffer (30 mM MOPS, 10 mM EGTA, 100 mM KCl, pH 7.2) and Ca-EGTA buffer (30 mM MOPS, 10 mM Ca-EGTA, 100 mM KCl, pH 7.2).
- $Mg^{2+}$ (for MaPMg): by mixing different proportions of buffers containing 20 mM $MgCl_2$ or 20 mM choline chloride.
- $Zn^{2+}$ (for MaPZn): by mixing different proportions of buffers containing 1 mM $ZnCl_2$ or 1 mM choline chloride.
- $Fe^{2+}$ (for MaPFe): by mixing different proportions of buffers containing 10 µM $FeSO_4$ (with 40 µM ascorbate) or 10 µM choline chloride.
- $Cu^+$ (for MaPCu): using thiourea as a competitive ligand to generate buffered solutions containing 0–2000 pM [$Cu^+$]. Stability constants for thiourea binding were taken from the

literature[4,5]: $\beta_{12} = 2.0 \times 10^{12}$, $\beta_{13} = 2.0 \times 10^{14}$, $\beta_{14} = 3.4 \times 10^{15}$. Cu(I) was delivered as $[Cu(MeCN)_4][PF_6]$ from a 20 mM acetonitrile stock.

Fluorescence spectra were recorded after 30 min incubation at r.t. with slow shaking on a Spark® 20M plate reader (Tecan) using the following excitation wavelengths: 520 nm (TMR-variants), 570 nm (CPY-variants) and 610 nm (SiR-variants) (bandwidth: 10 nm, step size: 1 nm). Absorbance spectra were obtained analogously in non-binding flat bottom transparent 96-well plates (ThermoScientific), at 5 µM final dye concentration (bandwidth: 1 nm). Dissociation constants $K_D(M^{n+})$ were calculated by linear regression of log $[(F - F_{min})/(F_{max} - F)]$ versus log $[M^{n+}]$, with the x-intercept corresponding to log $K_D(M^{n+})$.

**Bioluminescence measurements.** H-Luc protein (2 µM) was labeled with excess CA-MaPK-558 (10 µM) for 1 hour at r.t. The dye—protein mixture was diluted to 2 nM final protein concentration in 100 µL activity buffer supplemented with 1 mg $mL^{-1}$ BSA and NanoGlow furimazine solution from Promega (final concentrations: H-Luc: 2 nM, CA-MaPK-558: 10 nM, NanoGlow: 1000×, BSA: 1 mg $mL^{-1}$). Luminescence emission profile of each well were recorded on a microplate reader (Spark20M, Tecan) over the range 400-650 nm (18 measurements every 15 nm, BW 25 nm). For the photos of Figure 6b, the final H-Luc concentration was at 20 nM with 100 nM CA-MaPK-558, 1000× NanoGlow and 1 mg $mL^{-1}$ BSA.

**Cell culture.** U2OS Flp In T-REx and human embryonic kidney 293 (HEK293) Lenti-X cells were cultured in Dubeco's Modified Eagle Medium (DMEM, Gibco, 31966021) supplemented with 10% fetal bovine serum (FBS, Gibco) in a humidified incubator at 37°C with 5% $CO_2$. For imaging experiments, cells were maintained in phenol red-free DMEM (referred to as imaging medium; Gibco, 31053-028) supplemented with 1× GlutaMAX (Gibco, 35050-038), 1 mM sodium pyruvate (Gibco, 11360-070) and 10% FBS. Primary rat hippocampal neurons were cultured in Neurobasal medium (Gibco, 12348-017) supplemented with 1× GlutaMAX, 1× B-27 (Gibco, 17504044), 100 units $mL^{-1}$ penicillin, and 100 µg $mL^{-1}$ streptomycin (Gibco, 15140122). All cell lines were routinely tested for mycoplasma contamination and confirmed to be mycoplasma-free.

**Stable cell line establishment.** U2OS cell lines stably expressing either IgKchL-HA-HaloTag7-myc-PDGFRtmb, HaloTag7–P30–SNAP-tag2–NLS–P2A–NLS–mTurquoise2 and HaloTag7–P30–SNAP-tag2–P2A–NLS–mTurquoise2 were generated using the Flp-In system, as previously described[1,2]. Briefly, U2OS Flp-In cells were cultured to around 70% confluency in T-25 flasks and co-transfected with 440 ng of a pcDNA5/FRT/TO plasmid encoding the gene of interest (GOI) and 3,560 ng of the pOG44 Flp-recombinase expression plasmid (Invitrogen), using Lipofectamine 3000 according to the manufacturer's instructions. Twelve hours post-transfection, the medium was replaced, and after 24 hours, hygromycin B (Carl Roth, 250-545-5, 100 µg $mL^{-1}$) was added to select for cells that had stably integrated the GOI into the genome. Following 48–72 hours of selection, surviving cells were recovered in fresh medium until reaching confluency. Cells with high GOI expression were enriched by bulk fluorescence-activated cell sorting (FACS) using a FACSMelody cell sorter (BD Biosciences), based on either fluorescent protein signal (mEGFP; FITC filter) or HaloTag7/SNAP-tag2 labeling (SiR; APC filter).

**In-cell labeling kinetic measurements of MaPKs.** U2OS cells stably expressing HaloTag7–P30–SNAP-tag2–P2A–NLS–mTurquoise2 were seeded into eight-well imaging chambered coverslips with glass bottoms (ibidi, 80827) and cultured in 200 µL imaging medium per well. During timelapse imaging, four baseline frames were first captured. Then, the medium was exchanged with 200 µL of imaging medium containing 1 µM MaPKs, the cells were imaged every 90 s on a commercial Leica Stellaris 5 confocal microscope. Signal background from free MaPKs was subtracted using area without cell culture.

**Live cell imaging of MaPKs in U2OS cells.** U2OS cell lines were seeded into eight-well imaging chambered coverslips with glass bottoms and cultured in 200 µL imaging medium per well. Cells were labeled with 1 µM MaPKs for 2 h and then refreshed with imaging medium prior to imaging. For extracellular MaPK imaging, U2OS cells stably expressing IgKchL-HA-HaloTag7-myc-PDGFRtmb were used. Doxcycline (500 ng $mL^{-1}$) was added during seeding to induce expression. Imaging was performed before and after titration with 2 M KCl stock to final concentrations of 10–200 mM. For intracellular MaPK imaging, U2OS cells stably expressing HaloTag7–P30–SNAP-tag2–P2A–NLS–mTurquoise2 were used. Cells were incubated with 4 µM digitonin for 10 min prior to imaging, and imaging were repeated following with KCl solutions across the same concentration gradient.

**rAAVs production.** rAAVs were produced as previously described[6]. Briefly, HEK293 Lenti-X cells were co-transfected with four plasmids: pRV1 (containing AAV2 Rep and Cap sequences), pH21 (containing AAV1 Rep and Cap sequences), pFdelta6 (adenovirus-helper plasmid) and a pAAV plasmid containing the recombinant expression cassette driven by the hSyn1 promoter and flanked by AAV2 inverted terminal repeats (ITRs). Transfection was carried out using Lipofectamine 3000 (Invitrogen, L3000001) according to the manufacturer's instructions. After 64 hours, both cells and culture medium were collected by centrifugation at 1,000 g for 5 min at 4°C. The cell pellet were lysed in TNT extraction buffer (20 mM Tris pH 7.5, 150 mM NaCl, 1% Triton X-100, 10 mM $MgCl_2$). Lysates were cleared by centrifugation at 2,000 g for 5 min at 4°C, and the resulting supernatant was treated with 300 U $mL^{-1}$ benzonase nuclease (Millipore, 70746-3) for 30–60 min at 37°C, with gentle mixing every 20 min. Viral particles from the combined medium and supernatant were purified using an Äkta-Quick FPLC system (Cytiva) with AVB Sepharose HiTrap columns (Cytiva, 2841211). Columns were equilibrated with PBS, and virus were eluted with 50 mM glycine-HCl (pH 2.7). The eluate was concentrated and buffer-exchanged into PBS using Amicon Ultra centrifugal filters (Millipore, UFC810024; 100 kDa MWCO). Aliquots of purified rAAVs were flash-frozen and stored at −80 °C until use.

**Primary rat hippocampal neurons preparation.** All procedures were conducted in strict accordance with the Animal Welfare Act of the Federal Republic of Germany (Tierschutzgesetz der Bundesrepublik Deutschland, TierSchG) and the Animal Welfare Laboratory Animal Regulations (Tierschutzversuchsverordnung). According to these regulations, no ethical approval from an ethics committee is required for euthanizing rodents when the organs or tissues are used for scientific purposes. The euthanasia procedure for rats in this study was supervised by animal welfare officers of the Max Planck Institute for Medical Research and was carried out and documented in compliance with the TierSchG (permit number assigned by the Max Planck Institute for Medical Research: MPI/T-35/18). Primary rat hippocampal neurons were prepared from hippocampi isolated from postnatal day 0–1 (P0–P1) Wistar rats of both sexes, following established protocols[6]. Neurons were seeded onto 24-well glass bottom imaging plates coated with poly-L-ornithine (100 μg $mL^{-1}$ in water) and laminin (1 μg $mL^{-1}$ in HBSS), and maintained in a humidified incubator at 37°C and 5% $CO_2$.

**rAAV transduction.** On day 6, one-third of the neuronal culture medium was refreshed. On day 7, neurons were transduced with rAAVs (serotype 2/1) at concentrations ranging from $10^9$ to $10^{10}$ genome copies per milliliter. Cultures were maintained for 7 days to allow transgene expression, with one-third of the medium replaced every three days during this period.

**Live cell imaging of MaPKs in primary neurons.** Neurons were seeded into 24-well imaging chambered coverslips with glass bottoms and maintained in culture until 14–16 days in vitro before imaging. Neurons were incubated with 1 μM MaPKs for 2 h at 37°C, followed by a medium exchange prior to imaging. For extracellular MaPK imaging, neurons were transduced with rAAV to drive hSyn1 promoter-mediated expression of IgKchL-HA-HaloTag7-myc-PDGFRtmb. Imaging was performed both before and after titration with 2 M KCl stock to final concentrations of 30 mM or 200 mM. For intracellular MaPK imaging, neurons were transduced with rAAV carrying hSyn1 promoter-driven NES-HaloTag7-mEGFP. Baseline frames were acquired every 45 s for five frames, followed by image acquisition after treating with 500 μM glutamate.

**Microscopy.** Imaging was performed using a commercial Leica Stellaris 5 confocal microscope equipped with a supercontinuum white light laser (470–670 nm) and hybrid photodetectors for single-molecule detection (HyD SMD). The laser power output was set to 85% of its maximum and regularly calibrated. The microscope stage was maintained in an environmental chamber at 37°C with 5% $CO_2$. Unless otherwise specified, the following imaging settings were used: HC PL APO CS2 ×20/0.75-NA (numerical aperture) air/water objective, field-of-view 581.82×581.82 μm, scan speed 400 MHz, Z-stacks acquired with a 2 μm step size, and a total stack depth of 12 μm.

**Imaging processing and analysis.** All images were processed and analyzed using ImageJ/Fiji (version 2.16.0/1.54p)[7]. Unless otherwise specified, 16-bit Z-stack images were converted to maximum intensity projections (MIP). Region of interest (ROIs) were manually delineated or segmented using Cellpose 3.0.7[8]. Mean fluorescence intensities for individual ROIs were quantified across multiple fields of view. Unhealthy neurons were excluded from the analyses. Background fluorescence was determined by average mean intensities from 1–3 cell-free regions and subtracted for correction.

**Data representation, reproducibility and statistical analysis.** Numerical data was analyzed and plotted using Microsoft Excel (version 16.78.3), Origin 2024 and GraphPad Prism (version 10.3.0). Schematics and figures were assembled in Adobe Illustrator 2025. For fluorescence image presentation, brightness and contrast were adjusted identically across all channels. Unless otherwise specified, all in vitro measurements were performed in three technical replicates, and all cell-based experiments were conducted in at least two biological replicates.

**Data availability.** All data are available in the paper or the supplementary materials. Plasmids of interest from the study have been deposited at Addgene (https://www.addgene.org/Kai_Johnsson/), Reagents and materials are available from the corresponding authors upon request.

### 5. Chemical synthesis and characterization

#### 5.1 Synthesis of sulfonamides

**General Procedure A (GP A):**

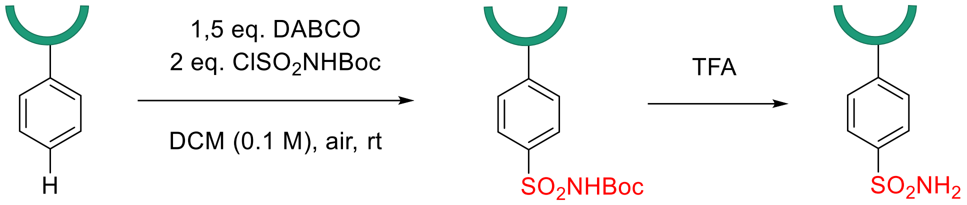

To a vial containing a solution of substrate (0.500 mmol, 1.0 equiv.) and 1,4-diazabicyclo[2.2.2]octane (DABCO) (84.1 mg, 0.750 mmol, 1.5 equiv.) in dichloromethane (2.5 mL) was added a solution of *N*-(*tert*-butoxycarbonyl)sulfamoyl chloride[9] ($ClSO_2NHBoc$) (216 mg, 1.00 mmol, 2.0 equiv.) in dichloromethane (2.5 mL) at room temperature under air. Upon addition of $ClSO_2NHBoc$, white precipitate was formed immediately. The suspension was kept stirring for 16 hours before being quenched by the addition of saturated $NaHCO_3$ (2 mL). The aqueous layer was then extracted with dichloromethane (3 x 10 mL). The organic phases were combined, dried over $Na_2SO_4$, filtered and concentrated *in vacuo*. The pure product was obtained after column chromatography on Biotage ($SiO_2$ or neutral $Al_2O_3$, 12 g, eluent with 10 - 70% ethyl acetate in n-Hexane, linear gradient).

**4-(1,4,7,10,13-Pentaoxa-16-azacyclooctadecan-16-yl)-3-(2-methoxyethoxy)benzenesulfonamide** (**1**)

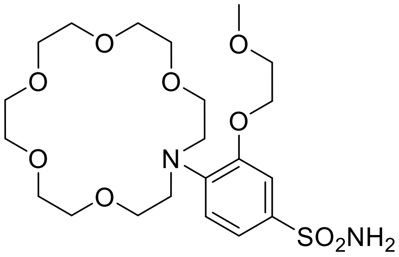

Following GP A, using 16-(2-(2-methoxyethoxy)phenyl)-1,4,7,10,13-pentaoxa-16-azacyclooctadecane (41.4 mg, 100 µmol), 4-(1,4,7,10,13-Pentaoxa-16-azacyclooctadecan-16-yl)-3-(2-methoxyethoxy)benzenesulfonamide **1** (22.8 mg, 46.2 µmol, 46%) was obtained as a beige oil.

**$^1$H NMR** (400 MHz, $CDCl_3$): δ = 7.88 (d, J = 1.8 Hz, 1H, ArH), 7.80 (d, J = 8.3 Hz, 1H, ArH), 7.68 (dd, J = 8.4, 1.7 Hz, 1H, ArH), 4.42 (t, J = 4.1 Hz, 2H, OC*H*2), 3.87 (s, 4H, OC*H*2), 3.78 – 3.57 (m, 15H, OC*H*2), 3.47 – 3.38 (m, 7H, OC*H*2 + NC*H*2), 3.36 (s, 3H, OC*H*3);

$^1$H NMR data correspond to the reported values[9].

**4-(1,4,7,10-Tetraoxa-13-azacyclopentadecan-13-yl)-3-methoxybenzenesulfonamide** (**2**)

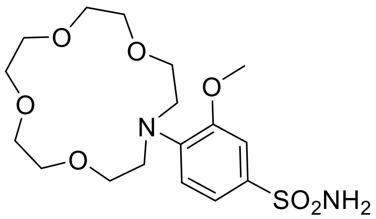

Following GP A, to a vial containing a solution of 13-(2-methoxyphenyl)-1,4,7,10-tetraoxa-13-azacyclopentadecane (65.1 mg, 200 μmol) and DABCO (33.7 mg, 300 μmol, 1.5 equiv.) in dichloromethane (1.0 mL) was added a solution of $ClSO_2NHBoc$ (86.3 mg, 400 μmol, 2.0 equiv.) in dichloromethane (1.0 mL) at room temperature under air for 16 hours. TFA (0.15 mL) was added to the reaction crude and the mixture was kept stirring for 2 hours. The volatiles were removed under vacuum and the crude was purified by column chromatography on Biotage (neutral $Al_2O_3$ 4 g, eluent with 0 – 15% MeOH in DCM, linear gradient) and fractions containing the corresponding product were combined. 4-(1,4,7,10-Tetraoxa-13-azacyclopentadecan-13-yl)-3-methoxybenzenesulfonamide **2** (76.0 mg, 0.188 mmol, 94%) was obtained as a pale yellow oil.

**$^1$H NMR** (400 MHz, $CDCl_3$): δ = 7.66 (s, 1H, ArH), 7.61 – 7.56 (m, 1H, ArH), 7.54 (dd, *J* = 8.3, 1.9 Hz, 1H, ArH), 3.98 (s, 3H, OC*H*3), 3.88 – 3.74 (m, 4H, OC*H*2), 3.69 (s, 4H, OC*H*2), 3.64 (dt, *J* = 5.3, 1.8 Hz, 4H, OC*H*2), 3.59 (p, *J* = 2.6, 2.2 Hz, 8H, OC*H*2 + NC*H*2);
**$^{13}$C NMR** (101 MHz, $CDCl_3$): δ = 151.9, 123.5, 119.8, 117.4, 114.5, 111.0, 70.3, 69.7, 69.4, 65.9, 56.6, 55.6;
**HRMS** (ESI) calcd. for $C_{17}H_{29}N_2O_7S$ $[M+H]^+$ 405.1690; Found 405.1691.

**((4-(*N*-(*tert*-Butoxycarbonyl)sulfamoyl)phenyl)azanediyl)bis(ethane-2,1-diyl) bis(4-methylbenzenesulfonate)** (**9**)

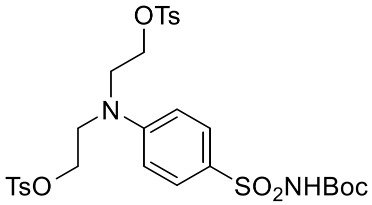

Following GP A, to a flask containing a solution of (phenylazanediyl)bis(ethane-2,1-diyl) bis(4-methylbenzenesulfonate) (1.06 g, 2.16 mmol) and DABCO (0.364 mg, 3.25 mmol, 1.5 equiv.) in dichloromethane (10 mL) was added a solution of $ClSO_2NHBoc$ (0.934 mg, 4.33 mmol, 2.0 equiv.) in dichloromethane (10 mL) at room temperature under air for 16 hours. The reaction was subsequently quenched by water (20 mL), extracted with dichloromethane (10 mL x 3). The organic phases were combined and dried over $Na_2SO_4$. The crude product was purified by recrystallization in dichloromethane. ((4-(*N*-(*tert*-Butoxycarbonyl)sulfamoyl)phenyl)azanediyl)bis(ethane-2,1-diyl) bis(4-methylbenzenesulfonate) **9** (1.34 g, 2.00 μmol, 93%) was obtained as a white solid.

**$^1$H NMR** (400 MHz, $CDCl_3$): δ = 7.76 – 7.71 (m, 2H), 7.70 – 7.65 (m, 4H), 7.28 (s, 4H), 7.13 (s, 1H), 6.48 – 6.41 (m, 2H), 4.13 (t, *J* = 5.7 Hz, 4H), 3.63 (t, *J* = 5.8 Hz, 4H), 2.43 (s, 6H), 1.41 (s, 9H);

**$^{13}$C NMR** (101 MHz, $CDCl_3$): $\delta$ = 149.9, 149.1, 145.4, 132.3, 130.4, 130.0, 127.8, 125.9, 110.7, 83.8, 66.0, 50.0, 28.0, 27.8, 21.6;
**HRMS** (ESI) calcd. for $C_{29}H_{37}N_2O_{10}S_3$ $[M+H]^+$ 669.1605; Found 669.1609.

***tert*-Butyl ((4-(bis(2-((2-(ethylthio)ethyl)thio)ethyl)amino)phenyl)sulfonyl)carbamate** (**3**)

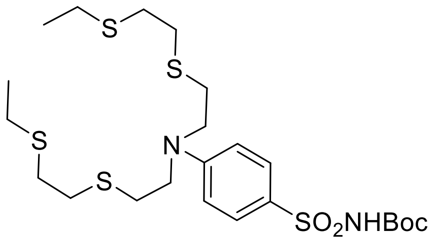

To a vial containing a solution of **9** (66.9 mg, 100 µmol) and $Cs_2CO_3$ (71.7 mg, 220 µmol, 2.2 equiv.) in DMF (0.5 mL) was added a solution of 2-(ethylsulfanyl)ethanethiol (26.9 mg, 220 µmol, 2.2 equiv.) in DMF (0.5 mL) at room temperature under air. After 16 hours, the crude was purified by column chromatography on Biotage ($SiO_2$ 4 g, eluent with 0 - 80% ethyl acetate in n-Hexane, linear gradient) and fractions containing the corresponding product were combined. *tert*-Butyl ((4-(bis(2-((2-(ethylthio)ethyl)thio)ethyl)amino)phenyl)sulfonyl)carbamate **3** (42.7 mg, 75.1 µmol, 75%) was obtained as a colorless oil.

**$^1$H NMR** (400 MHz, $CDCl_3$): $\delta$ = 7.88 – 7.79 (m, 2H), 7.11 (s, 1H), 6.72 – 6.63 (m, 2H), 3.69 – 3.59 (m, 4H), 2.76 (dddd, *J* = 12.6, 8.6, 5.0, 2.4 Hz, 12H), 2.57 (q, *J* = 7.4 Hz, 4H), 1.42 (s, 9H), 1.27 (t, *J* = 7.4 Hz, 6H);
**$^{13}$C NMR** (101 MHz, $CDCl_3$): $\delta$ = 150.5, 149.2, 130.6, 124.9, 110.6, 83.6, 51.5, 32.6, 31.8, 29.3, 28.0, 26.2, 14.8;
**HRMS** (ESI) calcd. for $C_{23}H_{41}N_2O_4S_5$ $[M+H]^+$ 569.1664; Found 569.1665.

**Dimethyl 2,2'-((((4-(*N*-(*tert*-butoxycarbonyl)sulfamoyl)phenyl)azanediyl)bis(ethane-2,1-diyl))bis(sulfanediyl))diacetate** (**4**)

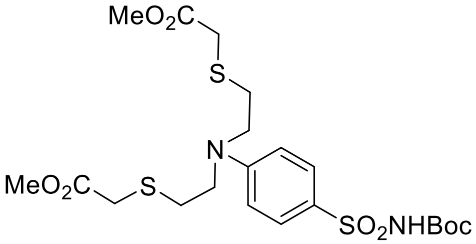

To a vial containing a solution of **9** (66.9 mg, 100 µmol) and $Cs_2CO_3$ (71.7 mg, 220 µmol, 2.2 equiv.) in DMF (0.5 mL) was added a solution of methyl thioglycolate (23.4 mg, 220 µmol, 2.2 equiv.) in DMF (0.5 mL) at room temperature under air. After 16 hours, the crude was purified by column chromatography on Biotage ($SiO_2$ 4 g, eluent with 0 - 80% ethyl acetate in n-Hexane, linear gradient) and fractions containing the corresponding product were combined. Dimethyl 2,2'-((((4-(*N*-(*tert*-butoxycarbonyl)sulfamoyl)phenyl)azanediyl)bis(ethane-2,1-diyl))bis(sulfanediyl))diacetate **4** (42.5 mg, 79.2 µmol, 79%) was obtained as a colorless oil.

**$^{1}$H NMR** (400 MHz, $CDCl_3$): δ = 7.87 – 7.78 (m, 2H), 7.27 (d, *J* = 2.4 Hz, 1H), 6.77 – 6.68 (m, 2H), 3.75 (s, 6H), 3.70 – 3.60 (m, 4H), 3.28 (s, 4H), 2.90 – 2.81 (m, 4H), 1.40 (s, 9H);
**$^{13}$C NMR** (101 MHz, $CDCl_3$): δ = 170.6, 150.6, 149.3, 130.5, 125.0, 110.6, 83.5, 52.6, 50.6, 33.3, 29.3, 27.9;
**HRMS** (ESI) calcd. for $C_{21}H_{33}N_2O_8S_3$ $[M+H]^+$ 537.1394; Found 537.1397.

**Bis(acetoxymethyl) 2,2'-((2-(2-(2-(bis(2-(acetoxymethoxy)-2-oxoethyl)amino)-5-sulfamoylphenoxy)ethoxy)phenyl)azanediyl)diacetate** (**5**)

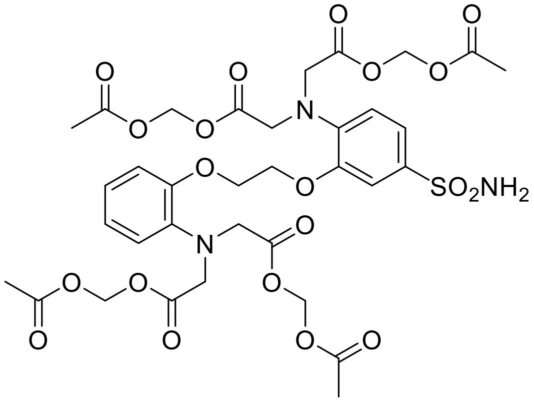

Following GP A, using BAPTA-AM (38.2 mg, 50.0 µmol), bis(acetoxymethyl) 2,2'-((2-(2-(2-(bis(2-(acetoxymethoxy)-2-oxoethyl)amino)-5-sulfamoylphenoxy)ethoxy)phenyl)azanediyl)diacetate **5** (22.4 mg, 26.6 µmol, 53%) was obtained as a brown oil.

**$^{1}$H NMR** (400 MHz, $CDCl_3$): δ = 7.46 – 7.41 (m, 2H, Ar*H*), 7.09 (td, *J* = 7.7, 1.6 Hz, 1H, Ar*H*), 7.03 (dd, *J* = 8.0, 1.6 Hz, 1H, Ar*H*), 6.99 – 6.92 (m, 2H, Ar*H*), 6.81 (d, *J* = 8.9 Hz, 1H, Ar*H*), 5.62 (s, 4H, OC*H*2O), 5.60 (s, 4H, OC*H*2O), 4.41 – 4.36 (m, 2H, OC*H*2), 4.36 – 4.31 (m, 2H, OC*H*2), 4.28 (s, 4H, NC*H*2), 4.21 (s, 4H, NC*H*2), 2.09 (s, 6H, C*H*3), 2.07 (s, 6H, C*H*3);

$^{1}$H NMR data correspond to the reported values[9].

**Diethyl 2,2'-((2-(2-ethoxy-2-oxoethoxy)-4-sulfamoylphenyl)azanediyl)diacetate** (**6**)

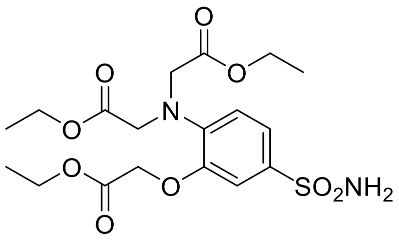

Following GP A, to a vial containing a solution of diethyl 2,2'-((2-(2-ethoxy-2-oxoethoxy)phenyl)azanediyl)diacetate (0.367 g, 1.00 mmol) and DABCO (0.168 g, 1.50 mmol, 1.5 equiv.) in dichloromethane (5.0 mL) was added a solution of $ClSO_2NHBoc$ (0.432 g, 2.00 mmol, 2.0 equiv.) in dichloromethane (5.0 mL) at room temperature under air for 16 hours. After completion, the crude was treated with TFA (1.5 mL) for 2 hours before being purified by column chromatography on Biotage ($SiO_2$ 12 g, eluent with 10 - 70% ethyl acetate in n-Hexane, linear gradient) and fractions containing the corresponding product were combined. Diethyl 2,2'-((2-(2-ethoxy-2-oxoethoxy)-4-sulfamoylphenyl)azanediyl)diacetate **6** (0.201 g, 0.450 mmol, 45%) was obtained as a beige solid.

**R$_f$:** 0.28 (silica, pentanes:ethyl acetate 1:1);
**$^1$H NMR** (400 MHz, $CDCl_3$): δ = 7.41 (dd, *J* = 8.4, 2.0 Hz, 1H, Ar*H*), 7.27 (d, *J* = 2.1 Hz, 1H, Ar*H*), 6.86 (d, *J* = 8.4 Hz, 1H, Ar*H*), 5.36 (s, 2H, N*H2*), 4.28 (q, *J* = 7.1 Hz, 2H, OC*H*2), 4.25 – 4.16 (m, 10H, OC*H*2 + NC*H*2), 1.32 (t, *J* = 7.2 Hz, 3H, C*H*3), 1.26 (t, *J* = 7.1 Hz, 6H, C*H*3);
**$^{13}$C NMR** (101 MHz, $CDCl_3$): δ = 170.7, 168.1, 148.6, 142.2, 135.4, 120.4, 118.1, 111.4, 65.4, 61.8, 61.2, 53.9, 14.1, 14.1;
**HRMS** (ESI) calcd. for $C_{18}H_{27}N_2O_9S$ $[M+H]^+$ 447.1432; Found 447.1421.

**Diethyl 2,2'-((2-methoxy-4-sulfamoylphenyl)azanediyl)diacetate** (**7**)

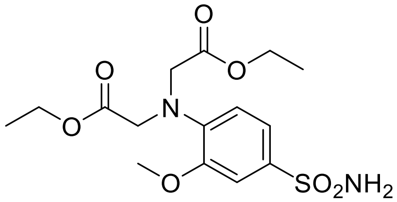

Following GP A, to a vial containing a solution of diethyl 2,2'-((2-methoxyphenyl)azanediyl)diacetate (0.250 g, 0.85 mmol) and DABCO (0.143 g, 1.27 mmol, 1.5 equiv.) in dichloromethane (3.0 mL) was added a solution of $ClSO_2NHBoc$ (0.367 g, 1.70 mmol, 2.0 equiv.) in dichloromethane (3.0 mL) at room temperature under air for 16 hours. After completion, the crude was treated with TFA (1.5 mL) for 2 hours before being purified by column chromatography on Biotage ($SiO_2$ 12 g, eluent with 10 - 70% ethyl acetate in n-Hexane, linear gradient) and fractions containing the corresponding product were combined. Diethyl 2,2'-((2-methoxy-4-sulfamoylphenyl)azanediyl)diacetate **7** (0.105 g, 0.280 mmol, 33%) was obtained as a pale yellow oil.

**R$_f$:** 0.20 (silica, pentanes:ethyl acetate 1:1);
**$^1$H NMR** (400 MHz, $CDCl_3$): δ = 7.42 (dd, J = 8.5, 2.1 Hz, 1H, Ar*H*), 7.34 (d, J = 2.1 Hz, 1H, Ar*H*), 6.75 (d, J = 8.5 Hz, 1H, Ar*H*), 4.82 (s, 2H, N*H2*), 4.21 (q, J = 7.1 Hz, 4H, OC*H*2), 4.14 (s, 4H, NC*H*2), 3.82 (s, 3H, OC*H*3), 1.28 (t, J = 7.1 Hz, 6H, C*H*3);
**$^{13}$C NMR** (101 MHz, $CDCl_3$): δ = 170.8, 150.3, 143.0, 133.6, 120.1, 117.1, 110.0, 61.0, 56.0, 54.1, 14.2;
**HRMS** (ESI) calcd. for $C_{15}H_{23}N_2O_7S$ $[M+H]^+$ 375.1220; Found 375.1224.

***tert*-Butyl (N-(1,10-phenanthrolin-5-yl)sulfamoyl)carbamate** (**8**)

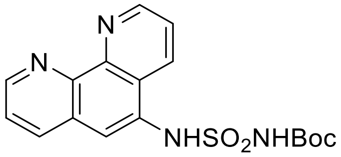

Following GP A, to a vial containing a solution of 5-amino-1,10-phenanthroline (39.0 mg, 200 μmol) and DABCO (33.6 mg, 300 μmol, 1.5 equiv.) in dichloromethane (1.0 mL) was added a solution of $ClSO_2NHBoc$ (86.4 mg, 400 µmol, 2.0 equiv.) in dichloromethane (1.0 mL) at room temperature under air. After 16 hours, the crude was purified by column chromatography on Biotage ($SiO_2$ 12 g, eluent with 0 - 10% methanol in dichloromethane, linear gradient) and fractions containing the corresponding product were combined. *tert*-Butyl (N-(1,10-phenanthrolin-5-yl)sulfamoyl)carbamate **8** (42.0 mg, 112 µmol, 56%) was obtained as a yellow solid.

**$^1$H NMR** (400 MHz, $CDCl_3$): δ = 9.11 (dt, *J* = 3.7, 1.8 Hz, 2H), 8.76 (dd, *J* = 8.4, 1.6 Hz, 1H), 8.19 (dd, *J* = 8.2, 1.7 Hz, 1H), 7.92 (s, 1H), 7.63 (ddd, *J* = 8.1, 4.4, 1.5 Hz, 2H), 1.39 (s, 9H);
**$^{13}$C NMR** (101 MHz, $CDCl_3$): δ = 150.6, 150.6, 150.2, 146.2, 144.9, 136.2, 131.9, 129.9, 127.8, 125.6, 123.6, 123.2, 122.5, 84.4, 27.9;
**HRMS** (ESI) calcd. for $C_{17}H_{19}N_4O_4S$ $[M+H]^+$ 375.1122; Found 375.1121.

#### 5.2 Synthesis of CA-MaP-558

**General Procedure B (GP B):**

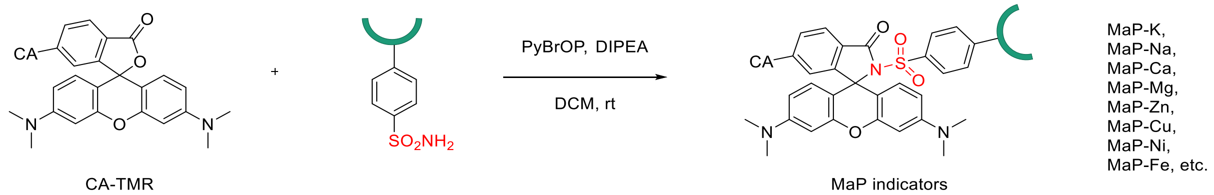

To a vial containing a solution of CA-TMR (5.0 mg, 7.8 µmol, 1.0 equiv.) and the corresponding sulfonamide (9.4 µmol, 1.2 equiv.; if the sulfonamide is in its Boc protected form, it was first treated with 0.1 mL TFA for 30 minutes and then concentrated) in dichloromethane (0.5 mL) was added bromo-tris-pyrrolidino-phosphonium hexafluorophosphate (PyBrOP) (5.5 mg, 11.8 µmol, 1.5 equiv.) and DIPEA (5.2 µL, 31.4 µmol, 4.0 equiv.) at room temperature under air. After stirring overnight, the reaction mixture was filtered and purified using preparative HPLC (8 mL/min, 30 - 90 % MeCN/$H_2O$ (0.1 % TFA) in 60 min) to give the corresponding $MaP_{Halo}$-558 sensor as a red solid.

**2-((4-(1,4,7,10,13-Pentaoxa-16-azacyclooctadecan-16-yl)-3-(2-methoxyethoxy)phenyl)sulfonyl)-N-(2-(2-((6-chlorohexyl)oxy)ethoxy)ethyl)-3',6'-bis(dimethylamino)-3-oxospiro[isoindoline-1,9'-xanthene]-6-carboxamide (CA-MaPK-558)**

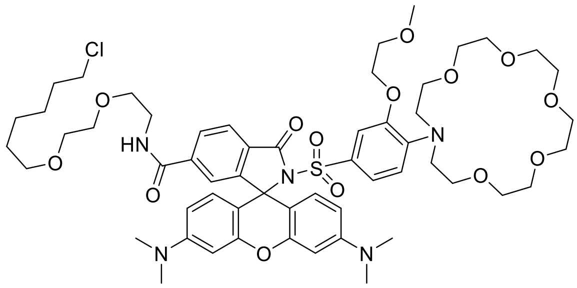

Following GP B, using **1** (4.6 mg, 9.4 µmol, 1.2 eq), **CA-MaPK-558** (2.6 mg, 2.9 µmol, 37%) was obtained as a red solid.

**$^{1}$H NMR** (400 MHz, $CD_3OD$): δ = 8.09 (dd, *J* = 8.1, 1.5 Hz, 1H), 8.00 (d, *J* = 8.1 Hz, 1H), 7.58 (s, 1H), 6.86 (s, 1H), 6.76 – 6.63 (m, 2H), 6.54 (s, 3H), 3.98 (s, 2H), 3.84 – 3.43 (m, 33H), 3.42 – 3.33 (m, 8H), 3.10 (s, 12H), 1.75 – 1.65 (m, 2H), 1.51 – 1.44 (m, 2H), 1.43 – 1.35 (m, 3H), 1.30 (tt, *J* = 9.3, 7.4, 3.0 Hz, 3H);
**HRMS** (ESI) calcd. for $C_{56}H_{78}N_5O_{14}ClS$ $[M+2H]^{2+}$ 555.7472; Found 555.7479.

**2-((4-(1,4,7,10-tetraoxa-13-azacyclopentadecan-13-yl)-3-methoxyphenyl)sulfonyl)-N-(2-(2-((6-chlorohexyl)oxy)ethoxy)ethyl)-3',6'-bis(dimethylamino)-3-oxospiro[isoindoline-1,9'-xanthene]-6-carboxamide (CA-MaPNa-558)**

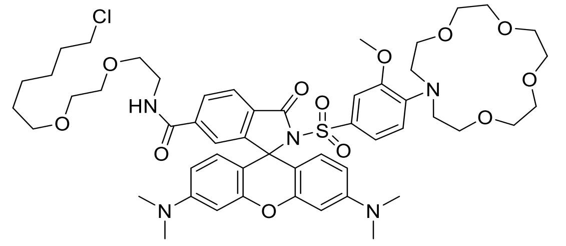

Following GP B, using **2** (3.8 mg, 9.4 µmol, 1.2 eq), **CA-MaPNa-558** (1.0 mg, 0.98 µmol, 12%) was obtained as a red solid.

**$^{1}$H NMR** (400 MHz, $CD_3OD$): δ = 8.71 (s, 1H), 8.16 (d, *J* = 8.3 Hz, 1H), 8.00 (d, *J* = 8.1 Hz, 1H), 7.74 (d, *J* = 26.4 Hz, 1H), 7.23 – 7.16 (m, 1H), 7.15 – 6.44 (m, 8H), 3.84 – 3.33 (m, 35H), 3.32 – 3.01 (m, 12H), 1.77 – 1.66 (m, 2H), 1.50 (p, *J* = 6.8 Hz, 2H), 1.41 (p, *J* = 6.9 Hz, 2H), 1.36 – 1.28 (m, 2H);
**HRMS** (ESI) calcd. for $C_{52}H_{70}N_5O_{12}ClS$ $[M+2H]^{2+}$ 511.7210; Found 511.7208.

**2-((4-(bis(2-((2-(ethylthio)ethyl)thio)ethyl)amino)phenyl)sulfonyl)-N-(2-(2-((6-chlorohexyl)oxy)ethoxy)ethyl)-3',6'-bis(dimethylamino)-3-oxospiro[isoindoline-1,9'-xanthene]-6-carboxamide (CA-MaPCu-558)**

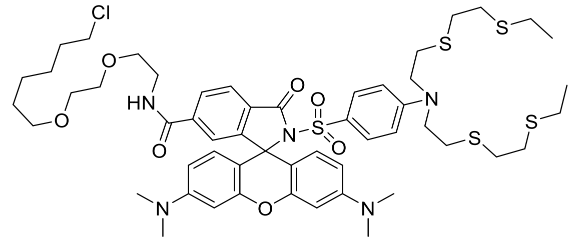

Following GP B, using **3** (5.4 mg, 9.4 µmol, 1.2 eq), **CA-MaPCu-558** (3.2 mg, 2.9µmol, 37%) was obtained as a red solid.

**$^{1}$H NMR** (400 MHz, DMSO-d6): δ = 8.68 (t, *J* = 5.6 Hz, 1H), 8.03 (dd, *J* = 8.1, 1.4 Hz, 1H), 7.89 (d, *J* = 8.0 Hz, 1H), 7.45 (d, *J* = 1.4 Hz, 1H), 7.05 (d, *J* = 8.9 Hz, 2H), 6.58 – 6.47 (m, 4H), 6.36 (dd, *J* = 8.9, 2.5 Hz, 2H), 6.29 (d, *J* = 8.8 Hz, 2H), 3.57 (td, *J* = 7.8, 7.2, 3.7 Hz, 6H), 3.44 (d, *J* = 5.7 Hz, 6H), 3.29 (q, *J* = 6.5, 5.9 Hz, 4H), 2.96 (s, 12H), 2.81 – 2.63 (m, 12H), 2.54 (d, *J* = 7.4 Hz, 4H), 1.69 – 1.60 (m, 2H), 1.39 (p, *J* = 7.0 Hz, 2H), 1.34 – 1.26 (m, 2H), 1.26 – 1.19 (m, 2H), 1.15 (t, *J* = 7.4 Hz, 6H);

**$^{13}$C NMR** (101 MHz, DMSO-d6): δ = 164.7, 164.6, 157.0, 153.0, 152.4, 151.3, 150.6, 140.2, 130.1, 129.7, 128.4, 124.0, 123.5, 122.8, 109.8, 108.5, 106.5, 98.3, 70.1, 69.5, 69.3, 68.6, 68.3, 50.5, 45.3, 44.4, 32.0, 31.5, 31.1, 29.0, 28.0, 26.1, 24.9, 24.9, 14.7;
**HRMS** (ESI) calcd. for $C_{53}H_{73}N_5O_7ClS_5$ $[M+H]^+$ 1086.3797; Found 1086.3798.

**Dimethyl 2,2'-((((4-((6-((2-(2-((6-chlorohexyl)oxy)ethoxy)ethyl)carbamoyl)-3',6'-bis(dimethylamino)-3-oxospiro[isoindoline-1,9'-xanthen]-2-yl)sulfonyl)phenyl)azanediyl)bis(ethane-2,1-diyl))bis(sulfanediyl))diacetate (CA-MaPNi-558)**

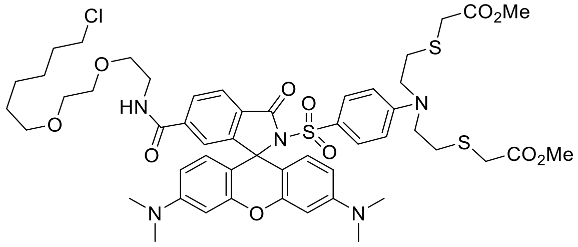

Following GP B, using **4** (5,1 mg, 9.4 µmol, 1.2 eq), **CA-MaPNi-558** (1.8 mg, 1.7 µmol, 22%) was obtained as a red solid.

**$^{1}$H NMR** (400 MHz, DMSO-d6): δ = 8.69 (t, *J* = 5.7 Hz, 1H), 8.03 (dd, *J* = 8.1, 1.5 Hz, 1H), 7.90 (d, *J* = 8.0 Hz, 1H), 7.46 (d, *J* = 1.4 Hz, 1H), 7.05 (d, *J* = 8.8 Hz, 2H), 6.58 (d, *J* = 9.0 Hz, 2H), 6.49 (d, *J* = 2.5 Hz, 2H), 6.41 – 6.27 (m, 4H), 3.64 (s, 6H), 3.57 (t, *J* = 6.6 Hz, 10H), 3.45 – 3.38 (m, 6H), 3.29 (dt, *J* = 12.7, 5.9 Hz, 4H), 2.95 (s, 12H), 2.74 (t, *J* = 7.6 Hz, 4H), 1.65 (p, *J* = 6.8 Hz, 2H), 1.39 (p, *J* = 7.0 Hz, 2H), 1.35 – 1.28 (m, 2H), 1.26 – 1.21 (m, 2H);
**$^{13}$C NMR** (101 MHz, DMSO-d6): δ = 170.8, 164.7, 164.6, 153.1, 152.4, 151.3, 150.6, 140.2, 130.1, 129.7, 128.4, 124.0, 123.5, 122.8, 109.8, 108.5, 106.5, 98.3, 70.1, 69.5, 69.3, 68.6, 68.3, 52.1, 49.7, 45.3, 40.2, 39.8, 32.5, 32.0, 29.0, 28.4, 26.1, 24.9;
**HRMS** (ESI) calcd. for $C_{51}H_{65}N_5O_{11}ClS_3$ $[M+H]^+$ 1054.3526; Found 1054.3534.

**Bis(acetoxymethyl) 2,2'-((2-(2-(2-(bis(2-(acetoxymethoxy)-2-oxoethyl)amino)-5-((6-((2-(2-((6-chlorohexyl)oxy)ethoxy)ethyl)carbamoyl)-3',6'-bis(dimethylamino)-3-oxospiro[isoindoline-1,9'-xanthen]-2-yl)sulfonyl)phenoxy)ethoxy)phenyl)azanediyl)diacetate (CA-MaPCa2-558)**

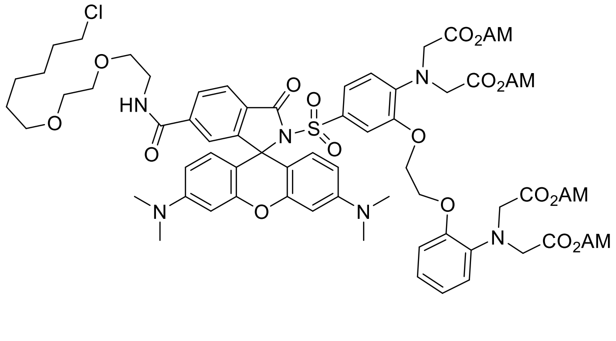

Following GP B, using **5** (8.0 mg, 9.4 µmol, 1.2 eq), **CA-MaPCa2-558** (2.6 mg, 1.7 µmol, 22%) was obtained as a red solid.

**$^{1}$H NMR** (400 MHz, $CD_3OD$): δ = 8.69 (t, *J* = 5.7 Hz, 1H), 8.04 (d, *J* = 8.2 Hz, 1H), 7.91 (d, *J* = 8.0 Hz, 1H), 7.46 (s, 1H), 7.03 (dd, *J* = 8.4, 2.1 Hz, 1H), 6.99 – 6.94 (m, 1H), 6.94 – 6.89 (m, 1H), 6.86 (t, *J* = 7.2 Hz, 1H), 6.76 (dd, *J* = 7.8, 1.8 Hz, 1H), 6.61 (d, *J* = 8.6 Hz, 2H), 6.48 (s, 2H), 6.34 (d, *J* = 2.3 Hz, 3H), 5.53 (d, *J* = 4.2 Hz, 8H), 4.18 (d, *J* = 36.1 Hz, 12H), 3.39 (dd, *J* = 5.9, 3.6 Hz, 3H), 3.29 (q, *J* = 6.4, 5.7 Hz, 5H), 2.93 (s, 12H), 2.00 (d, *J* = 9.0 Hz, 12H), 1.65 (p, *J* = 6.8 Hz, 2H), 1.40 (t, *J* = 7.2 Hz, 2H), 1.31 (q, *J* = 7.5, 6.9 Hz, 2H), 1.26 – 1.20 (m, 2H);
**HRMS** (ESI) calcd. for $C_{69}H_{83}N_6O_{25}ClS$ $[M+2H]^{2+}$ 731.2403; Found 731.2392.

**Diethyl 2,2'-((4-((6-((2-(2-((6-chlorohexyl)oxy)ethoxy)ethyl)carbamoyl)-3',6'-bis(dimethylamino)-3-oxospiro[isoindoline-1,9'-xanthen]-2-yl)sulfonyl)-2-(2-ethoxy-2-oxoethoxy)phenyl)azanediyl)diacetate (CA-MaPMg-558)**

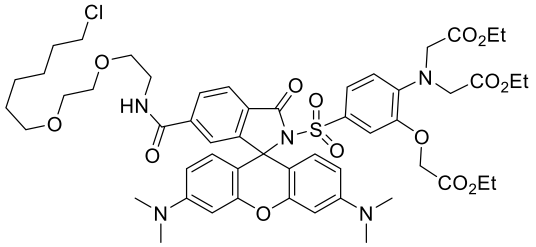

Following GP B, using **6** (4.2 mg, 9.4 µmol, 1.2 eq), **CA-MaPMg-558** (3.7 mg, 3.5 µmol, 45%) was obtained as a red solid.

**$^{1}$H NMR** (400 MHz, DMSO-d6): δ = 8.69 (t, *J* = 5.6 Hz, 1H), 8.03 (dd, *J* = 8.1, 1.5 Hz, 1H), 7.91 (d, *J* = 8.0 Hz, 1H), 7.44 (d, *J* = 1.4 Hz, 1H), 7.09 (dd, *J* = 8.7, 2.1 Hz, 1H), 6.63 (d, *J* = 8.8 Hz, 1H), 6.48 (d, *J* = 2.2 Hz, 2H), 6.43 (d, *J* = 2.1 Hz, 1H), 6.38 – 6.23 (m, 4H), 4.43 (s, 2H), 4.23 (s, 4H), 4.18 (q, *J* = 7.1 Hz, 2H), 4.08 (q, *J* = 7.1 Hz, 4H), 3.58 (t, *J* = 6.6 Hz, 4H), 3.47 – 3.37 (m, 8H), 3.33 – 3.25 (m, 4H), 2.95 (s, 12H), 1.65 (dt, *J* = 14.6, 6.7 Hz, 2H), 1.39 (p, *J* = 7.0 Hz, 2H), 1.35 – 1.27 (m, 2H), 1.23 (t, *J* = 7.1 Hz, 5H), 1.16 (t, *J* = 7.1 Hz, 6H);
**$^{13}$C NMR** (101 MHz, DMSO-d6): δ = 170.5, 168.1, 165.2, 165.0, 153.5, 152.9, 151.6, 146.6, 144.0, 140.8, 129.9, 129.1, 128.9, 124.1, 123.3, 116.5, 112.2, 108.7, 106.7, 98.6, 70.5, 70.0, 69.8, 69.1, 65.5, 61.4, 60.8, 54.0, 45.8, 40.7, 40.2,32.4, 29.4, 26.5, 25.3, 14.5, 14.4;
**HRMS** (ESI) calcd. for $C_{53}H_{67}N_5O_{14}ClS$ $[M+H]^{+}$ 1064.4088; Found 1064.4085.

**Diethyl 2,2'-((4-((6-((2-(2-((6-chlorohexyl)oxy)ethoxy)ethyl)carbamoyl)-3',6'-bis(dimethylamino)-3-oxospiro[isoindoline-1,9'-xanthen]-2-yl)sulfonyl)-2-methoxyphenyl)azanediyl)diacetate (CA-MaPZn-558)**

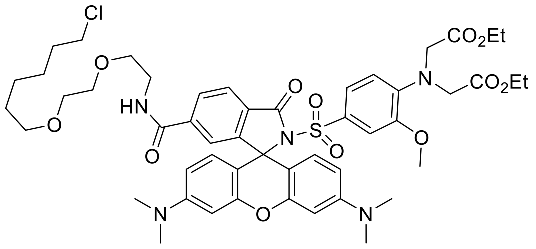

Following GP B, using **7** (3.5 mg, 9.4 µmol, 1.2 eq), **CA-MaPZn-558** (3.1 mg, 3.1 µmol, 40%) was obtained as a red solid.

**$^1$H NMR** (400 MHz, DMSO-d6): δ = 8.69 (t, *J* = 5.7 Hz, 1H), 8.04 (dd, *J* = 8.1, 1.4 Hz, 1H), 7.92 (d, *J* = 8.0 Hz, 1H), 7.46 (d, *J* = 1.5 Hz, 1H), 7.07 (dd, *J* = 8.7, 2.2 Hz, 1H), 6.54 (d, *J* = 8.8 Hz, 1H), 6.48 (dd, *J* = 7.6, 2.3 Hz, 3H), 6.37 – 6.26 (m, 4H), 4.11 (q, *J* = 7.3 Hz, 9H), 3.48 (s, 3H), 3.46 – 3.37 (m, 8H), 3.29 (dt, *J* = 12.8, 6.0 Hz, 4H), 2.94 (s, 12H), 1.65 (dt, *J* = 14.6, 6.8 Hz, 2H), 1.39 (p, *J* = 7.0 Hz, 2H), 1.35 – 1.28 (m, 2H), 1.27 – 1.21 (m, 2H), 1.19 (t, *J* = 7.1 Hz, 6H);
**$^{13}$C NMR** (101 MHz, DMSO-d6): δ = 170.2, 164.8, 164.5, 153.0, 152.5, 151.2, 148.1, 143.6, 140.3, 129.6, 128.6, 128.4, 123.7, 122.8, 122.4, 114.9, 111.1, 108.4, 106.3, 98.2, 70.1, 69.5, 69.3, 68.6, 60.3, 55.7, 53.9, 45.3, 40.2, 39.8, 32.0, 29.0, 26.1, 24.9, 14.1;
**HRMS** (ESI) calcd. for $C_{50}H_{63}N_5O_{12}ClS$ $[M+H]^+$ 992.3877; Found 992.3876.

**2-(N-(1,10-phenanthrolin-5-yl)sulfamoyl)-N-(2-(2-((6-chlorohexyl)oxy)ethoxy)ethyl)-3',6'-bis(dimethylamino)-3-oxospiro[isoindoline-1,9'-xanthene]-6-carboxamide (CA-MaPFe-558)**

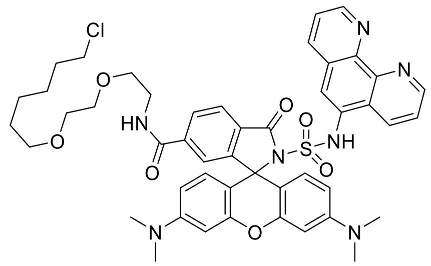

Following GP B, using **8** (2.6 mg, 9.4 µmol, 1.2 eq), **CA-MaPFe-558** (1.0 mg, 1,1 µmol, 14%) was obtained as a red solid.

**HRMS** (ESI) calcd. for $C_{47}H_{52}N_7O_7ClS$ $[M+2H]^{2+}$ 446.6663; Found 446.6666.

#### 5.3 Synthesis of CA-MaP-619, CA-MaP-656, TF-MaP-558

##### General Procedure C (GP C):

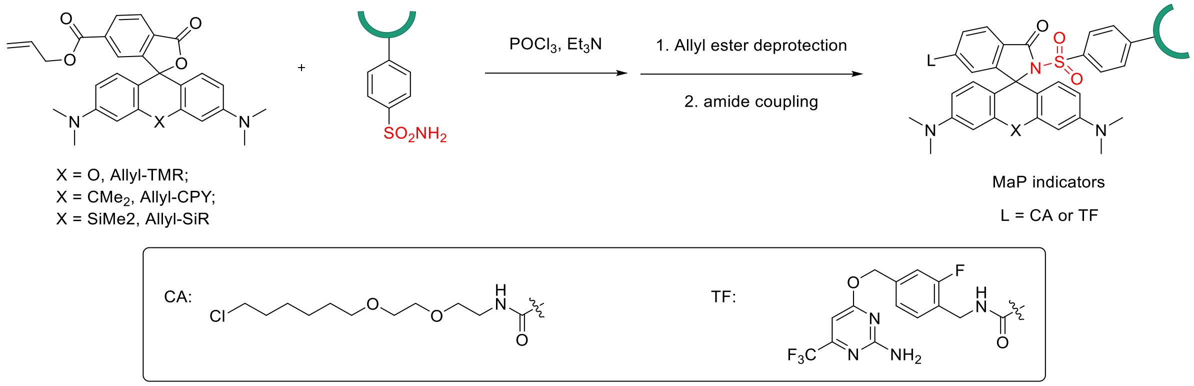

To a vial containing a solution of allyl-TMR (4.7 mg, 10 µmol, 1.0 equiv.) or allyl-CPY (5.0 mg, 10 µmol, 1.0 equiv.) or allyl-SiR (5.1 mg, 10 µmol, 1.0 equiv.) in dichloromethane (0.5 mL) was added phosphorus oxychloride (14.0 µL, 151 µmol, 15 equiv.). After stirring at 50 °C for 2 hours, a solution of DIPEA (49.8 µL, 301 µmol, 30 equiv.) and sulfonamide **1** (9.8 mg, 20 µmol, 2.0 equiv.) or sulfonamide **2** (8.0 mg, 20 µmol, 2.0 equiv.) in Acetonitrile (0.5 mL) was added and the reaction mixture was kept stirring at rt for overnight. The reaction was quenched with water (10 mL) and extracted with dichloromethane (10 mL x 3). The organic phases were combined, dried over $Na_2SO_4$ and concentrated. This residue was dissolved in MeOH/CH2Cl2 (5:1, 1.8 mL) in a vial, to which 1,3-Dimethylbarbituric acid (4.7 mg, 30.1 µmol, 3.0 eq) and tetrakis(triphenylphosphine)palladium (3.5 mg, 3.0 µmol, 0.3 eq) were added. After stirring at rt for 30 min, the solvent was evaporated, the crude product was dissolved in DMSO (1.4 mL) and purified by preparative HPLC (8 mL/min, 40 % to 80 % MeCN/$H_2O$ (0.1 % TFA) in 55 min) to obtain the intermediate. The corresponding ligand (CA or TF) was then installed via amide coupling by mixing the carboxylic acid intermediate with 2-(2-(6-chlorohexyloxy)ethoxy)ethanamine hydrochloride (CAS: 1035373-85-3, 1.2 eq.) or 4-((4-(aminomethyl)-3-fluorobenzyl)oxy)-6-(trifluoromethyl)pyrimidin-2-amine[2] (1.2 eq.), PyAOP (1.2 eq.) and DIPEA (4.0 eq.) in DMSO (1.0 mL). After stirring at rt for 1 hour, the product was purified by preparative HPLC (8 mL/min, 40 % to 80 % MeCN/$H_2O$ (0.1 % TFA) in 55 min).

**2-((4-(1,4,7,10,13-pentaoxa-16-azacyclooctadecan-16-yl)-3-(2-methoxyethoxy)phenyl)sulfonyl)-N-(4-(((2-amino-6-(trifluoromethyl)pyrimidin-4-yl)oxy)methyl)-2-fluorobenzyl)-3',6'-bis(dimethylamino)-3-oxospiro[isoindoline-1,9'-xanthene]-6-carboxamide (TF-MaPK-558)**

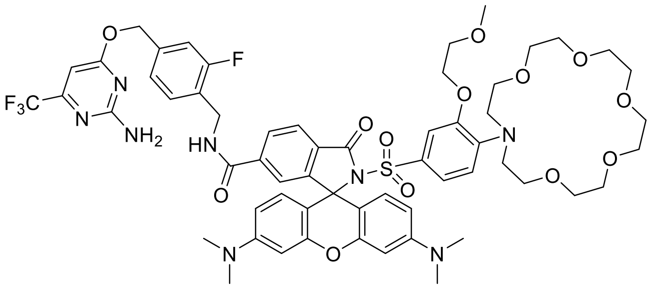

Following GP C, **TF-MaPK-558** (0.8 mg, 0.7 µmol, overall yield 13%) was obtained as a red solid.

**$^1$H NMR** (400 MHz, $CD_3OD$): δ = 8.13 (d, *J* = 8.2 Hz, 1H), 8.02 (d, *J* = 8.1 Hz, 1H), 7.65 (s, 1H), 7.35 (t, *J* = 7.8 Hz, 1H), 7.24 – 7.16 (m, 2H), 6.97 (s, 1H), 6.75 (s, 2H), 6.65 (s, 3H), 6.39 (s, 1H), 5.39 (s, 2H), 4.56 (s, 2H), 4.03 (s, 2H), 3.89 – 3.60 (m, 13H), 3.41 (s, 16H), 3.22 – 3.09 (m, 12H);
**$^{19}$F NMR** (376 MHz, $CD_3OD$): δ = -72.42, -77.35;
**HRMS** (ESI) calcd. for $C_{59}H_{68}N_8O_{13}F_4S$ $[M+2H]^{2+}$ 602.2276; Found 602.2303.

**2'-((4-(1,4,7,10,13-pentaoxa-16-azacyclooctadecan-16-yl)-3-(2-methoxyethoxy)phenyl)sulfonyl)-N-(2-(2-((6-chlorohexyl)oxy)ethoxy)ethyl)-3,6-bis(dimethylamino)-10,10-dimethyl-3'-oxo-10H-spiro[anthracene-9,1'-isoindoline]-6'-carboxamide (CA-MaPK-619)**

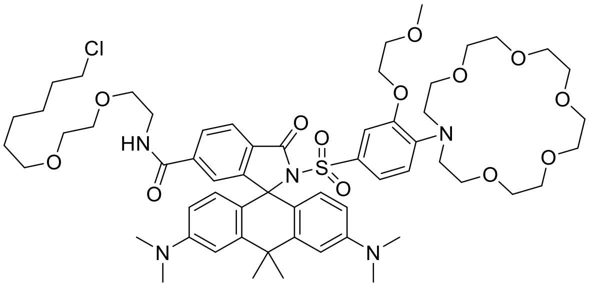

Following GP C, **CA-MaPK-619** (1.0 mg, 0.8 μmol, overall yield 16%) was obtained as a blue solid.

**$^1$H NMR** (400 MHz, $CD_3OD$): δ = 7.98 (d, *J* = 8.1 Hz, 1H), 7.92 (dd, *J* = 8.1, 1.4 Hz, 1H), 7.16 (s, 3H), 6.60 (d, *J* = 9.5 Hz, 3H), 6.45 (d, *J* = 8.8 Hz, 2H), 3.90 (d, *J* = 4.7 Hz, 2H), 3.81 – 3.59 (m, 15H), 3.58 – 3.34 (m, 26H), 3.05 (s, 12H), 2.01 (s, 3H), 1.93 (s, 3H), 1.68 (p, *J* = 6.7 Hz, 2H), 1.47 (p, *J* = 6.8 Hz, 2H), 1.38 (p, *J* = 7.0 Hz, 2H), 1.29 (q, *J* = 9.0, 8.1 Hz, 2H);
**HRMS** (ESI) calcd. for $C_{59}H_{84}N_5O_{13}ClS$ $[M+2H]^{2+}$ 568.7732; Found 568.7735.

**2'-((4-(1,4,7,10,13-pentaoxa-16-azacyclooctadecan-16-yl)-3-(2-methoxyethoxy)phenyl)sulfonyl)-N-(2-(2-((6-chlorohexyl)oxy)ethoxy)ethyl)-3,7-bis(dimethylamino)-5,5-dimethyl-3'-oxo-5H-spiro[dibenzo[b,e]siline-10,1'-isoindoline]-6'-carboxamide (CA-MaPK-656)**

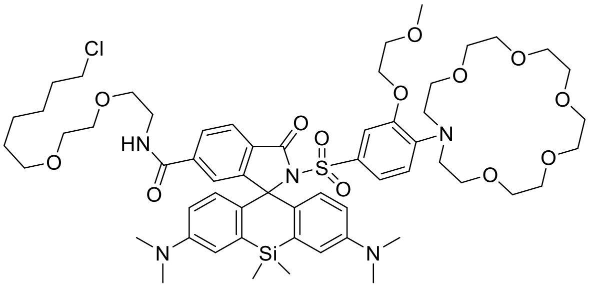

Following GP C, **CA-MaPK-656** (0.8 mg, 0.7 μmol, overall yield 8%) was obtained as a green-blue solid.

**$^1$H NMR** (400 MHz, $CD_3OD$): δ = 8.02 – 7.95 (m, 1H), 7.91 (dd, *J* = 8.0, 1.4 Hz, 1H), 7.15 (s, 2H), 6.65 (d, *J* = 9.1 Hz, 2H), 6.59 – 6.43 (m, 3H), 3.90 (t, *J* = 4.4 Hz, 2H), 3.83 – 3.61 (m, 12H), 3.60 – 3.45 (m, 11H), 3.45 – 3.32 (m, 17H), 3.03 (s, 12H), 1.68 (dt, *J* = 14.3, 6.7 Hz, 2H), 1.46 (dq, *J* = 12.3, 6.2, 5.7 Hz, 2H), 1.41 – 1.33 (m, 2H), 1.28 (q, *J* = 8.9, 8.1 Hz, 2H), 0.66 (s, 6H);
**HRMS** (ESI) calcd. for $C_{58}H_{84}N_5O_{13}ClSSi$ $[M+2H]^{2+}$ 576.7617; Found 576.7615.

**2-((4-(1,4,7,10-Tetraoxa-13-azacyclopentadecan-13-yl)-3-methoxyphenyl)sulfonyl)-N-(4-(((2-amino-6-(trifluoromethyl)pyrimidin-4-yl)oxy)methyl)-2-fluorobenzyl)-3',6'-bis(dimethylamino)-3-oxospiro[isoindoline-1,9'-xanthene]-6-carboxamide (TF-MaPNa-558)**

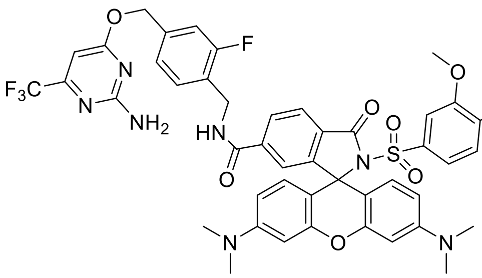

Following GP C, **TF-MaPNa-558** (1.8 mg, 1.5 µmol, overall yield 15%) was obtained as a red solid.

**$^{1}$H NMR** (400 MHz, $CD_3OD$): δ = 8.16 (dd, *J* = 8.1, 1.7 Hz, 1H), 7.98 (d, *J* = 8.1 Hz, 1H), 7.73 (s, 1H), 7.37 (t, *J* = 7.8 Hz, 1H), 7.24 – 7.15 (m, 3H), 7.12 (s, 1H), 6.89 (d, *J* = 7.7 Hz, 2H), 6.80 (d, *J* = 9.1 Hz, 3H), 6.37 (s, 1H), 5.38 (s, 2H), 4.58 (s, 2H), 3.67 (t, *J* = 7.4 Hz, 23H), 3.24 (s, 12H);
**$^{19}$F NMR** (376 MHz, $CD_3OD$): δ = -72.42, -77.41;
**HRMS** (ESI) calcd. for $C_{55}H_{60}N_8O_{11}F_4S$ $[M+2H]^{2+}$ 558.2014; Found 558.2018.

**2'-((4-(1,4,7,10-Tetraoxa-13-azacyclopentadecan-13-yl)-3-methoxyphenyl)sulfonyl)-N-(2-(2-((6-chlorohexyl)oxy)ethoxy)ethyl)-3,6-bis(dimethylamino)-10,10-dimethyl-3'-oxo-10H-spiro[anthracene-9,1'-isoindoline]-6'-carboxamide (CA-MaPNa-619)**

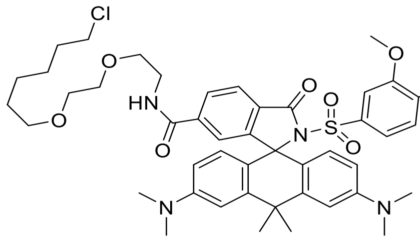

Following GP C, **CA-MaPNa-619** (1.0 mg, 0.9 µmol, overall yield 9%) was obtained as a blue solid.

**$^{1}$H NMR** (400 MHz, $CD_3OD$): 8.01 (d, *J* = 8.0 Hz, 1H), 7.94 (dd, *J* = 8.0, 1.4 Hz, 1H), 7.39 – 7.10 (m, 4H), 6.85 (s, 1H), 6.69 (d, *J* = 8.9 Hz, 2H), 6.51 (d, *J* = 8.9 Hz, 2H), 3.75 – 3.45 (m, 35H), 3.11 (s, 12H), 2.02 (s, 3H), 1.96 (s, 3H), 1.75 – 1.65 (m, 2H), 1.49 (p, *J* = 6.8 Hz, 2H), 1.40 (p, *J* = 7.0 Hz, 2H), 1.31 (td, *J* = 6.8, 6.4, 3.0 Hz, 2H);
**HRMS** (ESI) calcd. for $C_{55}H_{76}N_5O_{11}ClS$ $[M+2H]^{2+}$ 524.7470; Found 524.7474.

**2'-((4-(1,4,7,10-Tetraoxa-13-azacyclopentadecan-13-yl)-3-methoxyphenyl)sulfonyl)-N-(2-(2-((6-chlorohexyl)oxy)ethoxy)ethyl)-3,7-bis(dimethylamino)-5,5-dimethyl-3'-oxo-5H-spiro[dibenzo[b,e]siline-10,1'-isoindoline]-6'-carboxamide (CA-MaPNa-656)**

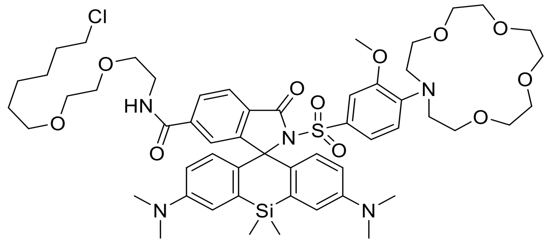

Following GP C, **CA-MaPNa-656** (0.7 mg, 0.6 μmol, overall yield 6%) was obtained as a green-blue solid.

**$^{1}$H NMR** (400 MHz, $CD_3OD$): δ = 7.99 (d, *J* = 8.1 Hz, 1H), 7.91 (d, *J* = 8.1 Hz, 1H), 7.24 – 7.09 (m, 4H), 6.79 (s, 1H), 6.66 (d, *J* = 9.0 Hz, 2H), 6.51 (d, *J* = 9.0 Hz, 2H), 3.72 – 3.43 (m, 35H), 3.05 (s, 12H), 1.70 (p, *J* = 6.9 Hz, 2H), 1.48 (p, *J* = 6.7 Hz, 2H), 1.38 (dd, *J* = 15.6, 7.9 Hz, 2H), 1.33 – 1.28 (m, 2H), 0.67 (s, 6H);

**HRMS** (ESI) calcd. for $C_{54}H_{76}N_5O_{11}ClSSi$ $[M+2H]^{2+}$ 532.7354; Found 532.7364.

#### 6. Supplementary References

(1)Frei, M. S.; Tarnawski, M.; Roberti, M. J.; Koch, B.; Hiblot, J.; Johnsson, K. Engineered HaloTag variants for fluorescence lifetime multiplexing. *Nat. Methods* **2022**, *19* (1), 65–70.

(2)Kühn, S.; Nasufovic, V.; Wilhelm, J.; Kompa, J.; De Lange, E. M. F.; Lin, Y.-H.; Egoldt, C.; Fischer, J.; Lennoi, A.; Tarnawski, M.; Reinstein, J.; Vlijm, R.; Hiblot, J.; Johnsson, K. SNAP-Tag2 for faster and brighter protein labeling. *Nat. Chem. Biol.* **2025**, 21, 1754–1761.

(3)Mertes, N.; Busch, M.; Huppertz, M.-C.; Hacker, C. N.; Wilhelm, J.; Gürth, C.-M.; Kühn, S.; Hiblot, J.; Koch, B.; Johnsson, K. Fluorescent and bioluminescent calcium indicators with tuneable colors and affinities. *J. Am. Chem. Soc.* **2022**, *144* (15), 6928–6935.

(4)Smith, R. M.; Martell, A. E. *Critical Stability Constants*; Springer US: Boston, MA, 1989.

(5)Zeng, L.; Miller, E. W.; Pralle, A.; Isacoff, E. Y.; Chang, C. J. A selective turn-on fluorescent sensor for imaging copper in living cells. *J. Am. Chem. Soc.* **2006**, *128* (1), 10–11.

(6)Sun, D.; Ng, S. W.; Zheng, Y.; Xie, S.; Schwan, N.; Breuer, P.; Hoffmann, D. C.; Michel, J.; Azorin, D. D.; Boonekamp, K. E.; Winkler, F.; Wick, W.; Boutros, M.; Li, Y.; Johnsson, K. Molecular Recording of Cellular Protein Kinase Activity with Chemical Labeling. *Nat. Chem. Biol.* **2025**. 21, 1818–1827.

(7)Schindelin, J.; Arganda-Carreras, I.; Frise, E.; Kaynig, V.; Longair, M.; Pietzsch, T.; Preibisch, S.; Rueden, C.; Saalfeld, S.; Schmid, B.; Tinevez, J.-Y.; White, D. J.; Hartenstein, V.; Eliceiri, K.; Tomancak, P.; Cardona, A. Fiji: An open-source platform for biological-image analysis. *Nat. Methods* **2012**, *9* (7), 676–682.

(8)Pachitariu, M.; Stringer, C. Cellpose 2.0: How to train your own model. *Nat. Methods* **2022**, *19* (12), 1634–1641.

(9)Wang, M.-M.; Johnsson, K. Metal-free introduction of primary sulfonamide into electron-rich aromatics. *Chem. Sci.* **2024**, *15* (31), 12310–12315.

### 7. NMR-Spectra of new compounds

**4-(1,4,7,10-Tetraoxa-13-azacyclopentadecan-13-yl)-3-methoxybenzenesulfonamide (2)**
**$^1$H-NMR (400 MHz, $CDCl_3$)**

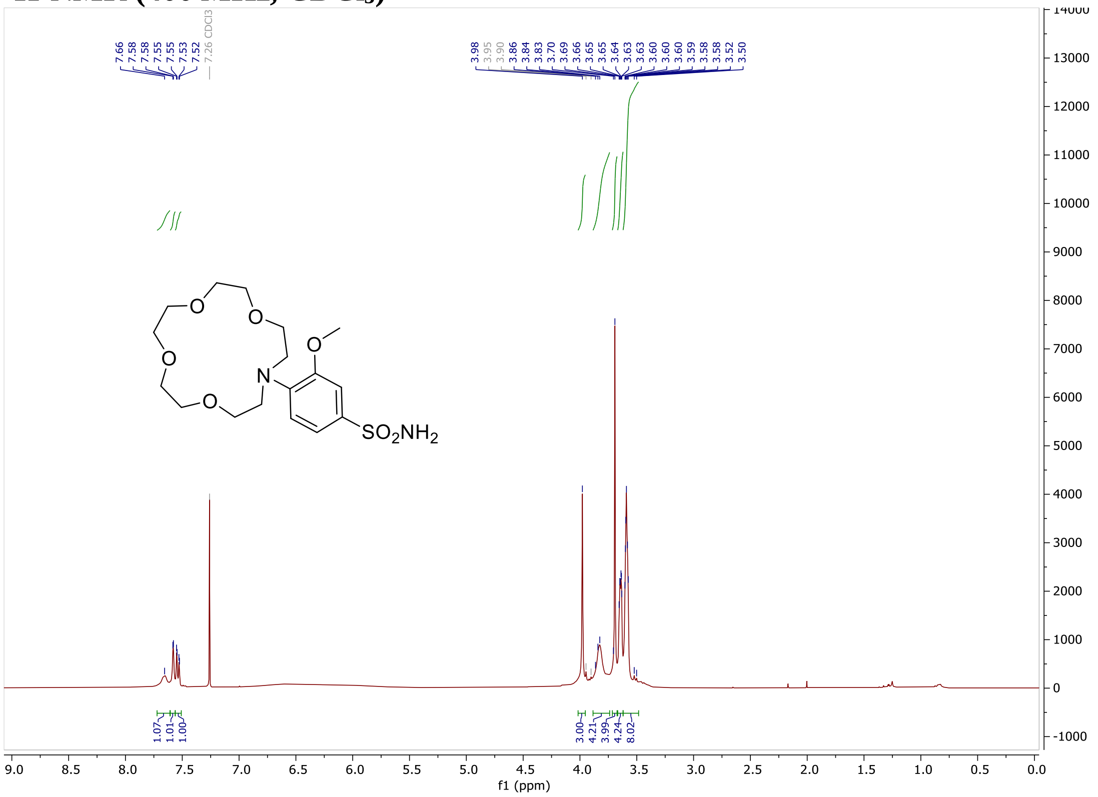

**$^{13}$C-NMR (101 MHz, $CDCl_3$)**

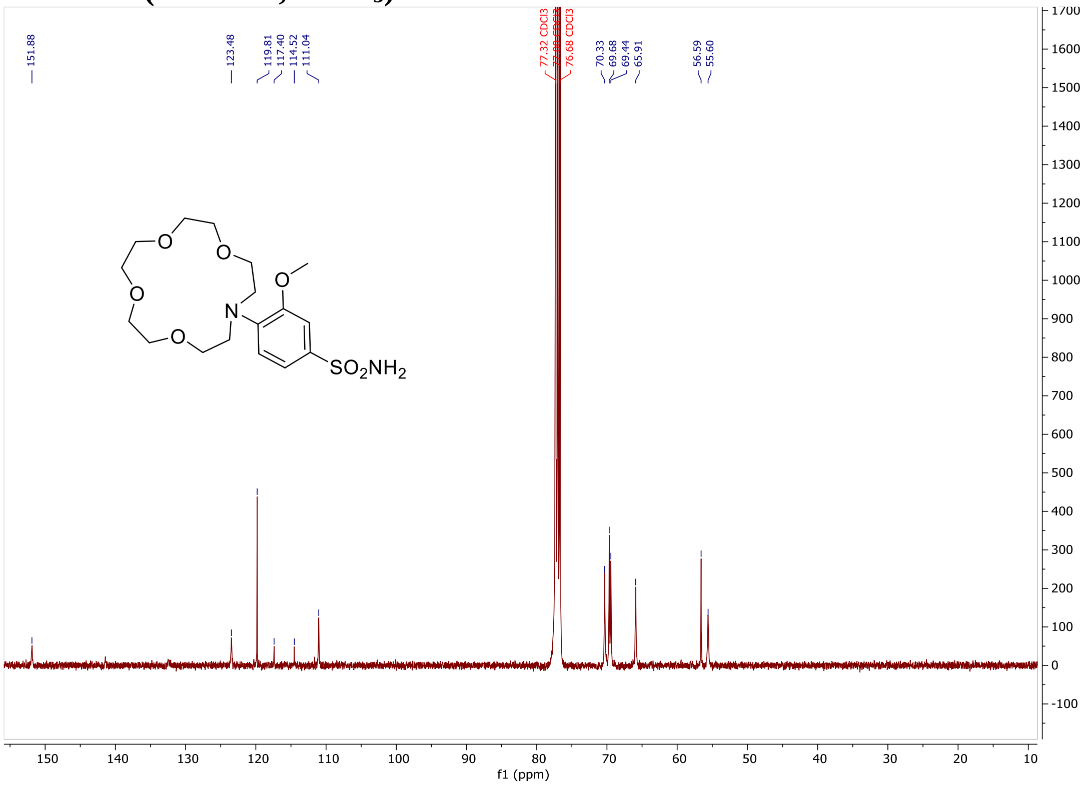

**((4-(N-(tert-Butoxycarbonyl)sulfamoyl)phenyl)azanediyl)bis(ethane-2,1-diyl) bis(4-methylbenzenesulfonate) (9)**

**$^{1}$H-NMR (400 MHz, $CDCl_3$)**

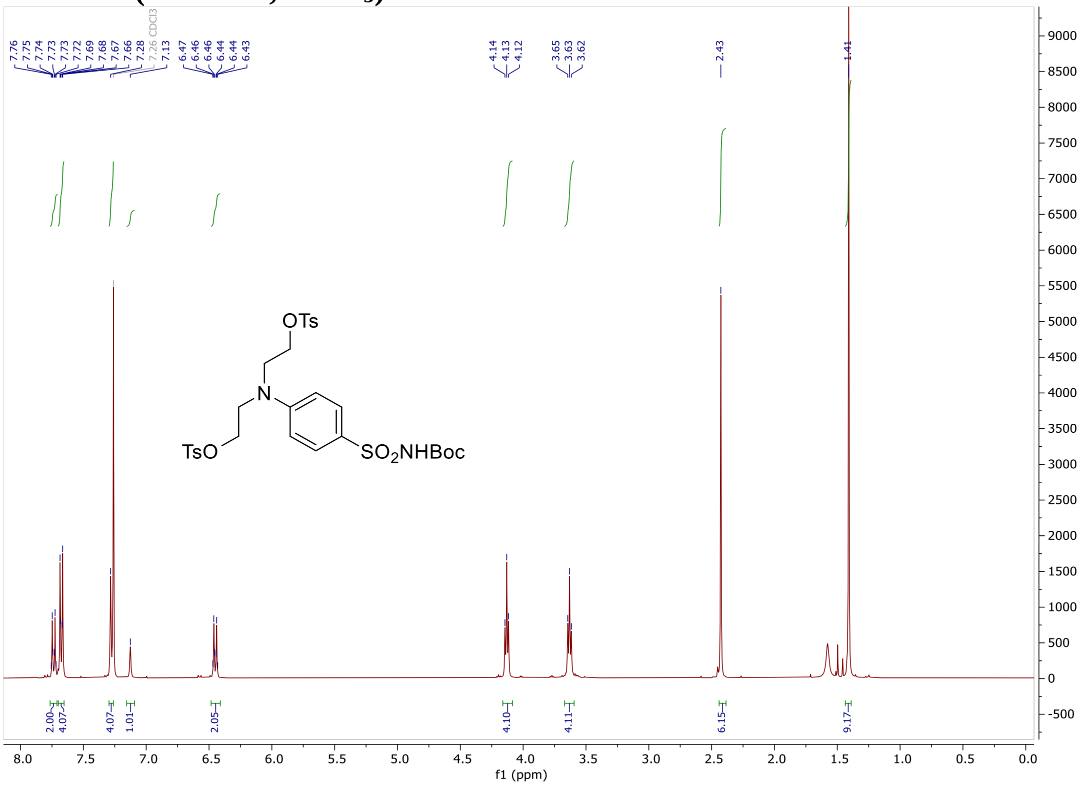

#### $^{13}$C-NMR (101 MHz, $CDCl_3$)

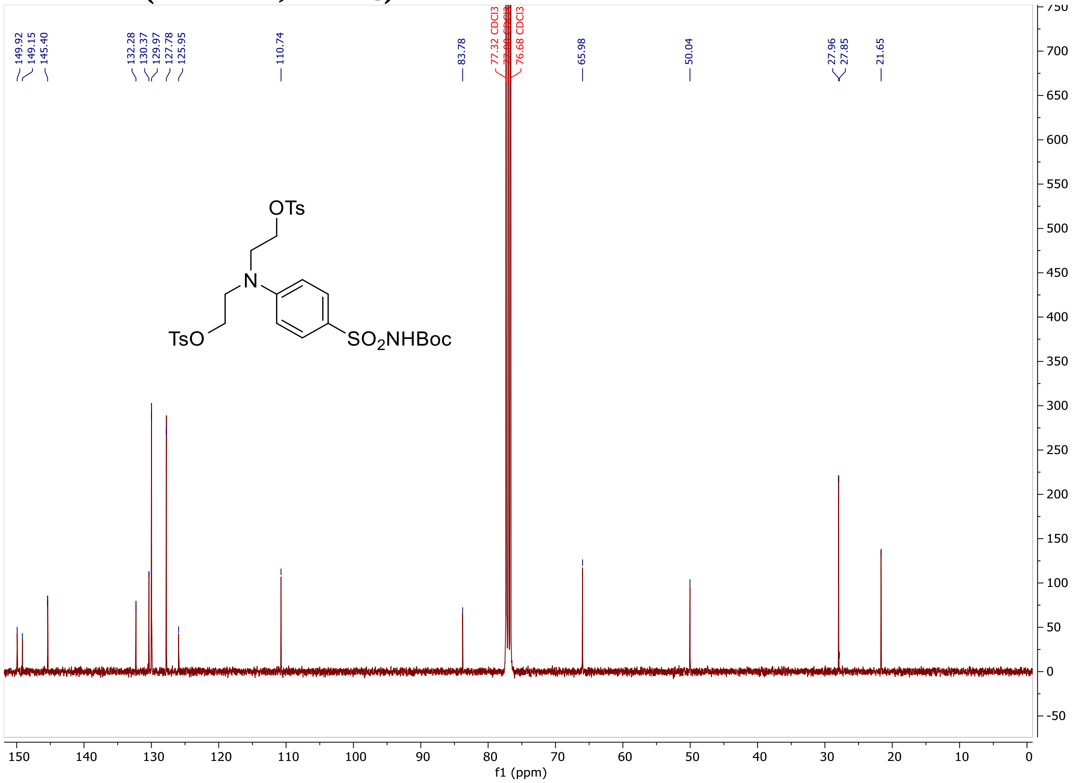

#### tert-Butyl ((4-(bis(2-((2-(ethylthio)ethyl)thio)ethyl)amino)phenyl)sulfonyl)carbamate (3)

#### $^{1}$H-NMR (400 MHz, $CDCl_3$)

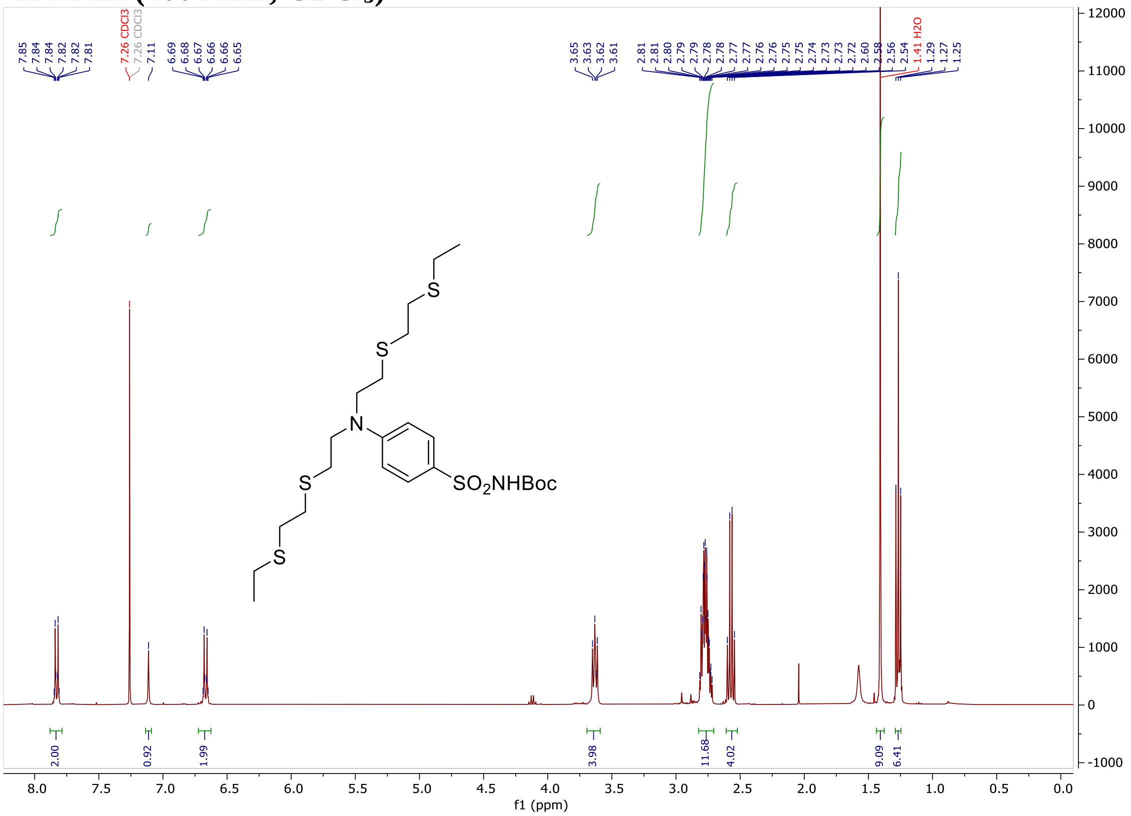

#### $^{13}$C-NMR (101 MHz, $CDCl_3$)

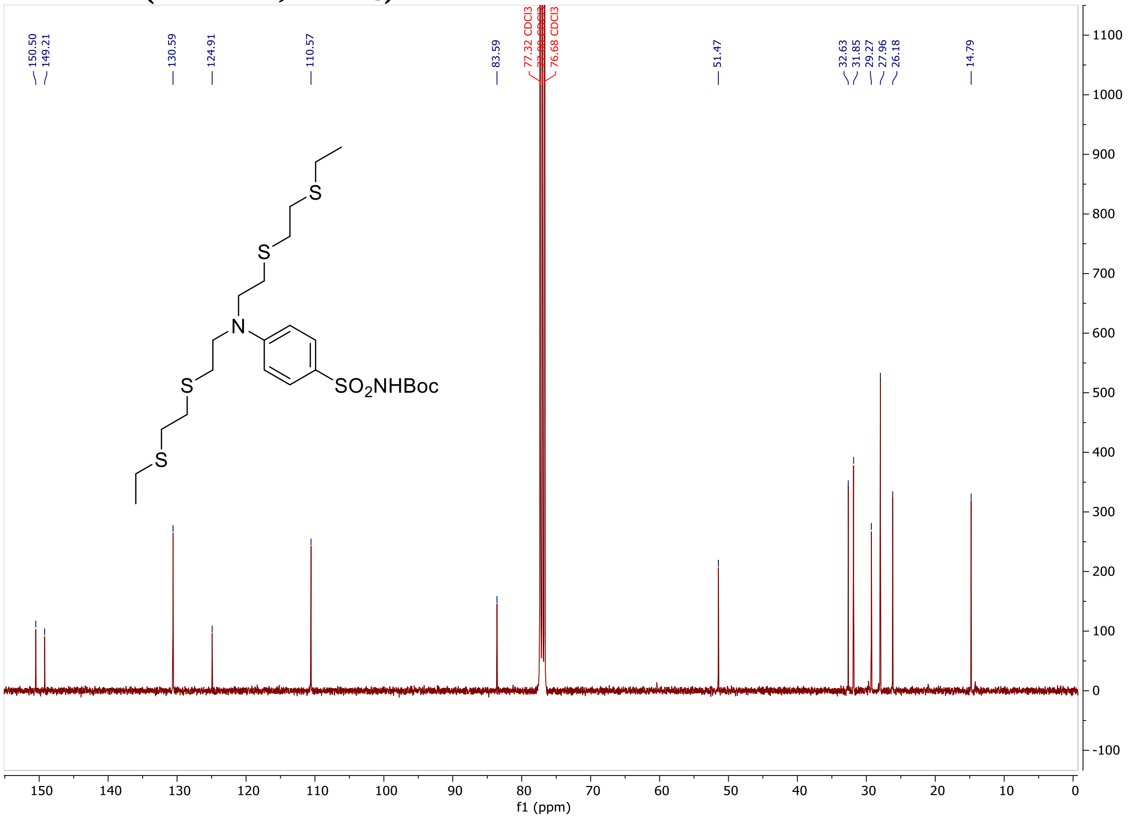

#### Dimethyl 2,2'-((((4-(N-(tert-butoxycarbonyl)sulfamoyl)phenyl)azanediyl)bis(ethane-2,1-diyl))bis(sulfanediyl))diacetate (4)

#### $^{1}$H-NMR (400 MHz, $CDCl_3$)

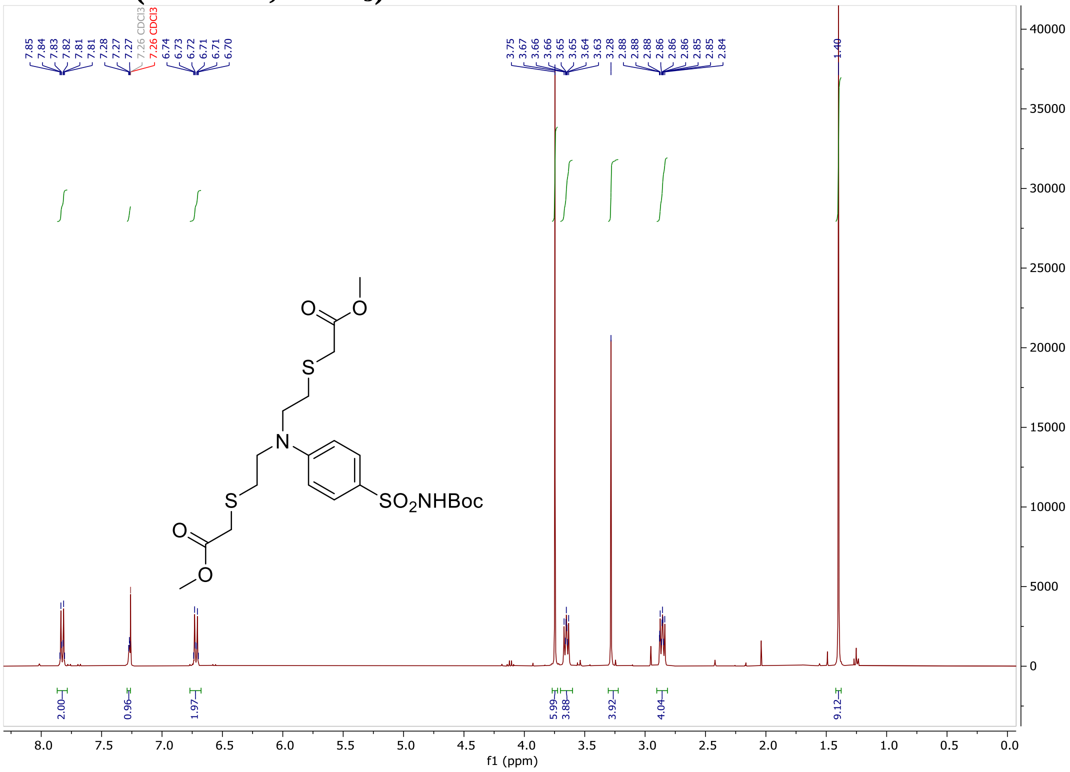

#### [13]C-NMR (101 MHz, $CDCl_3$)

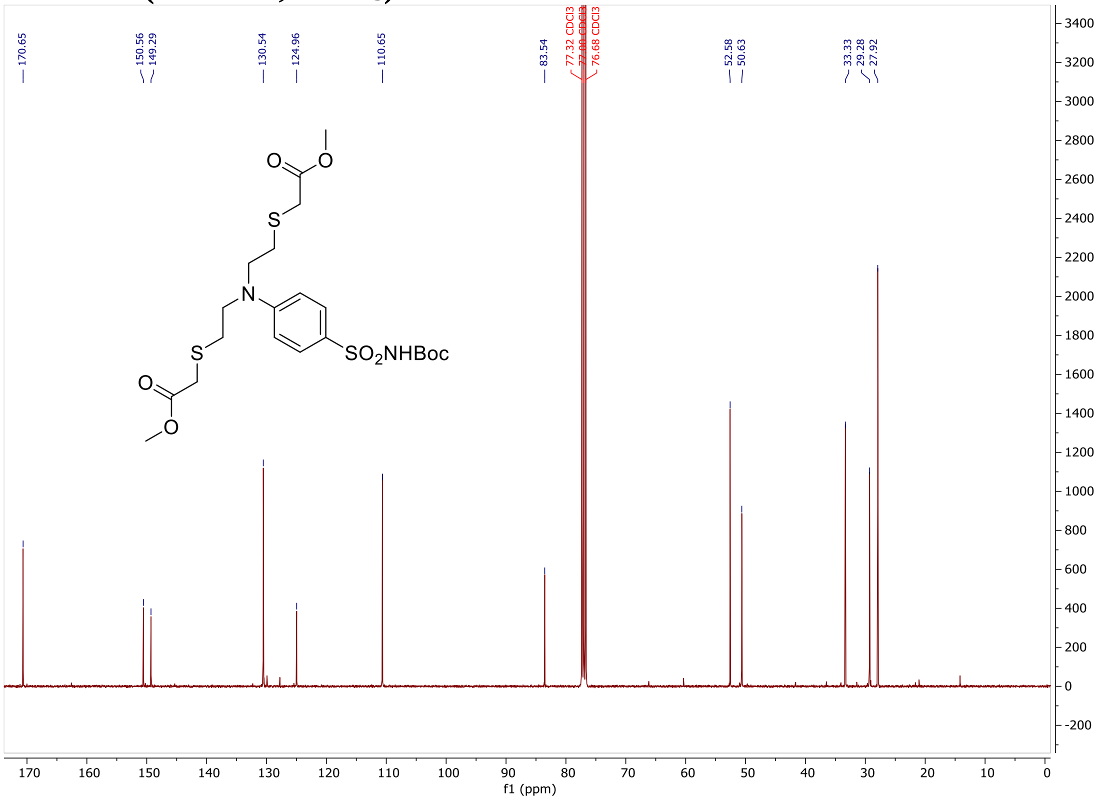

#### Diethyl 2,2'-((2-(2-ethoxy-2-oxoethoxy)-4-sulfamoylphenyl)azanediyl)diacetate (6)

#### [1]H-NMR (400 MHz, $CDCl_3$)

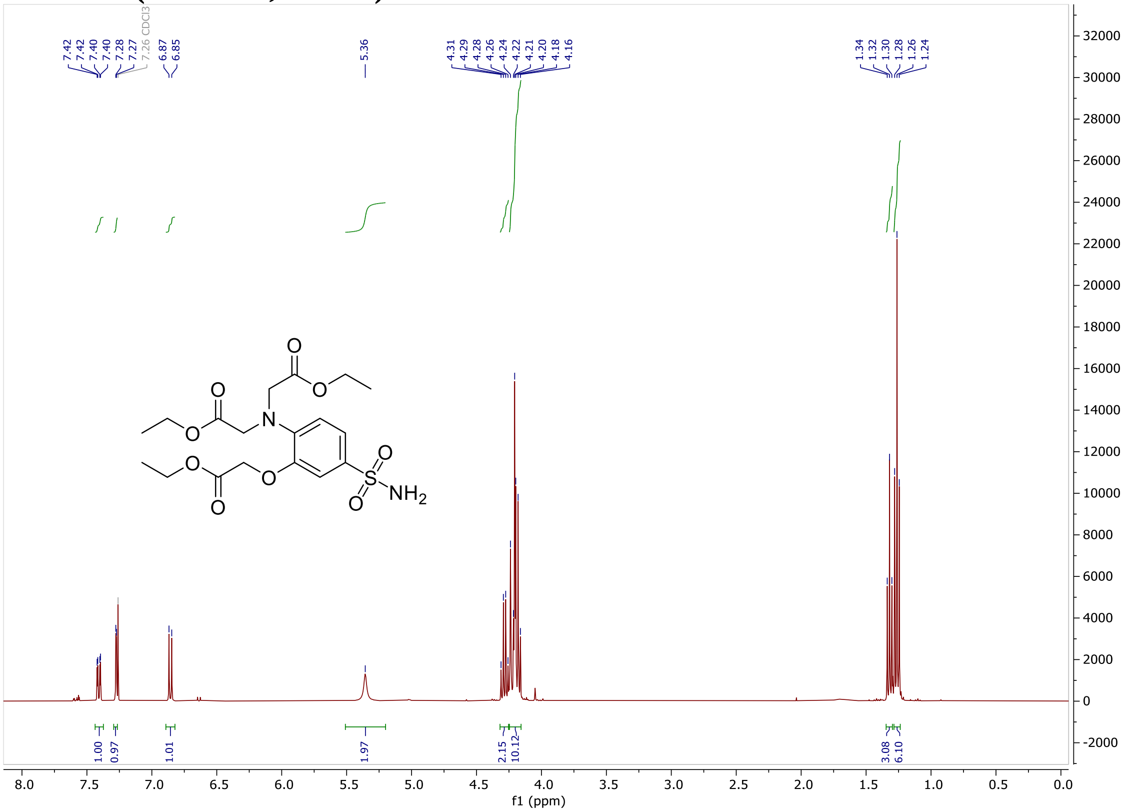

#### $^{13}$C-NMR (101 MHz, $CDCl_3$)

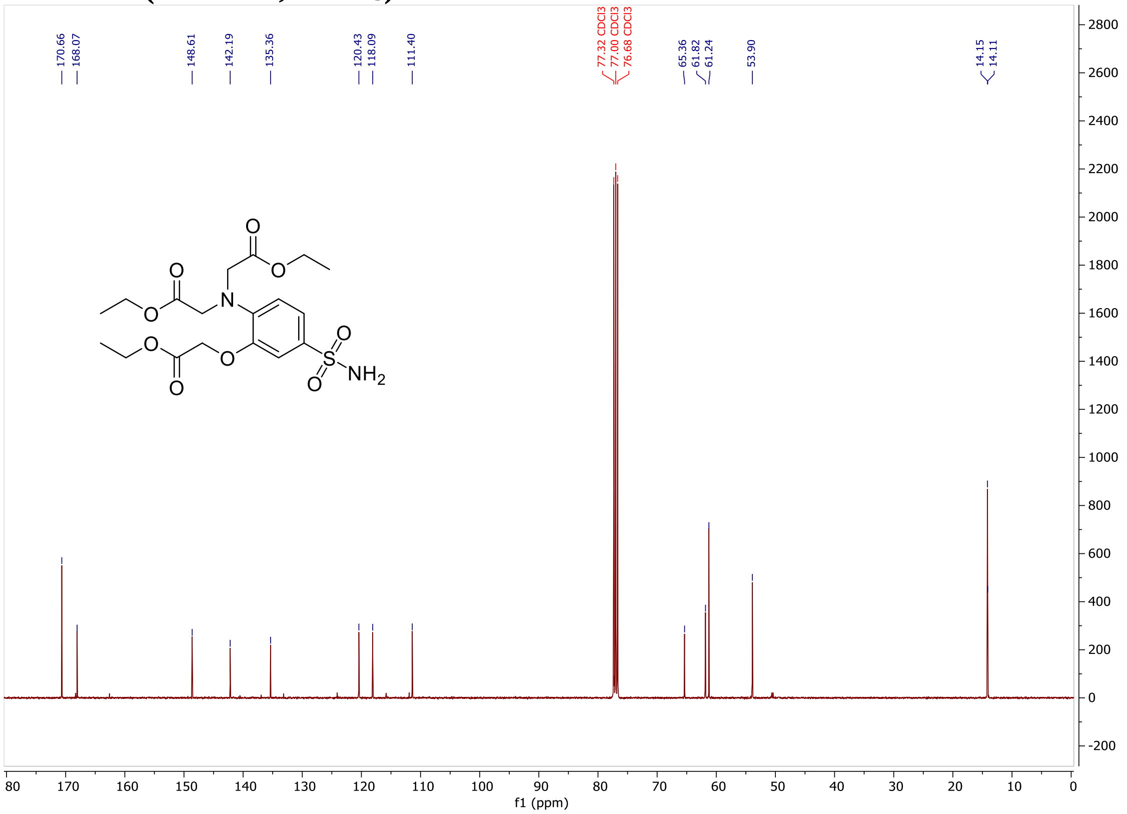

#### Diethyl 2,2'-((2-methoxy-4-sulfamoylphenyl)azanediyl)diacetate (7)

#### $^{1}$H-NMR (400 MHz, $CDCl_3$)

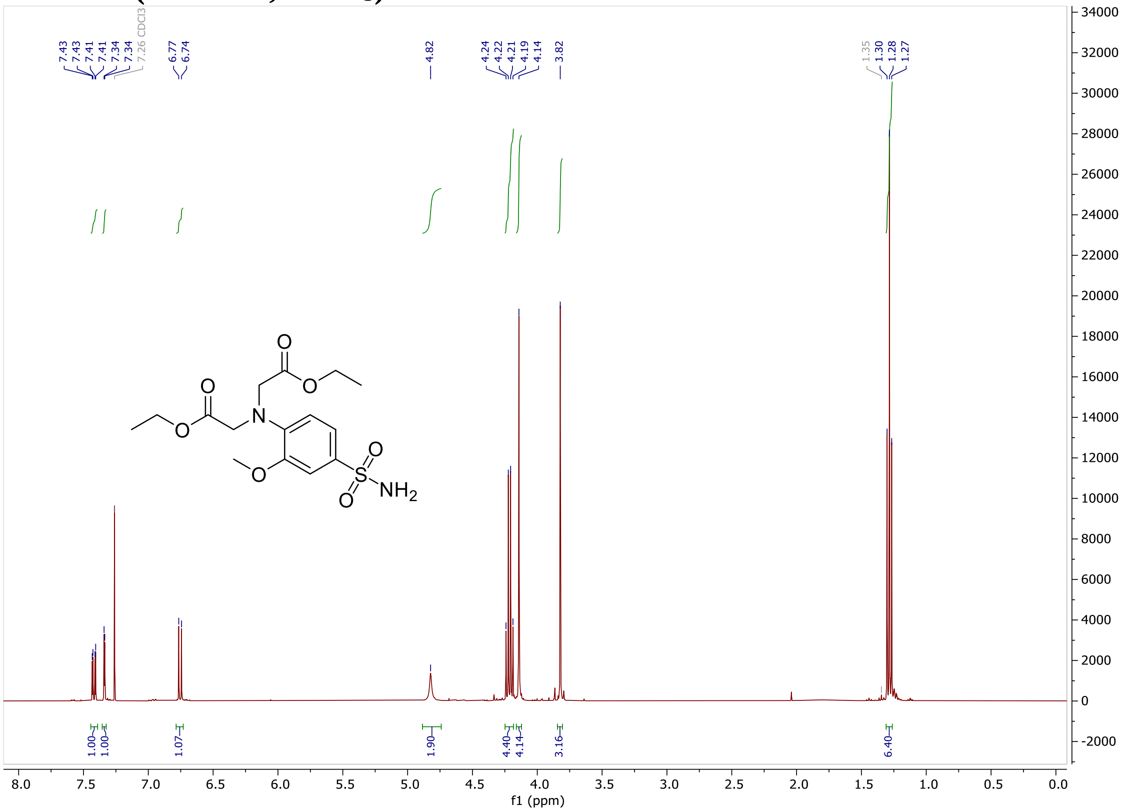

#### $^{13}$C-NMR (101 MHz, $CDCl_3$)

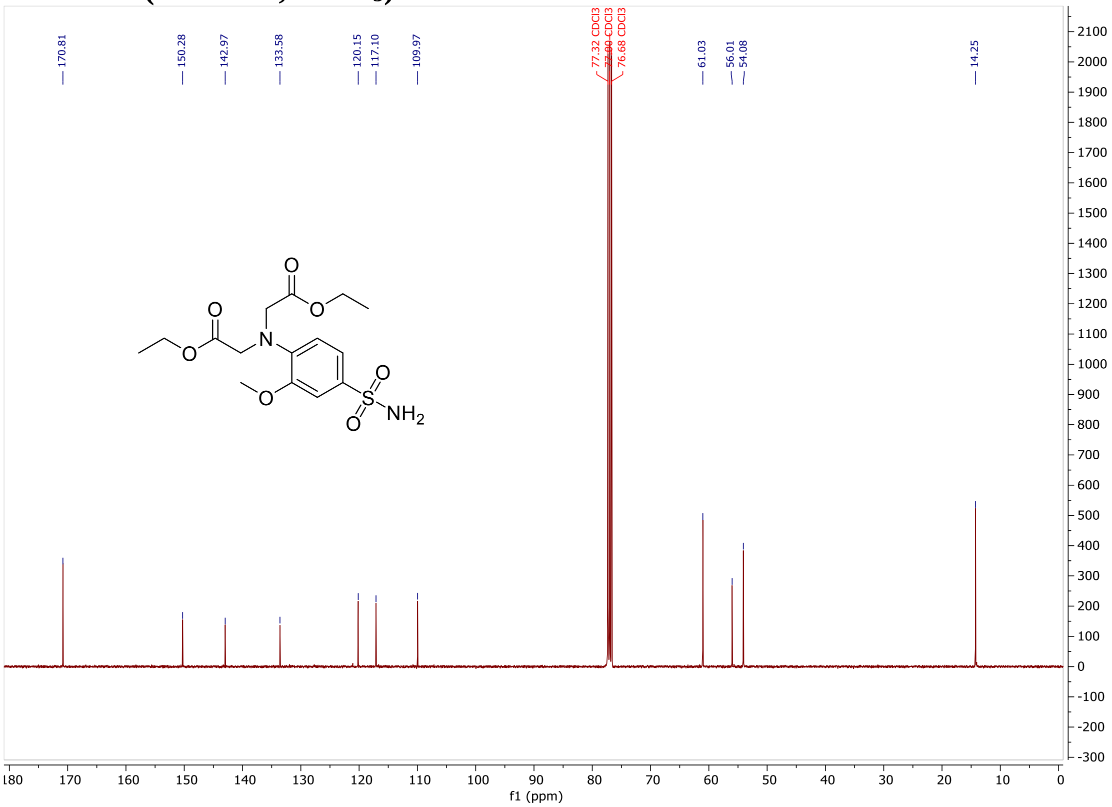

#### tert-butyl (N-(1,10-phenanthrolin-5-yl)sulfamoyl)carbamate (8)

#### $^{1}$H-NMR (400 MHz, $CDCl_3$)

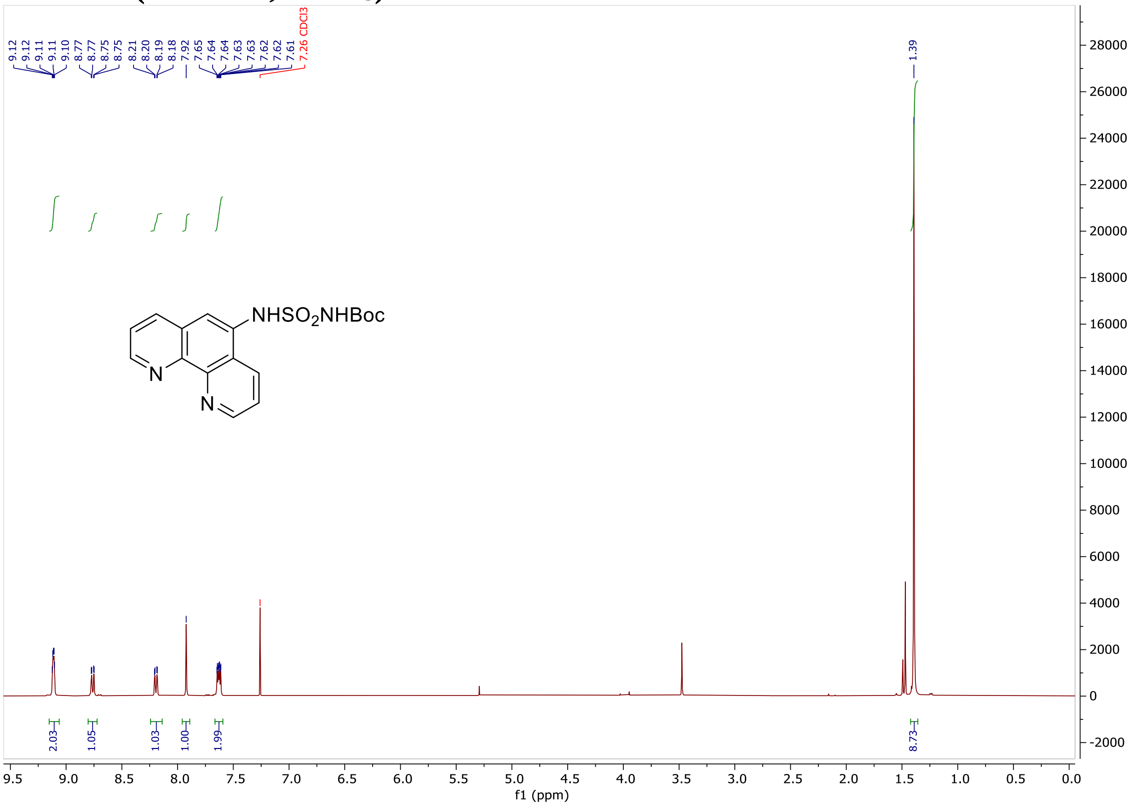

#### $^{13}$C-NMR (101 MHz, $CDCl_3$)

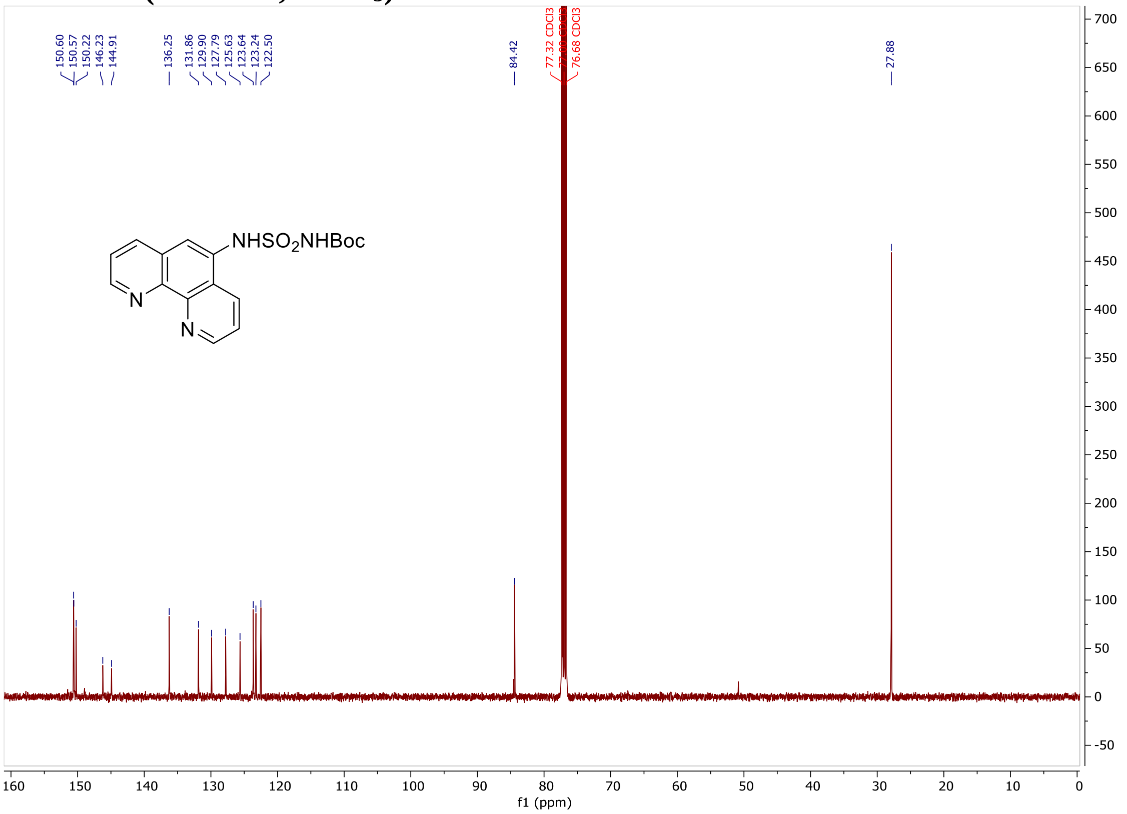

**2-((4-(1,4,7,10,13-Pentaoxa-16-azacyclooctadecan-16-yl)-3-(2-methoxyethoxy)phenyl)sulfonyl)-N-(2-(2-((6-chlorohexyl)oxy)ethoxy)ethyl)-3',6'-bis(dimethylamino)-3-oxospiro[isoindoline-1,9'-xanthene]-6-carboxamide (CA-MaPK-558)**
**$^1$H-NMR (400 MHz, $CD_3OD$)**

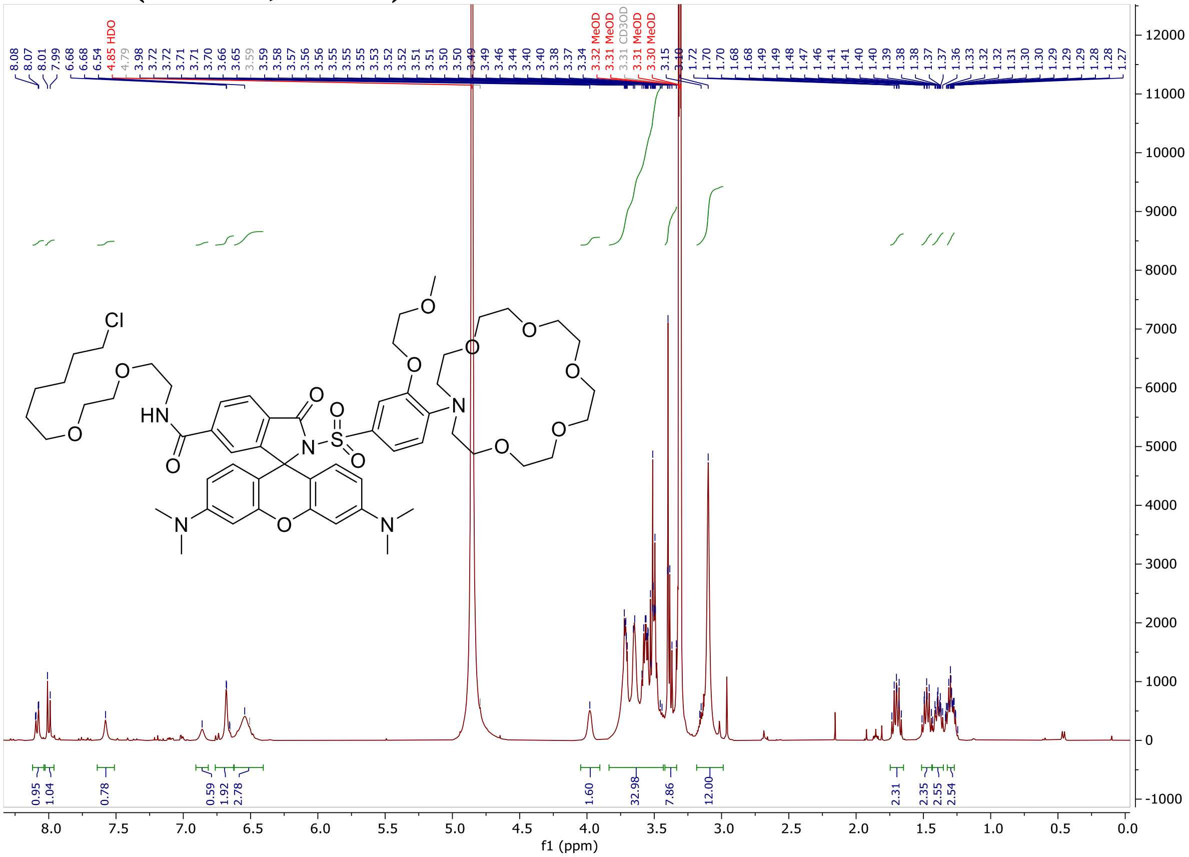

**2-((4-(1,4,7,10-tetraoxa-13-azacyclopentadecan-13-yl)-3-methoxyphenyl)sulfonyl)-N-(2-(2-((6-chlorohexyl)oxy)ethoxy)ethyl)-3',6'-bis(dimethylamino)-3-oxospiro[isoindoline-1,9'-xanthene]-6-carboxamide (CA-MaPNa-558)**

**$^{1}$H-NMR (400 MHz, $CD_3OD$)**

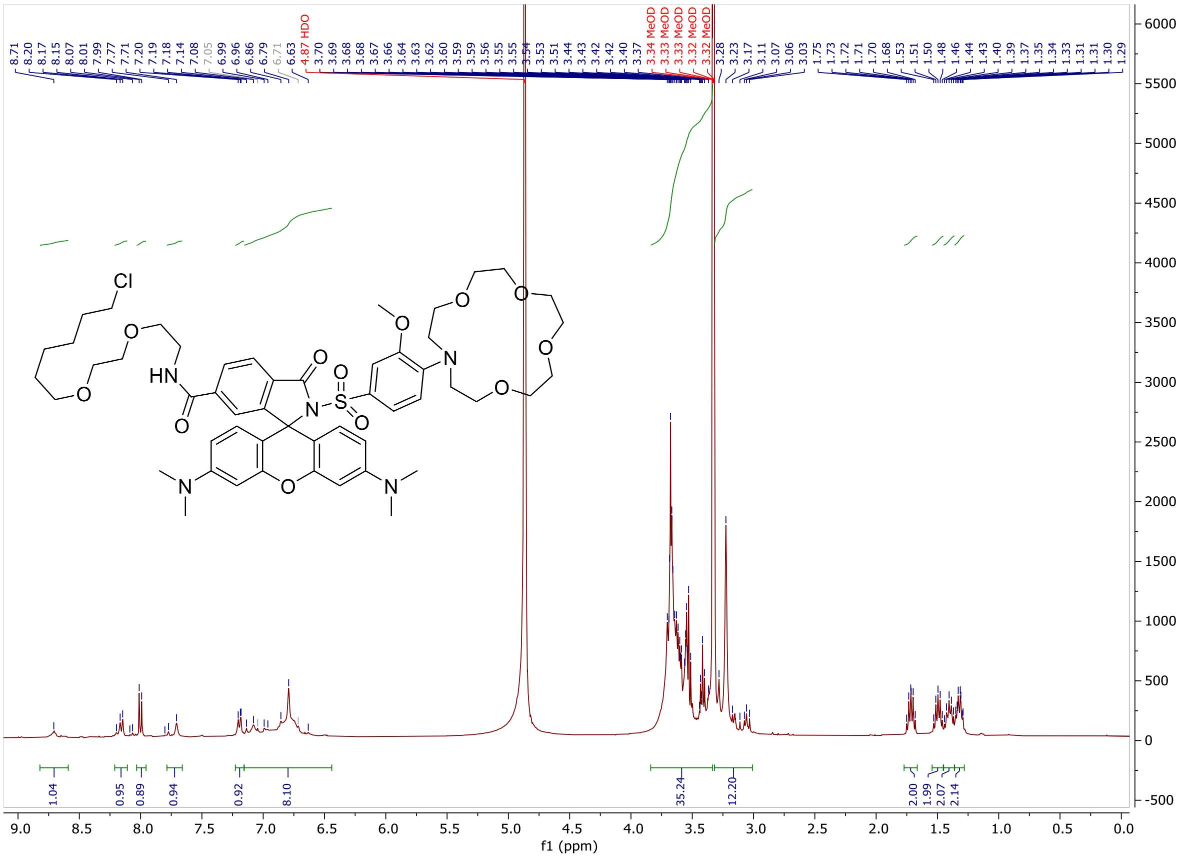

**Bis(acetoxymethyl) 2,2'-((2-(2-(2-(bis(2-(acetoxymethoxy)-2-oxoethyl)amino)-5-((6-((2-(2-((6-chlorohexyl)oxy)ethoxy)ethyl)carbamoyl)-3',6'-bis(dimethylamino)-3-oxospiro[isoindoline-1,9'-xanthen]-2-yl)sulfonyl)phenoxy)ethoxy)phenyl)azanediyl)diacetate (CA-MaPCa2-558)**

**$^1$H-NMR (400 MHz, DMSO-d6)**

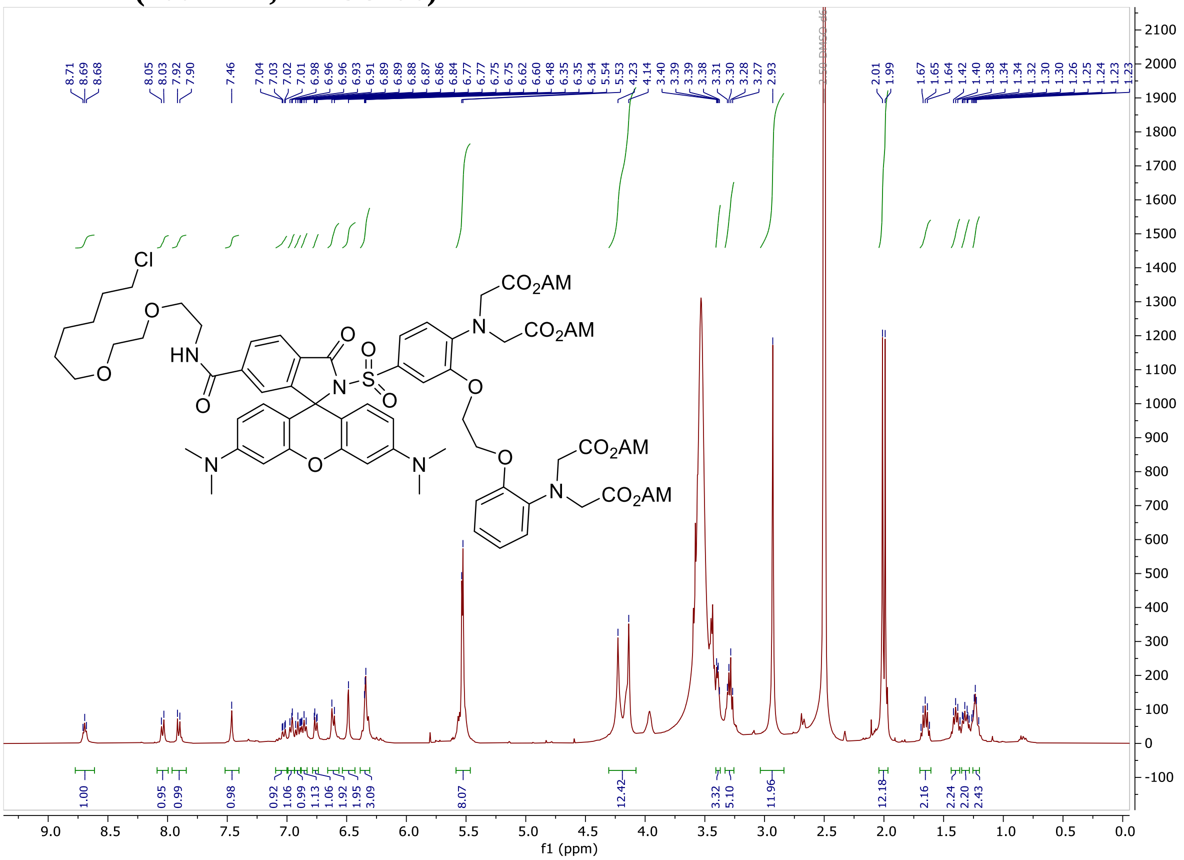

**Diethyl 2,2'-((4-((6-((2-(2-((6-chlorohexyl)oxy)ethoxy)ethyl)carbamoyl)-3',6'-bis(dimethylamino)-3-oxospiro[isoindoline-1,9'-xanthen]-2-yl)sulfonyl)-2-methoxyphenyl)azanediyl)diacetate (CA-MaPZn-558)**

**$^1$H-NMR (400 MHz, DMSO-d6)**

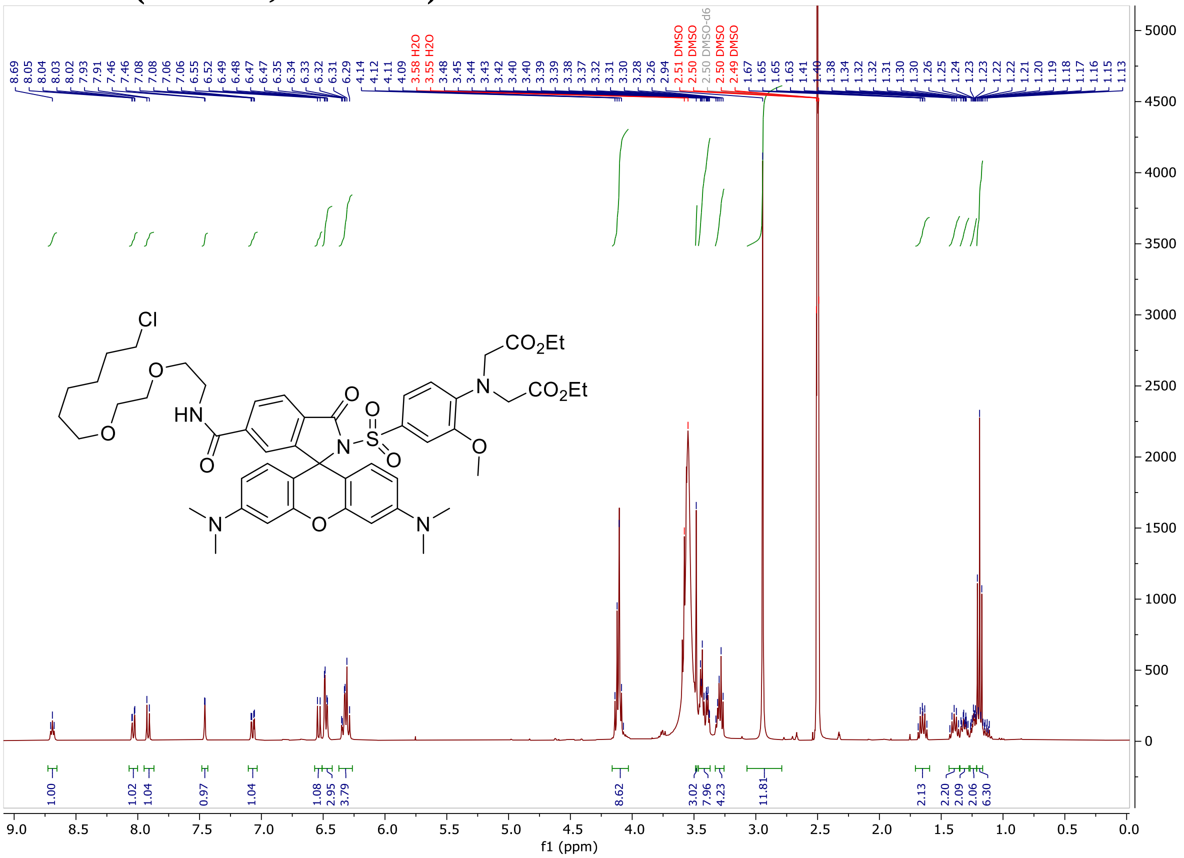

**$^{13}$C-NMR (101 MHz, DMSO-d6)**

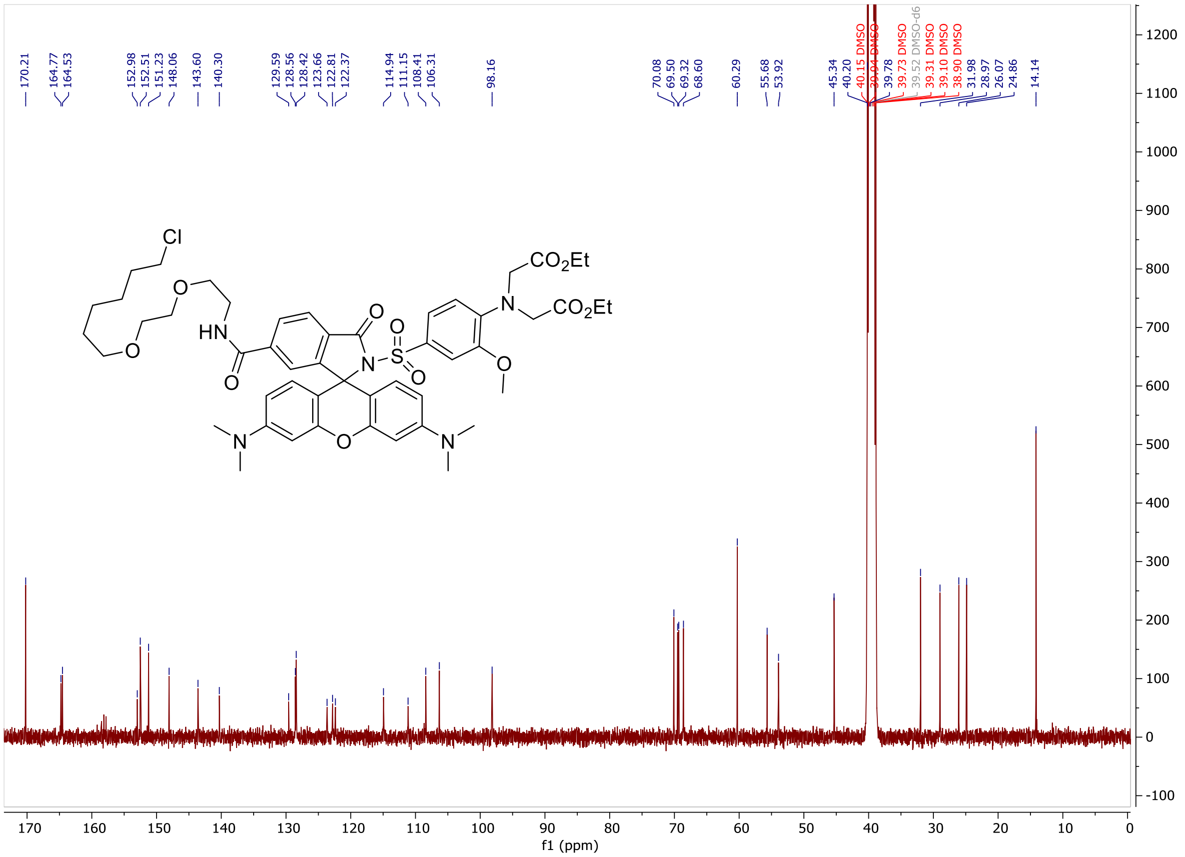
170.21
164.77
164.53
152.98
152.51
151.23
148.06
143.60
140.30
129.59
128.56
128.42
123.66
122.81
122.37
114.94
111.15
108.41
106.31
98.16
70.08
69.50
69.32
68.60
60.29
55.68
53.92
45.34
40.20
40.15 DMSO
39.94 DMSO
39.78
39.73 DMSO
39.52 DMSO-d6
39.31 DMSO
39.10 DMSO
38.90 DMSO
31.98
28.97
26.07
24.86
14.14
Cl
O
HN
O
O
N
S
O
O
N
CO2Et
CO2Et
O
N
O
N
f1 (ppm)

**Diethyl 2,2'-((4-((6-((2-(2-((6-chlorohexyl)oxy)ethoxy)ethyl)carbamoyl)-3',6'-bis(dimethylamino)-3-oxospiro[isoindoline-1,9'-xanthen]-2-yl)sulfonyl)-2-(2-ethoxy-2-oxoethoxy)phenyl)azanediyl)diacetate (CA-MaPMg-558)**

**$^1$H-NMR (400 MHz, DMSO-d6)**

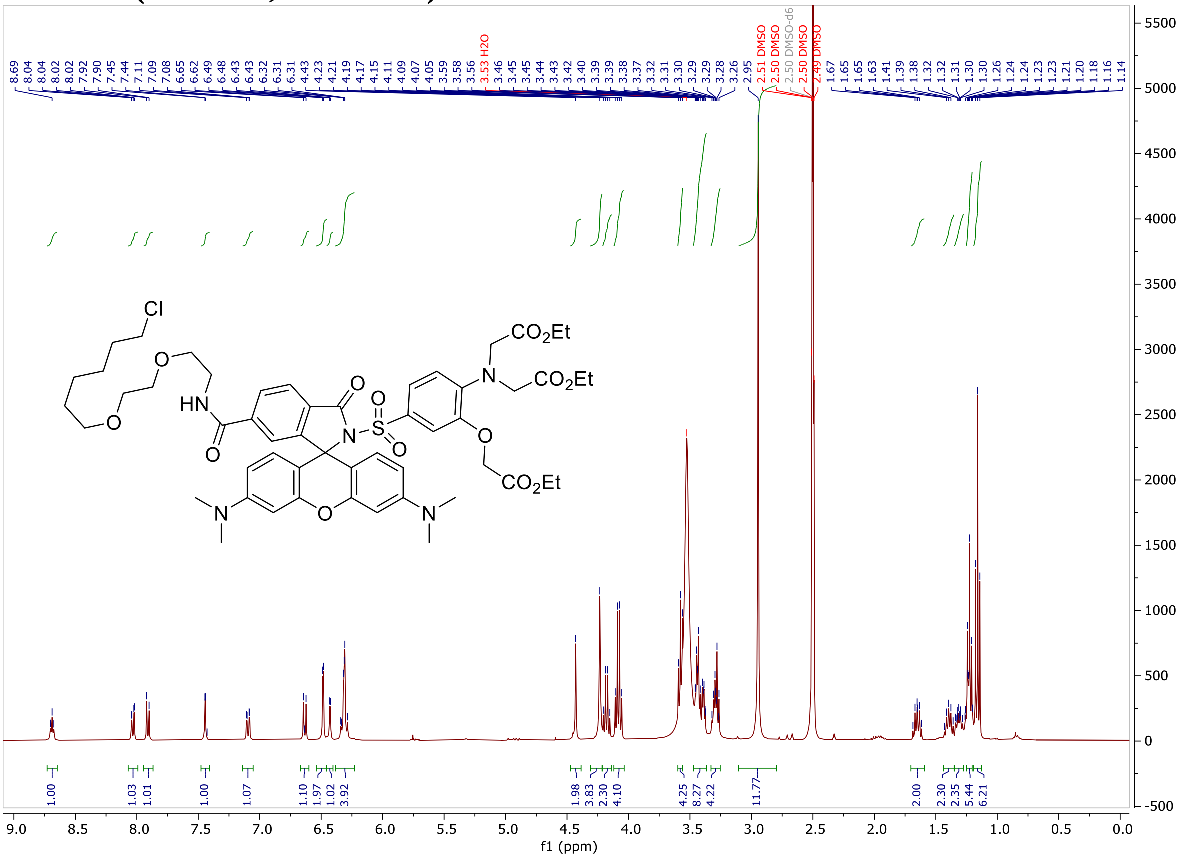

### $^{13}$C-NMR (101 MHz, DMSO-d6)

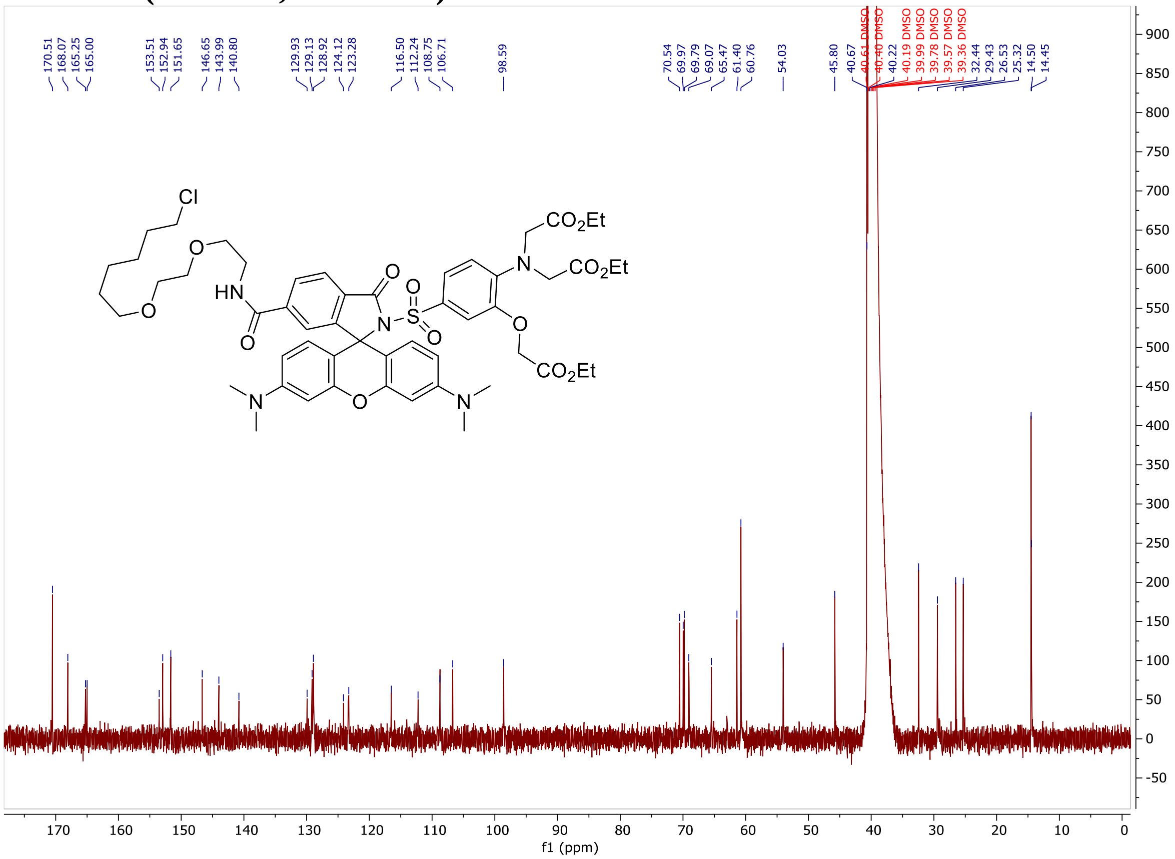

**Dimethyl 2,2'-((((4-((6-((2-(2-((6-chlorohexyl)oxy)ethoxy)ethyl)carbamoyl)-3',6'-bis(dimethylamino)-3-oxospiro[isoindoline-1,9'-xanthen]-2-yl)sulfonyl)phenyl)azanediyl)bis(ethane-2,1-diyl))bis(sulfanediyl))diacetate (CA-MaPNi-558)**

**[1]H-NMR (400 MHz, DMSO-d6)**

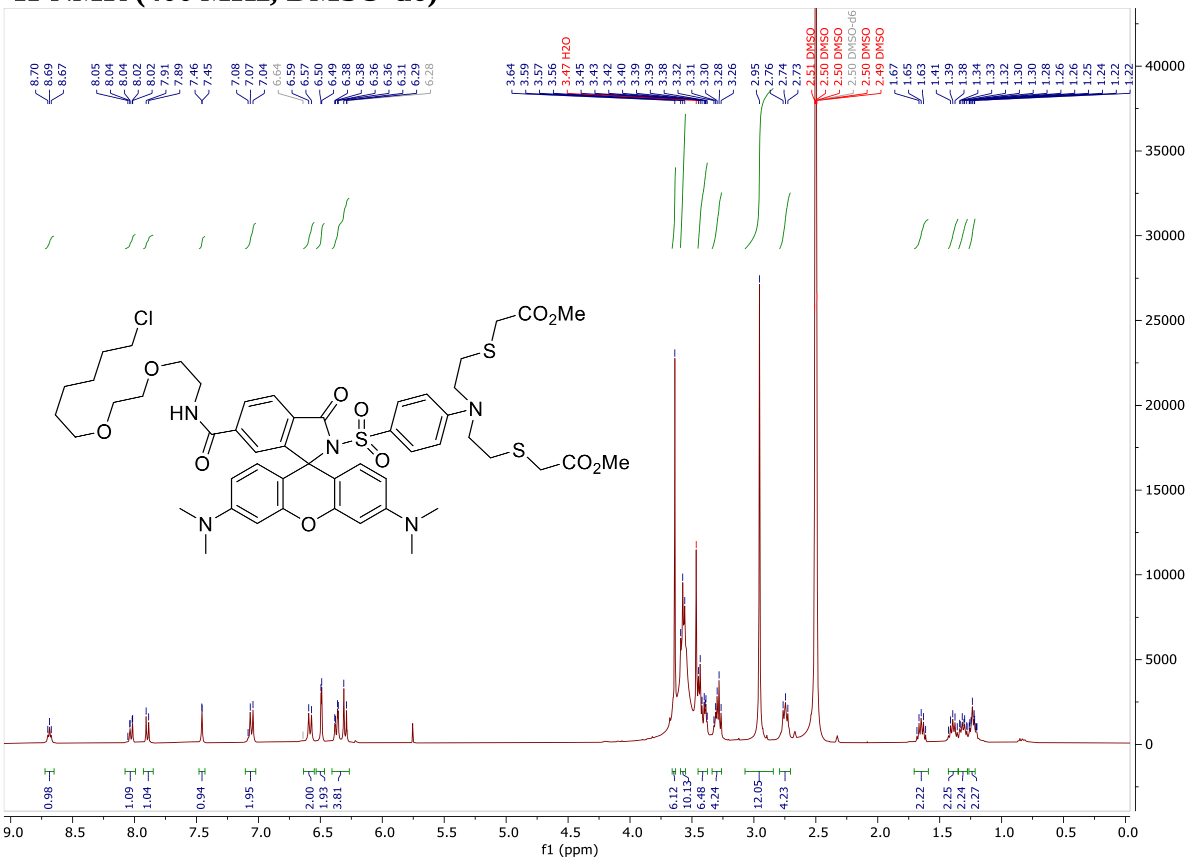

#### $^{13}$C-NMR (101 MHz, DMSO-d6)

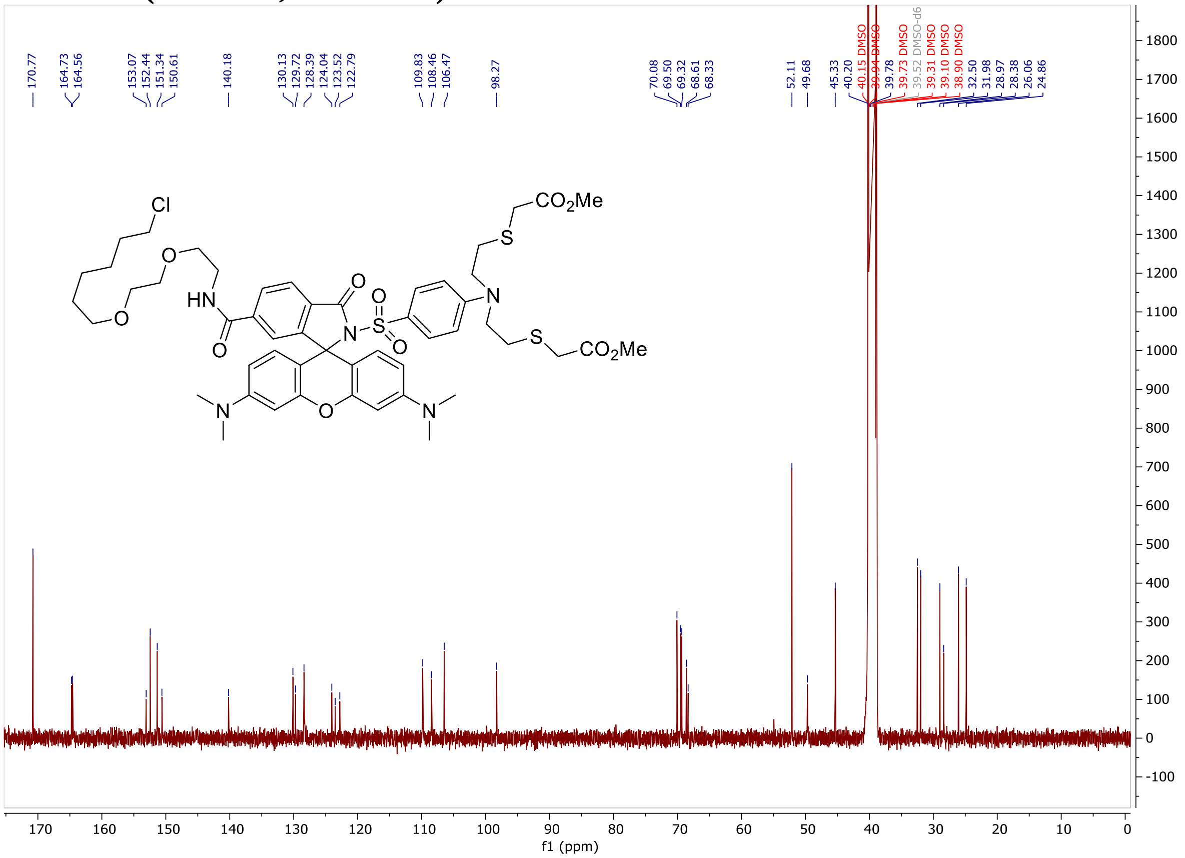

**2-((4-(bis(2-((2-(ethylthio)ethyl)thio)ethyl)amino)phenyl)sulfonyl)-N-(2-(2-((6-chlorohexyl)oxy)ethoxy)ethyl)-3',6'-bis(dimethylamino)-3-oxospiro[isoindoline-1,9'-xanthene]-6-carboxamide (CA-MaPCu-558)**
**$^{1}$H-NMR (400 MHz, DMSO-d6)**

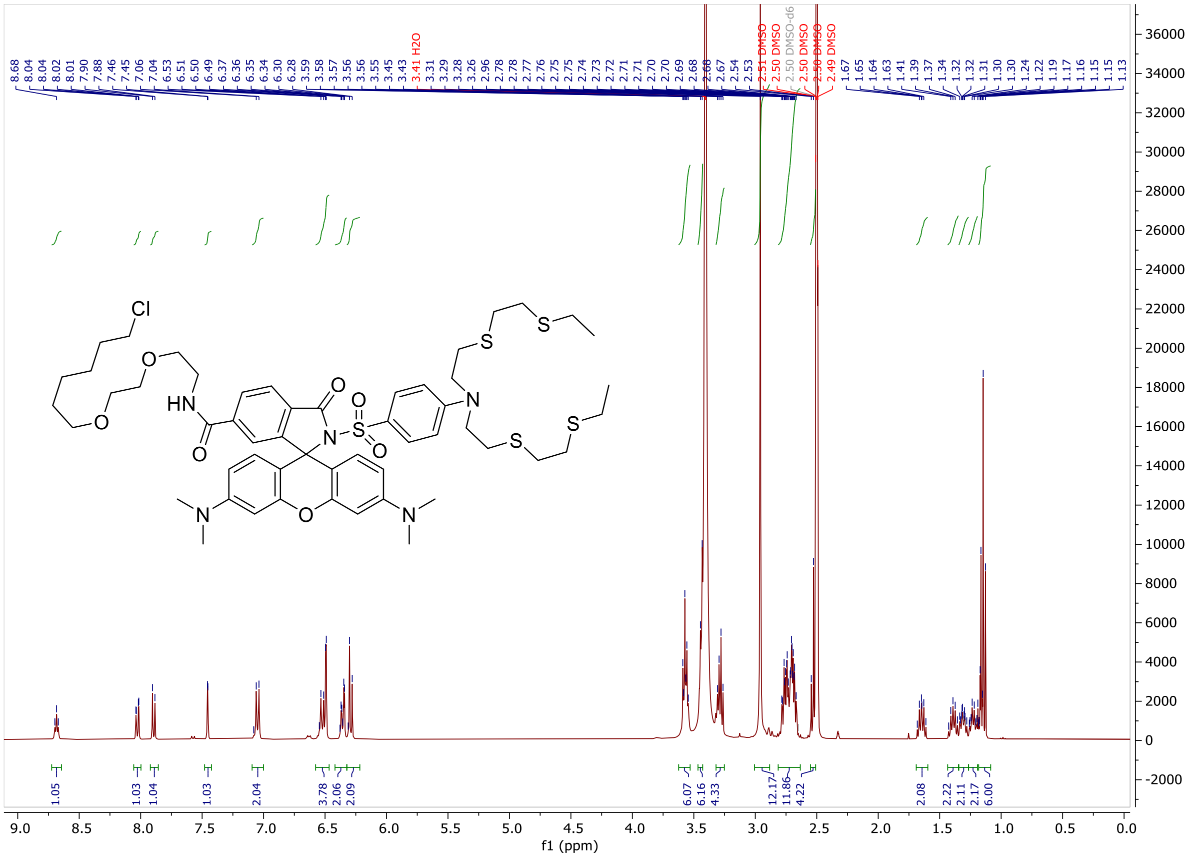

**$^{13}$C-NMR (101 MHz, DMSO-d6)**

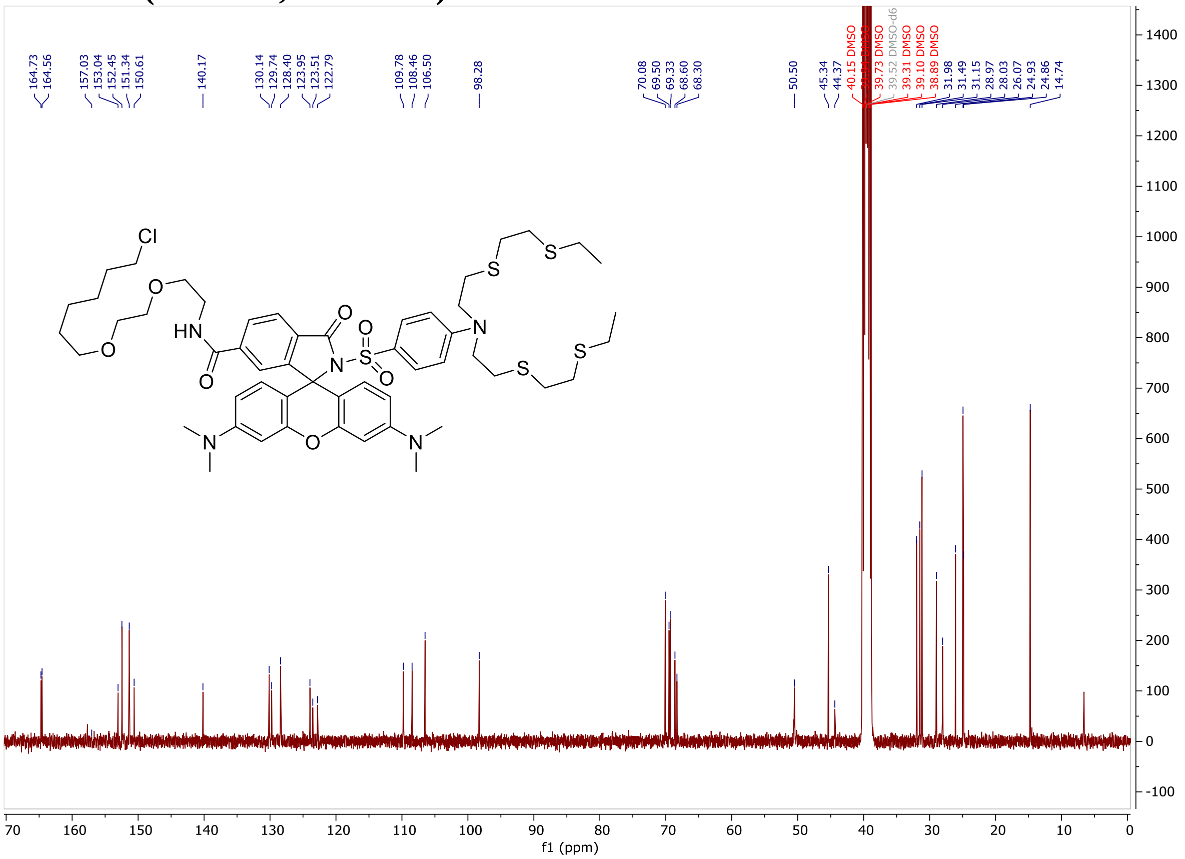

**2'-((4-(1,4,7,10,13-pentaoxa-16-azacyclooctadecan-16-yl)-3-(2-methoxyethoxy)phenyl)sulfonyl)-N-(2-(2-((6-chlorohexyl)oxy)ethoxy)ethyl)-3,6-bis(dimethylamino)-10,10-dimethyl-3'-oxo-10H-spiro[anthracene-9,1'-isoindoline]-6'-carboxamide (CA-MaPK-619)**

**$^1$H-NMR (400 MHz, $CD_3OD$)**

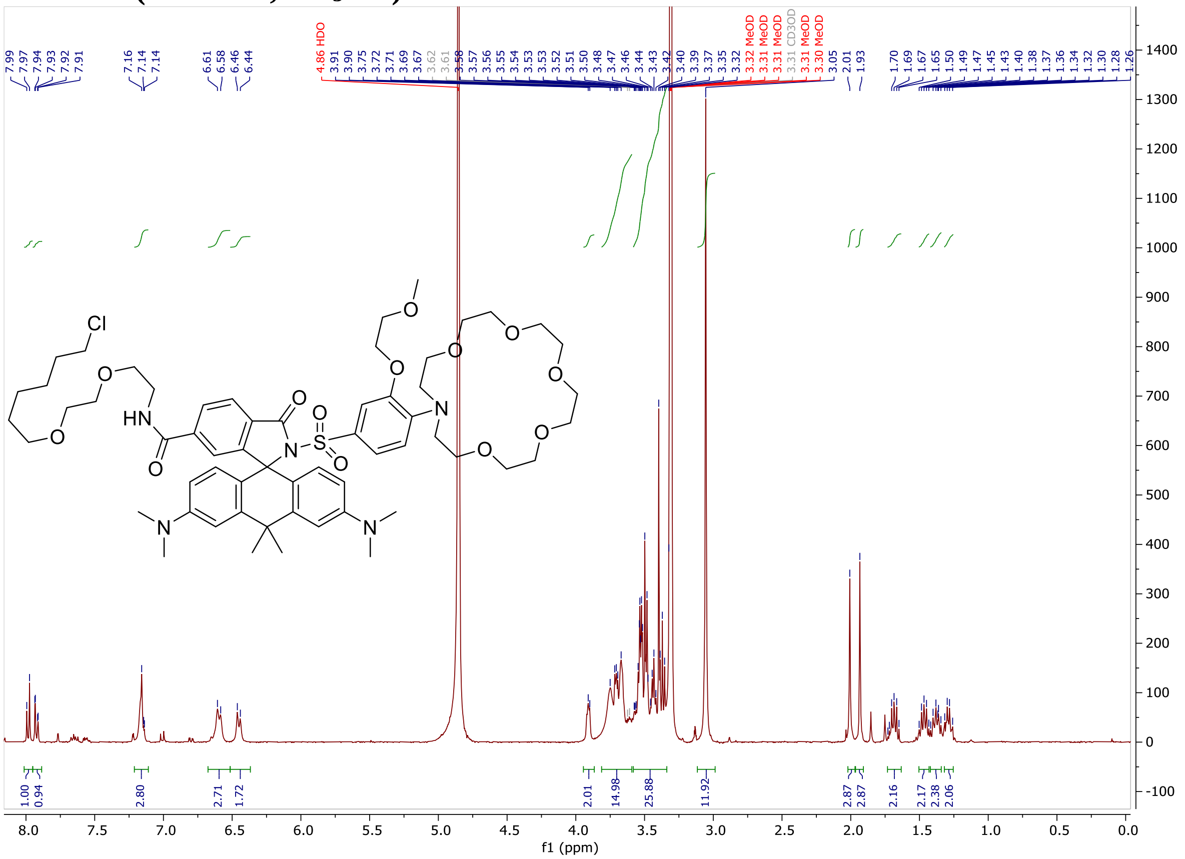

**2'-((4-(1,4,7,10,13-pentaoxa-16-azacyclooctadecan-16-yl)-3-(2-methoxyethoxy)phenyl)sulfonyl)-N-(2-(2-((6-chlorohexyl)oxy)ethoxy)ethyl)-3,7-bis(dimethylamino)-5,5-dimethyl-3'-oxo-5H-spiro[dibenzo[b,e]siline-10,1'-isoindoline]-6'-carboxamide (CA-MaPK-656)**

**$^1$H-NMR (400 MHz, $CD_3OD$)**

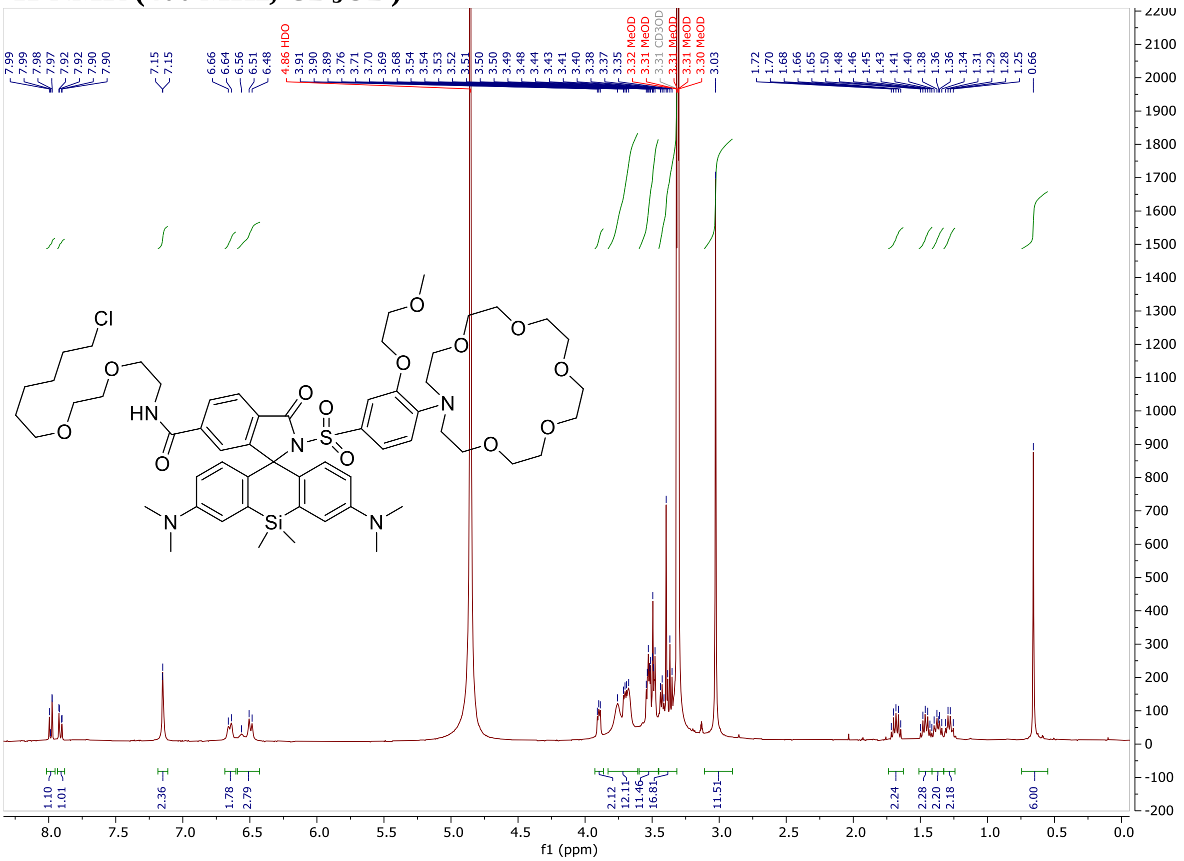

**2'-((4-(1,4,7,10-tetraoxa-13-azacyclopentadecan-13-yl)-3-methoxyphenyl)sulfonyl)-N-(2-(2-((6-chlorohexyl)oxy)ethoxy)ethyl)-3,6-bis(dimethylamino)-10,10-dimethyl-3'-oxo-10H-spiro[anthracene-9,1'-isoindoline]-6'-carboxamide (CA-MaPNa-619)**

**$^{1}$H-NMR (400 MHz, $CD_3OD$)**

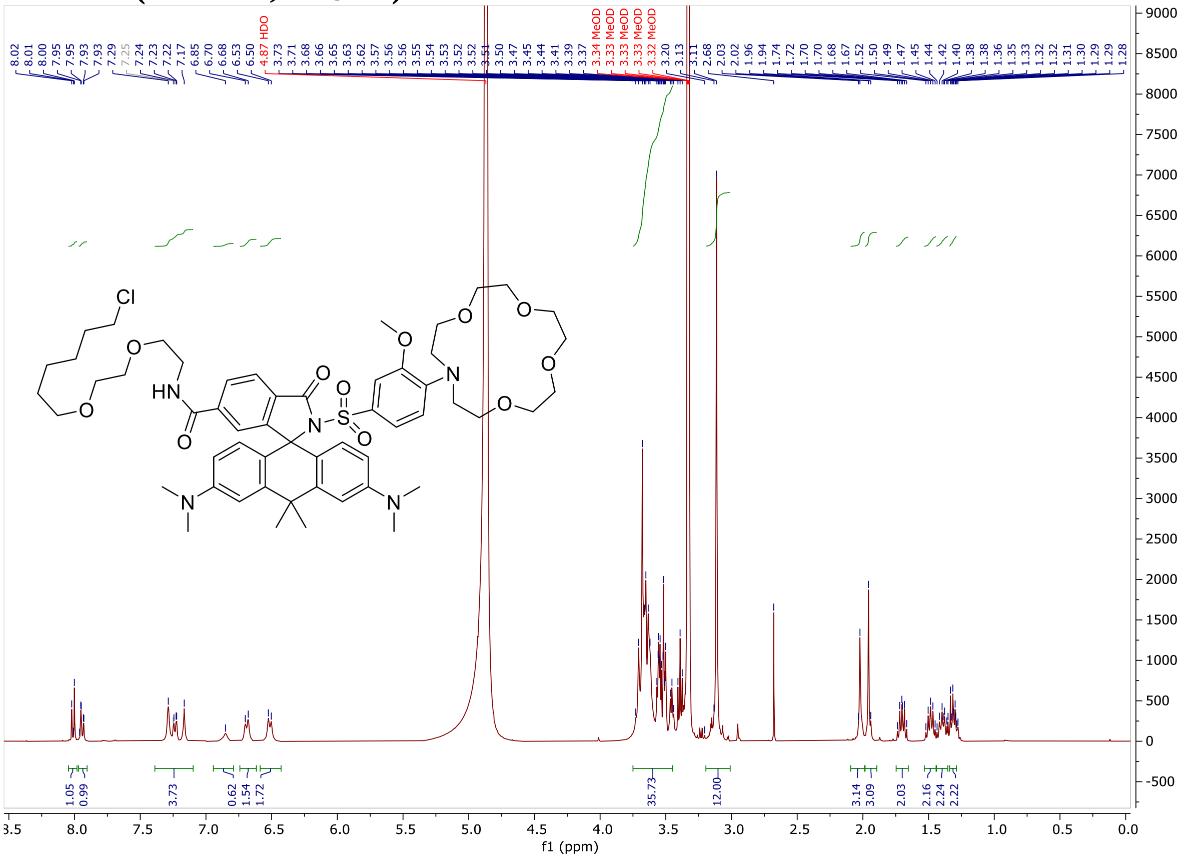

**2'-((4-(1,4,7,10-Tetraoxa-13-azacyclopentadecan-13-yl)-3-methoxyphenyl)sulfonyl)-N-(2-(2-((6-chlorohexyl)oxy)ethoxy)ethyl)-3,7-bis(dimethylamino)-5,5-dimethyl-3'-oxo-5H-spiro[dibenzo[b,e]siline-10,1'-isoindoline]-6'-carboxamide (CA-MaPNa-656)**

**$^{1}$H-NMR (400 MHz, $CD_3OD$)**

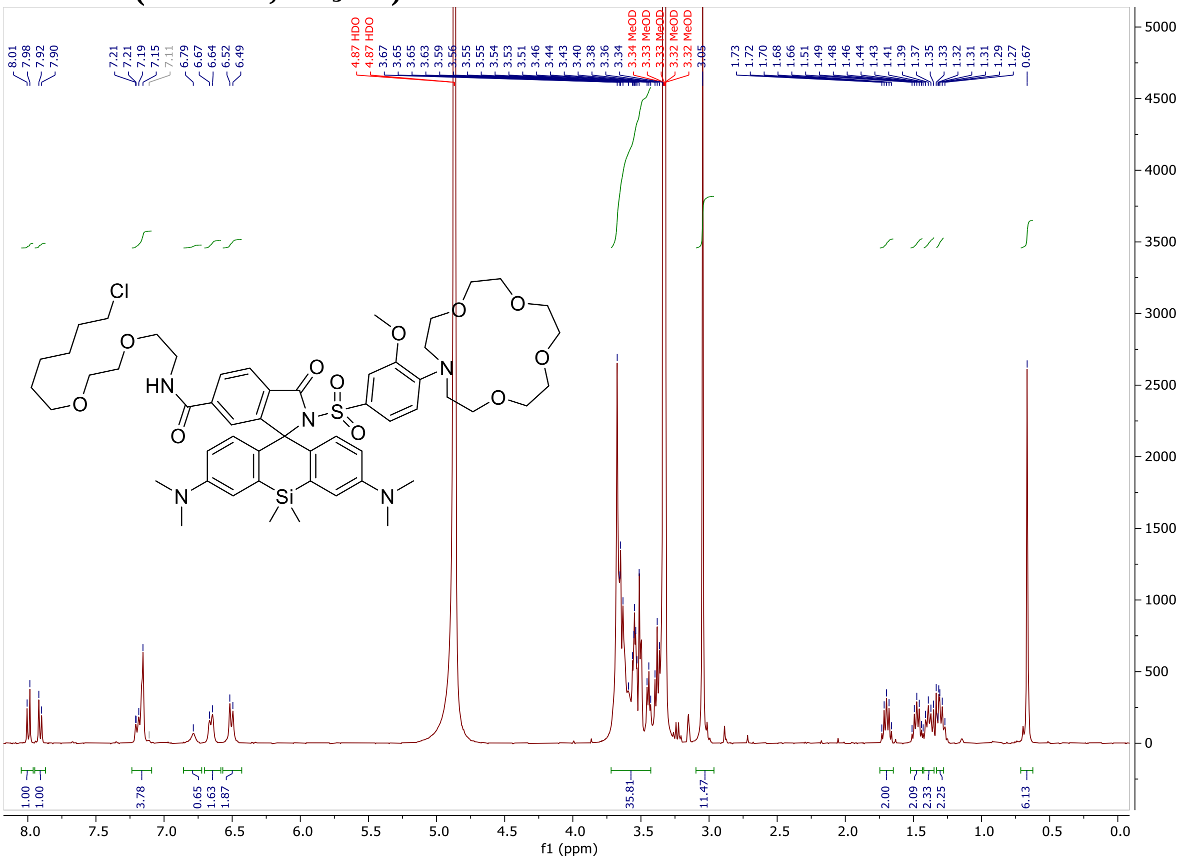

**2-((4-(1,4,7,10,13-pentaoxa-16-azacyclooctadecan-16-yl)-3-(2-methoxyethoxy)phenyl)sulfonyl)-N-(4-(((2-amino-6-(trifluoromethyl)pyrimidin-4-yl)oxy)methyl)-2-fluorobenzyl)-3',6'-bis(dimethylamino)-3-oxospiro[isoindoline-1,9'-xanthene]-6-carboxamide (TF-MaPK-558)**

**$^{1}$H-NMR (400 MHz, $CD_3OD$)**

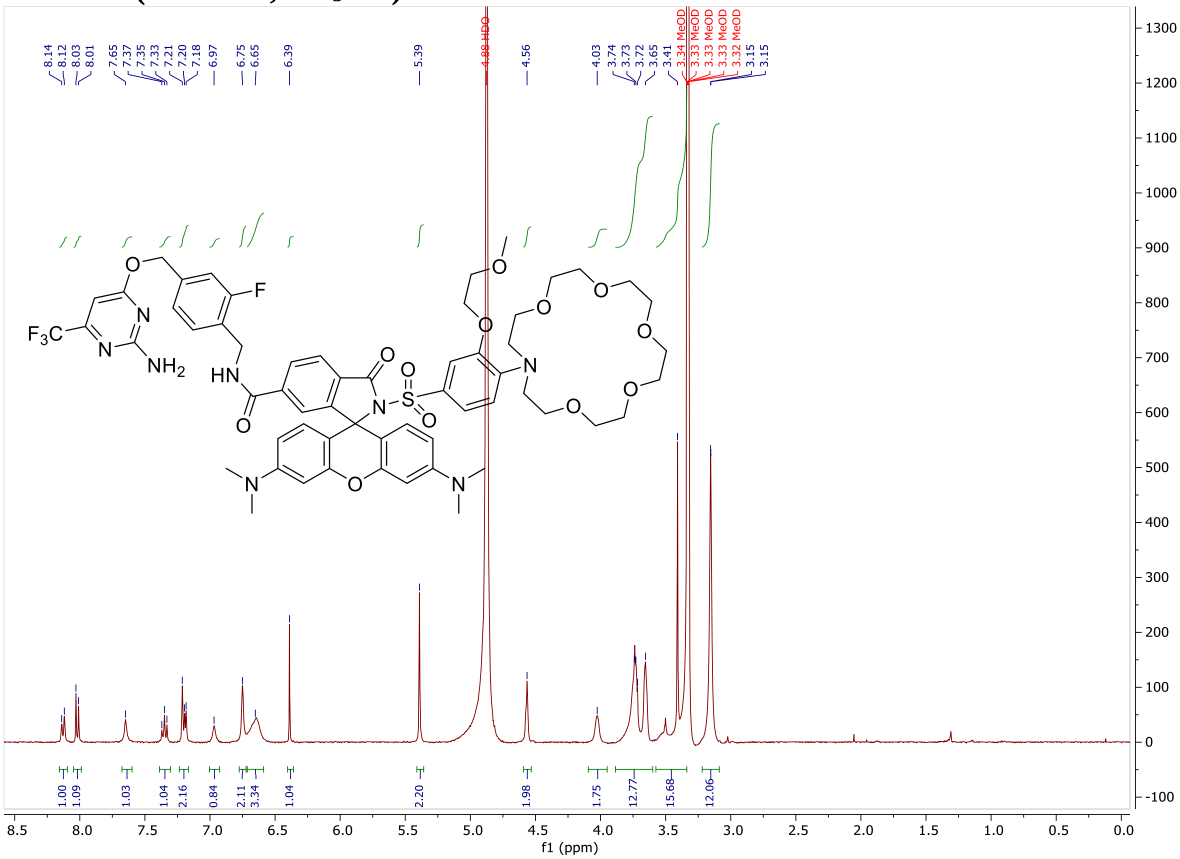

**2-((4-(1,4,7,10-tetraoxa-13-azacyclopentadecan-13-yl)-3-methoxyphenyl)sulfonyl)-N-(4-(((2-amino-6-(trifluoromethyl)pyrimidin-4-yl)oxy)methyl)-2-fluorobenzyl)-3',6'-bis(dimethylamino)-3-oxospiro[isoindoline-1,9'-xanthene]-6-carboxamide (TF-MaPNa-558)**

**$^{1}$H-NMR (400 MHz, $CD_3OD$)**

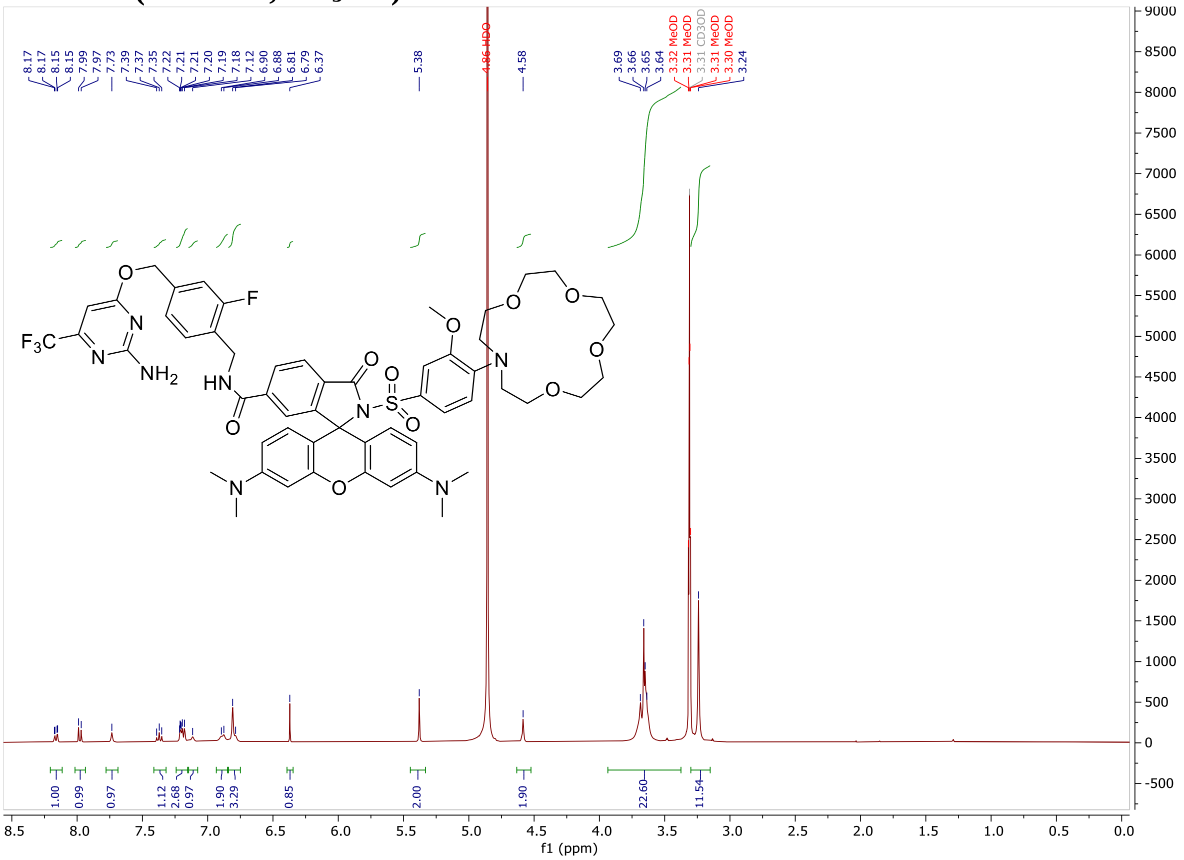
